# Accumulation of virtual tokens towards a jackpot reward enhances performance and value encoding in dorsal anterior cingulate cortex

**DOI:** 10.1101/2025.03.03.640771

**Authors:** Demetrio Ferro, Habiba Azab, Benjamin Hayden, Rubén Moreno-Bote

## Abstract

Normatively, our decisions ought to be made relative to our total wealth, but in practice, we make our decisions relative to variable, decision-time-specific set points. This predilection introduces a major behavior bias that is known as reference-point dependence in Prospect Theory, and that has close links to mental accounting. Here we examined neural activity in the dorsal anterior cingulate cortex (dACC) of macaques performing a token-based risky choice task, in which the acquisition of six tokens (accumulated over several trials) resulted in a jackpot reward. We find that subjects make faster and more accurate choices, and that they are less prone to risk-taking as offer contingencies are easier and the jackpot reward becomes more likely to be achieved. By comparing alternative models that accounted for progressive token accumulation, we found that subjective evaluations are best explained by a reference-dependent value ‘RDV’ model where offer values are considered as potential gains or losses with respect to a token-dependent reference. The reference-dependent model allows to implement a dynamical comparison of the two offered values to each other and to the number of missing tokens to reach the six tokens threshold as jackpot approached. In dACC, we find that gains in subjective values entail higher fractions of encoding cells than losses, and that the encoding tuning of expected utility variables is best aligned to choices in gains than in losses. These results suggest a neural basis of reference dependence biases in shaping decision-making behavior and highlight the critical role of value representations in dACC in driving evaluations.

## Introduction

In their seminal work, Kahneman and Tversky^1^ fundamentally reshaped our understanding of decision-making under risk. While utility and probability distortion components of Prospect Theory are important, the assessment of context-dependent references used in value-based decisions is equally critical. Yet, whereas the neural basis of risk and probability distortion is well understood, the neural processes underlying reference dependence remain less clearly delineated. In animal studies, reference dependence can be difficult to study on single trials, but it becomes tractable when considering behavioral variability across decision trials. Previous studies have examined the influence of behavioral history on decision-making^2–5^, typically using paradigms in which subjects choose between probabilistic options with immediate reward delivery. However, adopting token-based economic tasks allows to investigate the progressive effects of delayed rewards on choice and its neural representations.

Token-based decision-making tasks provide a powerful framework for studying the progressive effects of delayed rewards on subjective value (SV), influencing risk preferences and decision strategies^15^. In these paradigms, choices deliver virtual tokens rather than immediate rewards, contributing to a large jackpot reward once a predetermined token count is reached. Token accumulation naturally introduces shifting internal evaluation of reference-dependent gains or losses inspired by Prospect Theory.

Advances in the field of neurophysiology have substantially contributed to our understanding of neural mechanisms functionally involved in decision-making, which include brain regions associated with reward processing such as the orbitofrontal cortex (OFC), the ventromedial prefrontal cortex (vmPFC), the anterior cingulate cortex (ACC), and the ventral striatum^6–14^. The vmPFC and OFC are predominantly associated with the computation of relative offer value^15–21^ and have been functionally linked to cumulative reward signals in token-based tasks^22^, whereas the ACC plays a role in evaluating potential outcomes, monitoring action costs, and tracking motivational signals^23–33^. Importantly, the dorsal ACC (dACC) has been implicated in reward anticipation^34,35^, cognitive effort computation^36,37^, and in the integration of delayed or cumulative rewards across various decision-making contexts^29,38–42^. Neural signals in ACC have been previously associated with multi-trial^43^, post-decisional variables^42,44^ and virtual reward expectation^45^, suggesting that this region could support reference-dependent value encoding and behavioral adjustments^46,47^.

In this work we analyzed behavioral and neural data from two macaques performing a token-based decision-making task (Fig. 1A)^21,38–41^. On each trial, subjects chose between two probabilistic offers associated with token gains or losses (ranging from −2 to +3 tokens). Tokens accumulated across trials, and upon reaching a total of six, a large jackpot reward was delivered. We tested whether decision-making strategies and neural encoding in dACC reflected value-based signals alone or if they were shaped by a reference-dependent utility defined by token accumulation.

**Figure 1.**
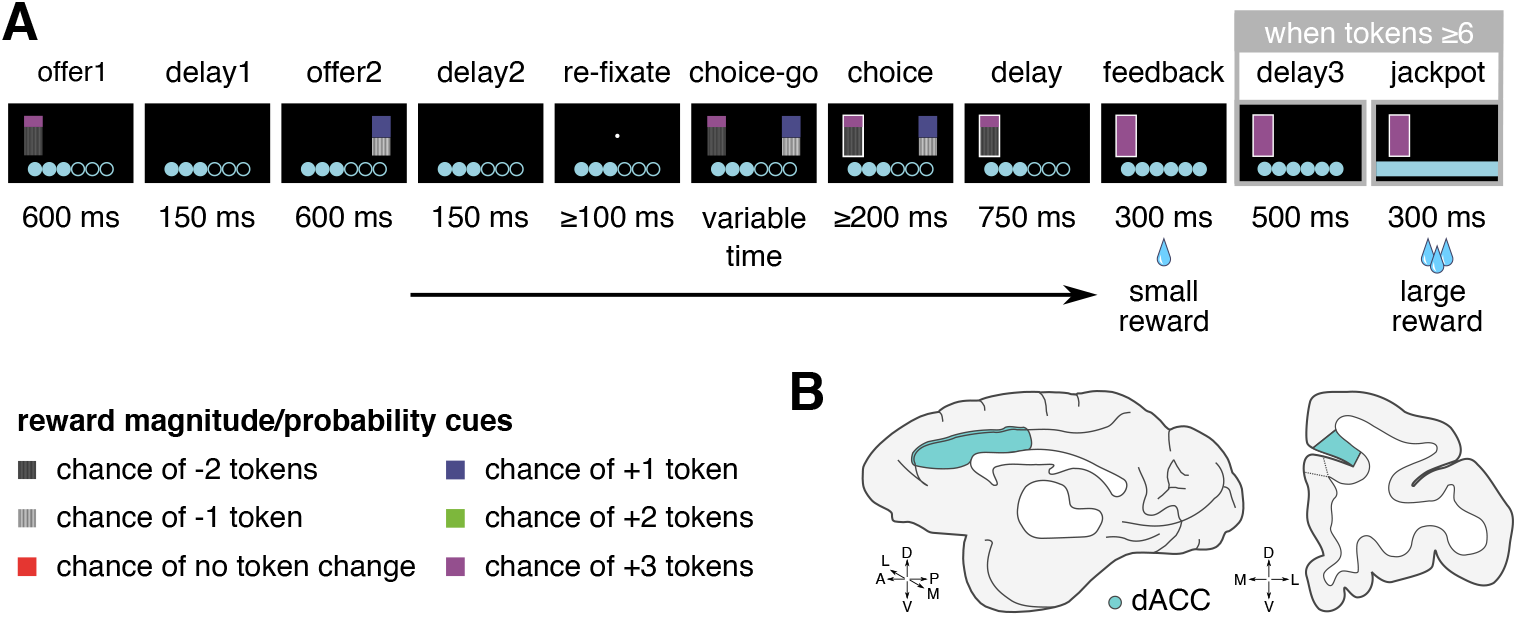
Behavioral task and recording sites. **A.** Two offers are sequentially presented (*offer 1-2*, 600 ms), interleaved by blank screen delays (*delay 1-2*, 150 ms). After the offers presentation, subjects are instructed to re-acquire fixation to a central cross (*re-fixate*) for at least 100 ms and report their choice after a *choice-go* cue, consisting of the presentation of both previous offer stimuli. Choice is reported via target fixation for at least 200 ms (*choice*). The outcome of the chosen probabilistic offer is drawn from uniform distributions, providing (positive or negative) virtual tokens upon risky choice resolution. A small fluid reward (100 µL) is provided in all trials. The token count is visible through execution time as initially unfilled circles at the bottom of the screen, filled by tokens as they are collected. At 6 tokens count, subjects receive a “jackpot”, large reward (300 µL), and the count is reset. The height of the bar stimuli is informative of the associated reward probability, while the color is informative of the magnitude. The probability of the outcomes color-cued by the top and bottom parts of the stimuli are complementary. The probabilities are discretized as 10%, 30%, 50%, 70% and 90%, the magnitudes consist of reward counts (−2, −1, 0, +1, +2, +3) and could include negative values. Offers could also include safe options where 0 (red) or 1 (blue) token are achieved with 100% probability. **B**. Illustration of dACC, brain area targeted for neural data recordings.

At the behavioral level, we found that choices were strongly influenced by the expected value (*EV*) and risk (*R*) of the two offers, but also by jackpots on previous trials (*JPT*), and by accumulated tokens count (*ATC*), which modulated accuracy, reaction time, and risk propensity (Figures 2-4). As subjects approached the jackpot threshold, choices became more accurate and faster, and risk-seeking declined. To formalize these effects, we compared three models of subjective value (SV): two logistic models that related *EV* and *R* (with or without *ATC* interactions) to choice, and a reference-dependent value (RDV) model that treated *ATC* as a dynamic reference point separating gains and losses in a logistic model of *SV* and the choice. The RDV model followed the intuition that subjects dynamically switched the way the evaluated offered values as potential gains or losses relative to the number of missing tokens to jackpot (6 − *ATC*) whenever jackpot was achievable in current trial, or relative to zero whenever jackpot was not achievable. Comparing the three models, we found converging evidence that the RDV model provides the best fit across all behavioral metrics, supporting the hypothesis that token accumulation induces reference-dependent valuation.

**Figure 2.**
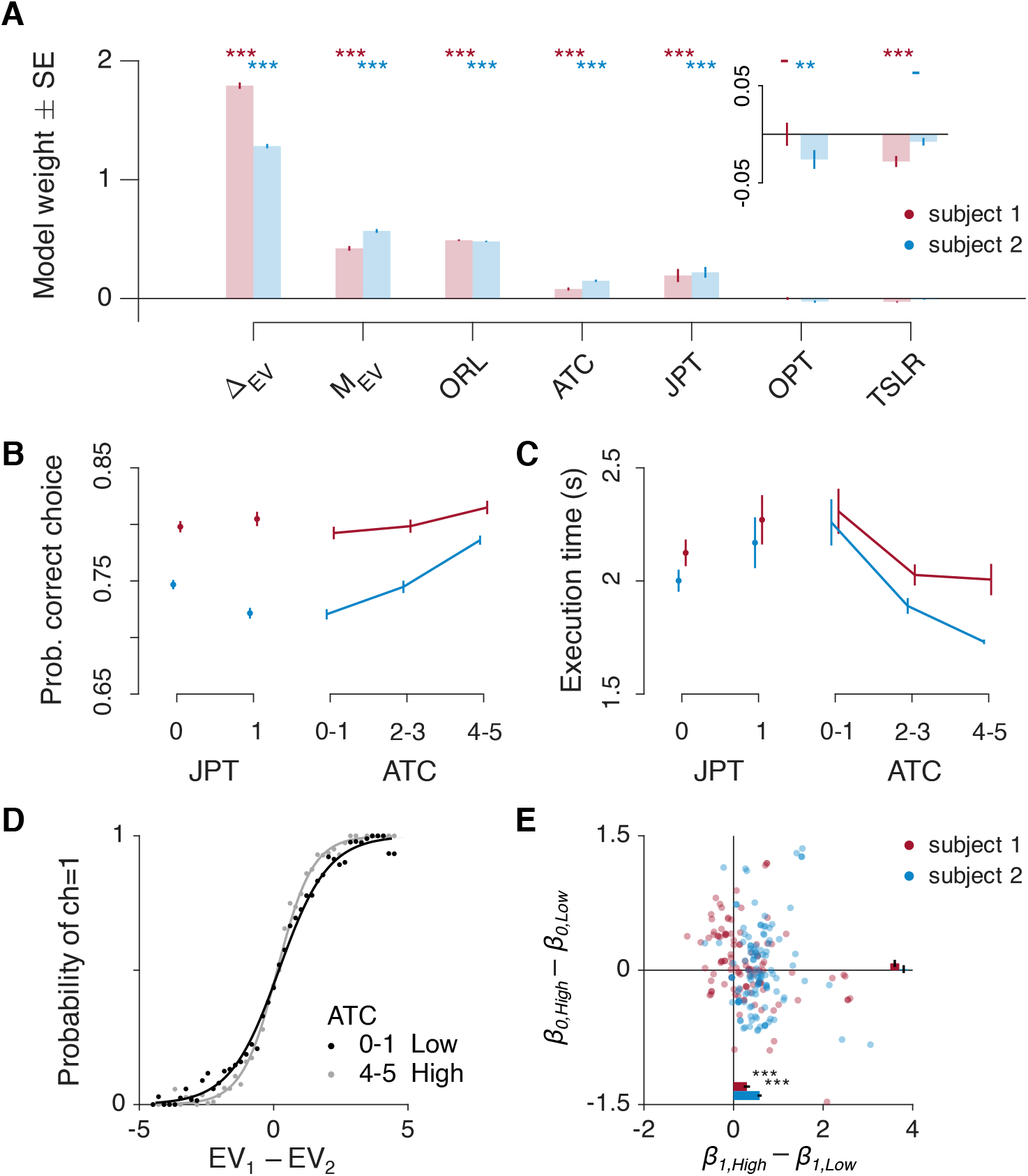
Value-based behavioral variables and accumulated tokens count influence decision-making performances. **A.** Logistic model of the correct choice (Methods 2.1), i.e., choice for the offer with the best expected value (*EV*). We use Δ_*EV*_ = |*EV*_1_ − *EV*_2_| to define the discriminability of best *EV*, inversely related to task difficulty; *M*_*EV*_ = (*EV*_1_ + *EV*_2_)/2 the mean offer *EV*; the offer risk level (*ORL*) the difference between the risk of the offer with highest *EV* and the risk of the offer with lowest *EV*; the accumulated tokens count (*ATC*) as of the the start of current trial; the presence of a jackpot on previous trial (*JPT* = 1 jackpot, *JPT* = 0 no jackpot); the number of trials since last jackpot reward (*TSLR*); and the outcome of previous trial (*OPT*), i.e. the number of tokens resulting from the risky choice made on previous trial. The analysis is applied separately for the two subjects (red, subject 1, *n* = 47030 trials; blue, subject 2, *n* = 58911, Supp. Table ST1). The intercept term *β*_0_ = −0.18 ± 0.04 in subject 1, and −0.46 ± 0.04 in subject 2, *p* < 0.001 in both subjects. Inserts on the top right of the panel show zoomed results for the logistic weights of OPT and TSLR. Numerical model weight estimates ± SE and numerical p-values are reported in Supp. Table ST3. Model coefficients were estimated using maximum likelihood, and significance was assessed via two-sided Wald tests based on the standard errors of the estimated coefficients. **B**. Probability of correct choice (± s.e.m.) vs accumulated reward at the start of each trial (Low: *ATC* = [0, 1]; Medium: *ATC* = [2, 3]; High: *ATC* = [4, 5]). Sample size for each condition (number of trials pooled across sessions) is detailed in Supp. Table ST1. Numeric means ± s.e.m. are reported in Supp. Table ST4. **C)** Same as B but showing the task execution time. Numeric means ± s.e.m. are reported in Supp. Table ST4. **D)** Logistic regression of first offer choice (ch1), designed as logit(*ch*1) = *β*_0_ + *β*_1_(*EV*_1_ − *EV*_2_) for Low (*ATC* = [0, 1]) and High (*ATC* = [4, 5]) accumulated tokens count (data combined for the two subjects, High: β_0_ = −0.2, *p* = 3.6 ⋅ 10^−31^, β_1_ = 1.52, *p* < 10^−308^, *n* = 24099, Low: β_0_ = −0.21, *p* = 5.6 ⋅ 10^−82^, β_1_ = 1.09, *p* < 10^−308^, *n* = 47563; Methods 2.2; Supp. Table ST1). Model coefficients were estimated using maximum likelihood, and significance was assessed via two-sided Wald tests based on the standard errors of the estimated coefficients. **E)** Difference in (*β*_0_, *β*_1_) weights in D for High minus Low accumulated tokens count. Dots are weights differences in each session (*n* = 109 sessions, red for subject 1, *n* = 118, blue for subject 2). Bars show mean ± s.e.m. across sessions, significance is assessed via two-tailed Wilcoxon signed-rank tests, *p* = 1.77 ⋅ 10^−3^ for *β*_1_ in subject 1, *p* = 1.34 ⋅ 10^−19^ in subject 2.

Finally, we examined neural correlates of these computations in dACC by regressing the spike rate of each recorded cell to *SV* defined according in the three behavioral models tested. Across models, substantial fractions of neurons encoded the two *SV*s, peaking during offer presentation epochs. Critically, the RDV model revealed a gain-dominant recruitment of cells in the encoding pattern: gains were represented by significantly larger fractions of cells than losses, and the neural tuning of gain-related *EV* encoding predicted choice behavior more accurately than loss-related *EV*, or *R* variables in either gains or losses. These findings indicate that dACC dynamically integrates expected value with *ATC*-dependent reference signals to guide goal-directed behavior.

## Results

The two subjects performed correct choices, i.e., choices that maximized outcomes, as measured by the offer expected value (*EV*), in most trials (subject 1: 79.78 ± 0.19% mean ± s.e.m., subject 2: 74.19 ± 0.18%). By using a generalized model of the choice (Methods 2.1), we found that correct choices, i.e., choices for the offer with best *EV*, were predominantly influenced by the difference in expected values (Δ_*EV*_ = |*EV*_1_ − *EV*_2_|), the average magnitude of the two *EV*s (*M*_*EV*_ = *EV*_1_/2 + *EV*_2_/2), the difference in risk between the offer with the best *EV* and the offer with worst *EV* (the offers risk level, *ORL*), the tokens collected in task trials previous to current trial (accumulated tokens count, *ATC*), and the achievement of a jackpot on previous trial (*JPT*) (Fig. 2A, Supp. Table ST3). These factors had a strong and statistically significant impact on the decisions made by the two subjects, suggesting that both relied on these variables to form heuristics and choose offers with the best *EV*. Here, the influence of *ATC* is particularly critical, as it not only reflects the motivational drive toward jackpot but also provides a progressive context for value computations. The tokens count shifts the decision frame from evaluating offer values relative to each other when jackpot is out of reach to comparing offer values to jackpot threshold when nearing jackpot, a reference-dependent evaluation paradigm that is central to our investigations. Additionally, we find that subject 2 tended to make fewer choices for the offer with the best *EV* for better outcomes on previous trial (*OPT*), and both subjects made fewer best *EV* choices as the number of trials since the last jackpot reward (*TSLR*) increased.

### The role of accumulated tokens in choice performance

We found that the accumulated tokens count (*ATC*) has an important impact on the subject’s choices. When considering choices for the offer with the best offer *EV* (Methods 2.2), we found that subjects were more accurate (Fig. 2B) and faster (Fig. 2C) when they had collected more tokens, i.e. when they got closer to the possibility of achieving a jackpot reward (Supp. Table ST4). Conversely, the subjects took longer to perform the task and were less accurate when the accumulated tokens count was lower. To investigate this aspect further, we modeled the probability of choosing one of the two offers in relation to the difference in expected value of the two offers, by using a logistic model logit(*ch*_1_) = *β*_0_ + *β*_1_(*EV*_1_ − *EV*_2_), and fit the model separately in trials with ‘Low’ (*ATC* = 0 − 1) or ‘High’ (*ATC* = 4 − 5) accumulated tokens count (Fig. 2D). We found a significant increase in the slope of the logistic relationship (β_1,*High*_ − β_1,*Low*_, p= 1.77 ⋅ 10^−4^ in subject 1, p= 1.34 ⋅ 10^−19^ in subject 2, two-tailed Wilcoxon signed rank test for zero median), corroborating previous insights about higher task engagement for higher accumulated tokens count (Fig. 2E).

### Jackpot, accumulated tokens count, and best offer discriminability improve choice performances

We computed the fraction of errors in choosing the offer with the best *EV* based on whether the current trial was preceded or not by a jackpot reward, for different ranges of accumulated tokens count, and for different levels of difficulty in discriminating between the best and the worst offer. For accumulated tokens count, we used the ranges ‘Low’ (*ATC* = [0, 1]), ‘Medium’ (*ATC* = [2, 3]) and ‘High’ (*ATC* = [4, 5]) (Supp. Table ST1). For the difficulty, we used Δ_*EV*_ and the respective median(*Δ*_*EV*_) = 1 as difficulty discriminant, leading to split data in ‘Hard’ and ‘Easy’ trials, respectively based on whether Δ_*EV*_ < 1 or Δ_*EV*_ ≥ 1 (Supp. Table ST2). Combining the jackpot conditions, accumulated tokens count, and difficulty, we find that the fraction of errors in choosing the offer with best *EV* most differentiate based on the trial difficulty (Fig. 3, Supp. Table ST5). The presence of jackpot before the current trial did not have a significant impact on the fraction of errors of subject 1 (Easy trials: No Jackpot versus Jackpot, p= 0.067, two-tailed Wilcoxon signed-rank test; Hard trials: p= 0.24), while in subject 2 the fraction of errors significantly increased after jackpot only in easy trials (Easy trials:, p= 1.21 ⋅ 10^−10^; Hard trials: p= 0.31, Fig. 3A, Supp. Table ST5).

**Figure 3.**
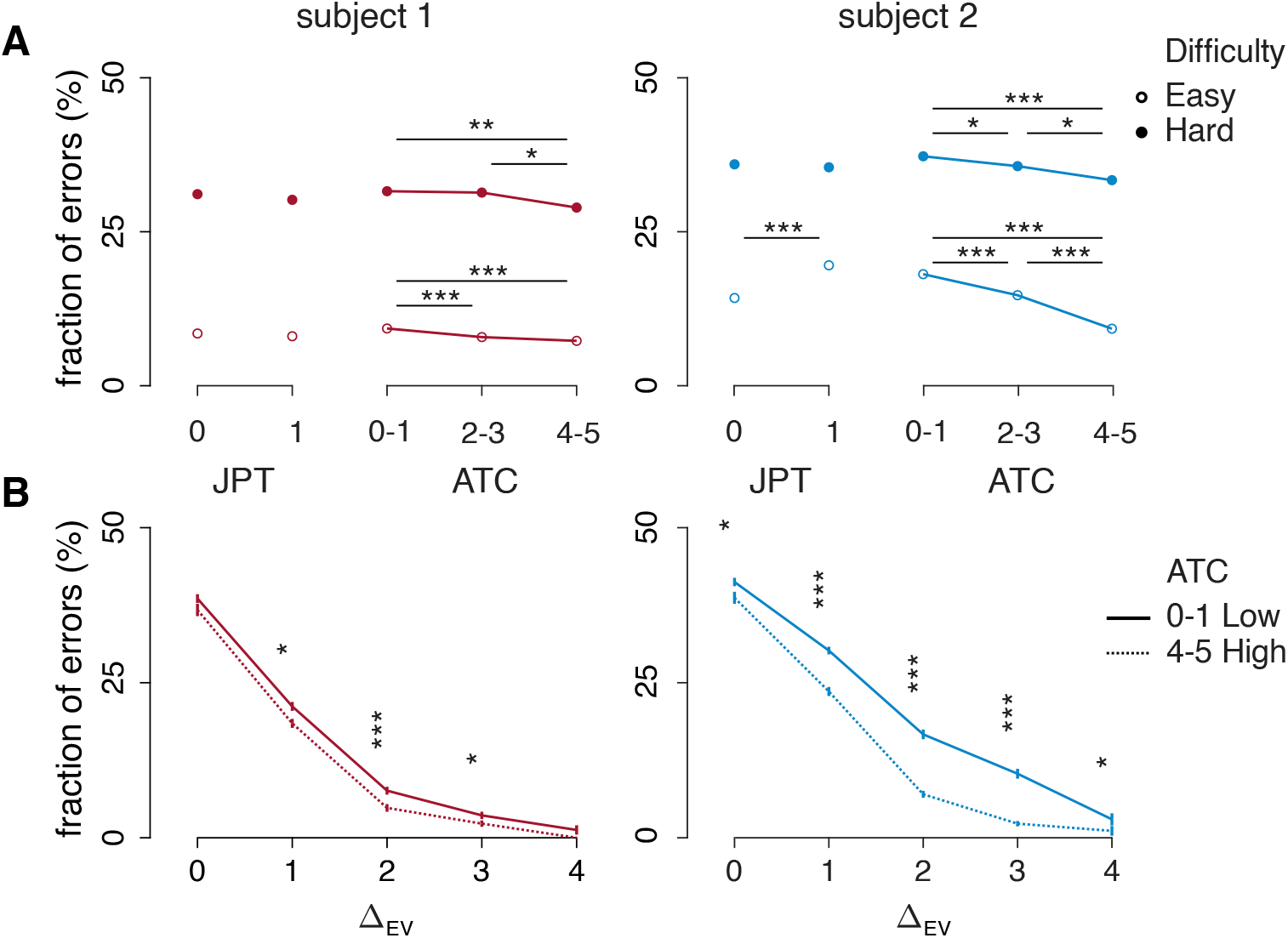
Fraction of errors in reporting correct choice decreases with accumulated tokens count and task difficulty. **A)** Average fraction of errors (subject 1, n=109 sessions; subject 2, n=118) for cases of no jackpot (*JPT* = 0) or jackpot (*JPT* = 1) on previous trial and for different ranges of accumulated tokens count as of the start of current trial (Low: *ATC* = [0, 1]; Medium: *ATC* = [2, 3]; High: *ATC* = [4, 5]). Sample size for each condition (number of trials pooled across sessions) is detailed in Supp. Table ST1. Data are split based on difficulty as ‘Hard’ (filled markers, Δ_*EV*_ < median(Δ_*EV*_)) and ‘Easy’ (empty markers, Δ_*EV*_ ≥ median(Δ_*EV*_)). Sample size for each condition is detailed in Supp. Table ST2. Left: subject 1, right: subject 2. Differences in *ATC* ranges are tested via two-sided Wilcoxon signed-rank tests, FDR corrected via Benjamini-Hochberg procedure (*p< 0.05, **p< 0.01, ***p< 0.001). **B)** Average fraction of errors ± s.e.m. (subject 1, n=109 sessions; subject 2, n=118) for binned values of difficulty, inversely related to Δ_*EV*_ = |*EV*_1_ − *EV*_2_|. Sample size for each condition is detailed in Supp. Table ST2. Data are split in ‘Low’ (solid, *ATC* = [0, 1]) and ‘High’ (dotted, *ATC* = [4, 5]) accumulated tokens count. Left: subject 1, right: subject 2. Differences in bins are assessed via two-tailed Wilcoxon rank sum (*p< 0.05, **p< 0.01, ***p< 0.001). Numeric means ± s.e.m. and p-values for all comparisons are reported in Supp. Table ST5.

The accumulated tokens count had a significant impact on the fraction of errors of both subjects, showing a decreasing trend in the fraction of errors in choosing the offer with the best *EV* as the *ATC* increased (Fig. 3A, Supp. Table ST5). Note that the task trial occurrences become fewer and fewer as *ATC* values grow from Low to High, regardless of the task difficulty (Supp. Table ST1). Since the most prominent difference in fractions of errors was based on task difficulty, we also investigated fractions of errors for binned values of Δ_*EV*_, which we used as an inverse metric of difficulty. As expected, the fraction of errors decreased with Δ_*EV*_, as larger Δ_*EV*_ indicates easier detection of the best offer. By comparing the fractions of errors in High and Low accumulated tokens counts, we found a lower fraction of errors in High accumulated tokens, indicating that accumulating a larger number of tokens boosts the best *EV* discrimination, most prominent and significant in both subjects for intermediate difficulty (Δ_*EV*_=[1, 3] subject 1, significant for all Δ_*EV*_ in subject 2, Fig. 3B). In similar way as for *ATC*, we find fewer trial occurrences when considering larger Δ_*EV*_ bins, coinciding with easier trials (Supp. Table ST2). These results align with the observation that subjects tend to be more accurate and faster whenever they accumulate more tokens before the current trial, adding a further level of stratification in considering the difficulty, that has a prominent role in committing best offer contingency detection errors.

### Jackpot, accumulated tokens count, and difficulty in best offer discriminability impact risk-taking propension

We asked whether previous results suggesting improvement in choice speed and accuracy with *ATC* could relate to a higher propensity for risky options whenever subjects felt that achieving a jackpot reward was less likely and far apart, requiring further accumulation of tokens. By comparing the fraction of trials where subjects chose the offer with the highest risk, we observed that the presence of a jackpot on the previous trial decreased the fraction of choices for the riskiest option, significant for subject 1 (subject 1: p= 1.04 ⋅ 10^−3^, two-tailed Wilcoxon signed-rank test, subject 2: p= 0.37, Supp. Table ST6). We found that both subjects significantly made more risky choices when choosing the offer with best *EV* was hard (Δ_*EV*_ < 1) as opposed to easy (Δ_*EV*_ ≥ 1) trials (subject 1: p= 1.47 ⋅ 10^−19^, two-tailed Wilcoxon signed-rank test, subject 2: p= 4.21 ⋅ 10^−21^, Supp. Table ST6).

By combining difficulty and jackpot conditions, we found that subjects consistently chose risky offers less frequently after a jackpot reward in Hard trials (Fig.4A, Supp. Table ST6), while in Easy trials risky choices after jackpot were less frequent in subject 1 (Fig. 4A, Supp. Table ST6), and more frequent in subject 2 (Fig. 4A, Supp. Table ST6). We reported a remarked difference in the fraction of choices for the riskiest option between Easy and Hard trials, with Hard trials involving higher fractions of choices for the riskiest option across *ATC* conditions (Fig. 4A, Supp. Table ST6). We found a consistent inverted U-shape relationship between the fraction of risky choices and *ATC* ranges in Hard trials in both subjects, showing increasing trend for *ATC* = [0, 2], and decreasing trend for *ATC* = [3, 5] (Fig. 4A; Supp. Table ST6). To further investigate the effect of risk on the decisions of the two subjects, we quantified the risk attitude of subjects by using a Markowitz model, i.e., computing the ratio between the weight of the risk (*β*_2_) and *EV* (*β*_1_) differences between the two offers in a logistic model of the choice including these two variables as choice predictors (Methods 2.4). The model parameter *θ* = −*β*_2_/*β*_1_ describes the subjective behavioral attitude as risk aversion (*θ* > 0) or risk-seeking (*θ* < 0). We report a risk-seeking attitude in both subjects, with higher risk aversion (reduced magnitude of *θ*) as the number of accumulated tokens increases, and in Easy trials (Fig. 4B, Supp. Table ST6). Comparing jackpot conditions, we find that *θ* has smaller, negative magnitude for Hard trials following a jackpot reward, consistent in the two subjects (Fig. 4B; Supp. Table ST6), in line with lower risky choice fractions results (Fig. 4A, Supp. Table ST6).

**Figure 4.**
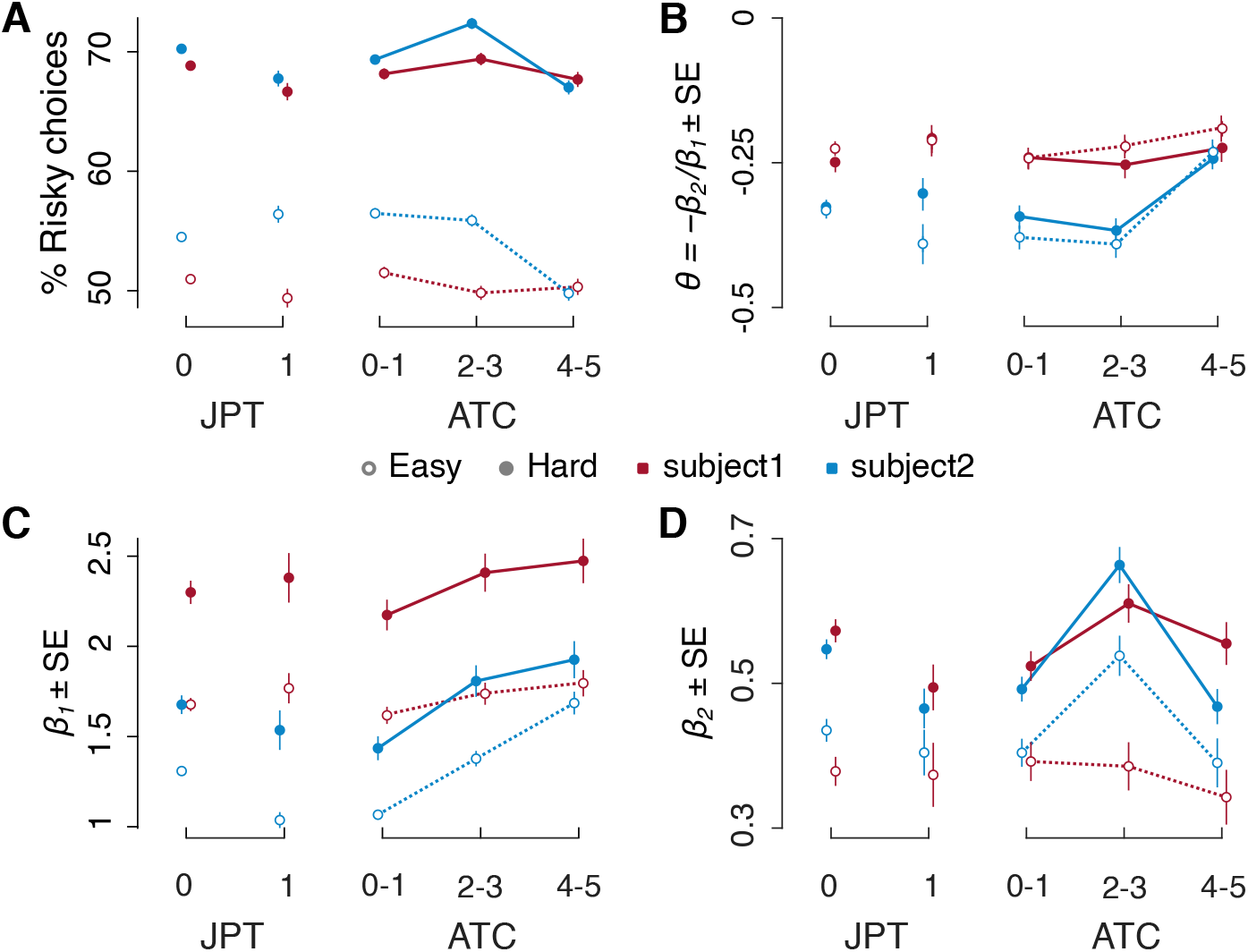
Risk seeking attitude increases with accumulated tokens count and easier best offer discriminability. **A.** Fraction of trials (mean ± s.e.m.) with choice for the offer with higher risk. The data are split in ‘No Jackpot’ (*JPT* = 0), ‘Jackpot’ (*JPT* = 1), and for accumulated tokens count (‘Low’ *ATC* = [0, 1], ‘Medium’ *ATC* = [2, 3], ‘High’ *ATC* = [4, 5]). Data are split for subject 1 (red) and subject 2 (blue), and for Easy (Δ_*EV*_ ≥ 1, empty markers) and Hard (Δ_*EV*_ < 1, filled markers). **B**. Markowitz risk return model for the offer utility based on the mean value (*EV*) and risk (*R*) of the offers. The model parameter (*θ*) describes risk attitude (*θ* < 0 risk seeking, *θ* > 0 risk avoiding) for jackpot cases and for values of accumulated token counts. Data is split in difficulty and across subjects as in A. **C**. Markowitz risk return model, parameter *β*_1_ relative to *EV* weights for jackpot cases and for values of accumulated token counts. Data is split as in A. **D**. Markowitz risk return model, parameter *β*_2_ relative to risk *R* weights for jackpot cases and for values of accumulated token counts. Data is split as in A. **A-D)** Sample size for each condition is detailed in Supp. Table ST1. Numeric values for mean ± s.e.m. or mean ± CI across sessions are reported in Supp. Table ST6.

We found that the *EV* difference has an increasing impact (*β*_1_) on choices for the riskiest option, consistent in the two subjects for Hard trials (Fig. 4C, Supp. Table ST6) than in Easy trials (Fig. 4C, Supp. Table ST6). The risk difference weight (*β*_2_) shows an inversed U-shape, most prominent and consistent in the two subjects for Hard trials (Fig. 4D, Supp. Table ST6), larger than for Easy trials (Fig. 4D, Supp. Table ST6), both results are in line with the fraction of risky choices (Fig. 4A, Supp. Table ST6). The Markowitz variable *θ* = −*β*_2_/*β*_1_, shows decreased magnitude as the accumulated token counts increases for ‘High’ *ATC* ranges, in both Hard (Fig. 4D, Supp. Table ST6), suggesting that when the jackpot reward becomes more likely in tokens count, the subjects are more averse to risk. All the above results found for Low, Medium, High *ATC* ranges are also found for discrete values of *ATC* (Supp. Fig. S1).

### Models of choice: Subjective Values and reference-dependent evaluation in token-based decision-making

To further investigate how subjects integrate behavioral variables and contextual information into their decision, we formalized choice behavior using predictive models of Subjective Value (*SV*). Given the observed effects of *EV, R*, and *ATC* on behavior, we compared three alternative models that differed in how these factors were combined. The first model, a ‘linear model of choice without *ATC* interaction’ defined *SV* as a linear combination of *EV, R* and *ATC*, without interaction terms (Methods 2.4.1). This was extended to a second ‘linear model of choice with *ATC* interaction’ in which *ATC* modulated the influence of *EV* and *R* through interaction terms to capture how the influence of *EV* and *R* varied with *ATC* (Methods 2.4.2). We then introduced a reference-dependent value ‘RDV’ model that incorporates the key idea that token accumulation dynamically defines a reference point for evaluating gains and losses (Methods 2.4.3). In this model, the perceived value of each offer is computed relative to a shifting internal reference *r*(*ATC*), which approaches zero when *ATC* is low and jackpot is out of reach, and transitions to the number of tokens required to reach the jackpot (6 − *ATC*) as jackpot achievement becomes closer. The offer values, instructed by cue colors (*v*, ranging −2 to +3 tokens) was shaped by a token-dependent utility function *u*(*v, ATC*) adapted from Prospect Theory. The curvature of this function depend on *ATC* in two ways: the value sensitivity parameter *γ*(*ATC*) increases linearly with *ATC*, reflecting stronger weighting of *EV* with token accumulation (Fig. 4C, Supp. Fig S1C); the loss-aversion parameter *λ*(*ATC*) varies quadratically with *ATC*, consistent with the inverted U-shaped relationship observed between risk taking and *ATC* (Fig. 4D, Supp. Fig. S1D). This formulation captures how motivational proximity to the jackpot reshapes value computation and risk sensitivity, enabling a smooth reference-dependent transition.

We trained each model using choice, *EV, R* and *ATC* features to estimate regression weights via maximum likelihood and assessed performance using cross-validated test data (*k* = 4 folds). Across all models, *EV* difference emerged as the strongest predictor of choice (Supp. Fig. S2, Supp. Table ST7). In the first model, risk was the second-largest positive weight, followed by a negative constant term and the *ATC* regressor weight, showing relatively low magnitude, statistically significant in subject 1 and in data pooled for the two subjects (Supp. Fig. S2, Supp. Table ST7). In the second model, the *EV* and *ATC* interaction term had second-largest positive weight, followed in magnitude by *R*, a negative constant term, the linear and quadratic interactions between *R* and *ATC* (Supp. Fig. S2, Supp. Table ST7). In the RDV model, we also found meaningful parameter estimates, consistent between subjects. The reference *r*(*ATC*) was defined by two parameters: a threshold *κ*_1_, determining the *ATC* value above which jackpot achievement became behaviorally relevant (subject 1: 3.28±0.05 mean ± s.e.m across sessions, subject 2: 3.09±0.06), and a steepness parameter *κ*_0_ controlling the sharpness of the transition (subject 1: 2.61±0.25, subject 2: 0.97±0.12). The value sensitivity function increased linearly with *ATC* (*γ*(*ATC*) ≈ 0.79 + 0.02*ATC* in subject 1, 0.66 + 0.03*ATC* in subject 2). Since the parameters *γ*(*ATC*) is used as exponent of reference-dependent values, it makes value utility tend to linearity as subjects approach jackpot. The loss-aversion parameter followed an inverted-U profile (*λ*(*ATC*) ≈ 0.49 + 0.18*ATC* − 0.02*ATC*^2^ in subject 1, and ≈ 0.27 + 0.16*ATC* − 0.01*ATC*^2^ in subject 2), aligning with observed shifts in risk preference (Fig. 4D, Supp. Fig. S1D). The RDV model shows strongest regression weight for reference-dependent *EV* terms, followed in magnitude by reference-dependent *R* terms, and by a negative constant term.

Model comparison showed that RDV consistently outperformed the two first models across all metrics, including cross-validated choice prediction accuracy, adjusted coefficient of determination, negative log likelihood, Akaike Information Criteria (AIC), and Bayesian Information Criteria (BIC) (Supp. Fig. S3, Supp. Table S8), providing strong evidence that a reference-dependent, token-based *SV* framework best explains choice behavior in this task. We also assessed that RDV consistently outperformed concurrently adopted models implementing normative temporal discount of delayed reward values^48^ and adaptive biases based on cumulative reward history^22^.

### Neural encoding of Subjective Value: the role of token accumulation and value tuning in behavioral choice readout

We tracked the neural encoding of task-related variables by designing time-resolved spike-rate analyses to detect the fractions of cells (*n* = 129, *n* = 55 in subject 1, *n* = 74 in subject 2) significantly encoding relevant task variables during execution time (Methods 3.1). We designed linear spike-rate models using the same SV definitions introduced for the three choice models assessed. In addition, we tested the possibility to predict behavioral choices based on neural encoding weights by designing Receiver Operating Characteristics (ROC) analyses using regressed spike rate model weights multiplied by test variable data. Lastly, we correlated the Area Under the Curve of ROC analyses to time-averaged spike-rate model weights, investigating how neural tuning properties to task variables also aligned with choice prediction.

The first spike-rate model, which excluded *ATC* interactions, revealed significant fractions of cells significantly encoding the two *SV*s defined as in the ‘linear choice model without *ATC* interactions’ (Methods 2.4.1, 3.1.1). The fractions of significantly encoding cells peaked around the respective offer presentation epochs (Figure 5A-B, Supp. Fig S4). This model also included a regressor for *ATC*, which was associated with notably higher fractions of significantly encoding cells during late task execution epochs, posterior to offer presentation epochs (Figure 5A-B, Supp. Fig S4). We quantified the fraction of cells exclusively or synergistically encoding task variables in task epochs, considering the exclusive encoding of *SV*_1_, *SV*_2_, *ATC*, and their combinations (*SV*_1_&*SV*_2_; *SV*_1_&*ATC*; *SV*_2_&*ATC*; *SV*_1_&*SV*_2_&*ATC*). The significance of fractions of cells were assessed by comparison to the 95^th^ percentile of equivalent fractions from trial-order shuffled data. Since pre-offer 1 and throughout task epochs, cells exclusively encoding *ATC* represented the largest and significant fraction, in line with the task design by which current trial *ATC* is cumulated at the end of previous trial. The fraction of cells encoding *ATC* tended to lower during offer 1 and offer 2 epochs, when *SV* encoding peaked. Following offer 1 onset, a significant proportion of cells encoded *SV*_1_, either exclusively or in combination with *ATC*, while the proportion of cells encoding *SV*_2_, either exclusive or simultaneous to *SV*_1_ and/or *ATC*, was significant after offer 2 onset. Following delay 2, the fractions of cells exclusively encoding the two *SV*s gradually lowered, and *ATC* emerged as the most encoded variable across cells, with minor, though significant fractions of cells showing joint encoding of the two *SV*s and/or *ATC* (Fig. 5C, Supp. Fig. S5).

**Figure 5.**
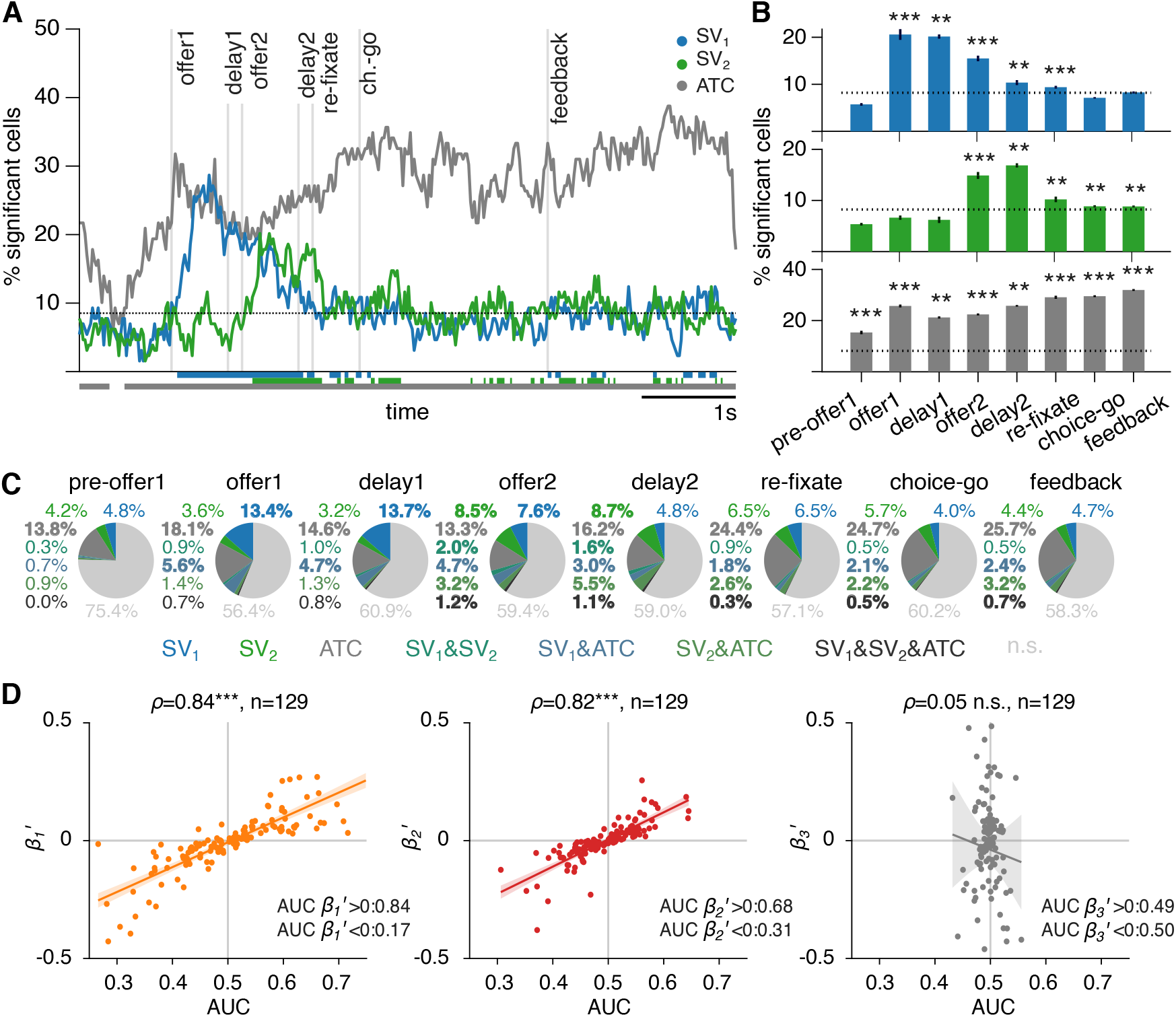
Neural encoding of *SV*s and *ATC* in a spike-rate model without *ATC* interactions and their link to choice prediction. **A.** Solid lines show the fraction of cells significantly encoding *SV*_1_, *SV*_2_ and *ATC* in the ‘linear model without *ATC* interaction’ (Methods 3.1.1). The dotted line represents the 95^th^ percentile threshold from trial-order shuffled data. The percentiles are computed independently and coincide across regressors, reflecting chance level of significance expected under null interaction. Bottom lines indicate time bins where significant fractions exceed the percentile threshold, further assessed in length by cluster-based run length analysis, considering runs whose length is significantly longer than the 95^th^ percentile of equivalent lengths computed within shuffled data. Results are shown for the two subjects separately in Supp. Fig. S4. **B**. Fractions of significant cells for *SV*_1_ (top), *SV*_2_ (middle) and *ATC* (bottom) in task epochs (mean ± s.e.m. across time bins, *n* = 129 total cells). Dotted lines represent the 95^th^ percentile of fractions of significant cells run over trial-order shuffled data, computed separately for each regressor. Significance is assessed via one-tailed signed rank, FDR corrected via Benjamini-Hochberg procedure, testing that empirical data exceed 95^th^ percentile thresholds (\**p* < 0.05, \*\**p* < 0.01, \*\*\**p* < 0.001, Supp. Table ST9). **C**. Time-averaged fractions of *n* = 129 total cells that exclusively encode *SV*_1_, *SV*_2_, *ATC*, or jointly encode (*SV*_1_ and *ATC*), (*SV*_2_ and *ATC*), (*SV*_1_, *SV*_2_ and *ATC*), or are non-significant (n.s.). Results in B represent aggregate categories from C. For example, the fraction of cells encoding *SV*_1_ includes cells exclusively encoding *SV*_1_ as well as cells encoding *SV*_1_ jointly with *SV*_2_ and/or *ATC*. Fractions in bold exceed the 95^th^ percentile of equivalent fractions from trial-order shuffles. Subject-specific results, percentiles and significance are in Supp. Fig. S5. **D**. Correlations between Area Under the Curve (AUC) from ROC analyses of choice and model weights 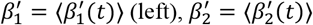 (middle), 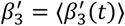 (right) time-averaged at times posterior to the offer 2 onset, for the linear spike-rate model without interactions (Methods 3.2.1). Each dot represents a cell (*n* = 129, subjects combined), with AUC and model weights averaged in time bins posterior to offer 2 onset and cross-validation folds. Pearson’s coefficients *ρ* are assessed in significance via two-tailed *t*-tests (\*\*\**p* < 0.001, *p* = 5.4 · 10^−36^ for β_1_’, *p* = 3.6 · 10^−32^ for β_2_’, *p* = 0.57 for β_3_’). Solid lines show linear regression lines, shaded areas indicate ±CI computed using the standard error of the regression coefficients. The values at the bottom-right of each panel report the median AUC across time bins posterior to offer 2 onset, cells, cross-validation folds for positive (top) and negative (bottom) weights. Results are shown separately for the two subjects in Supp. Fig. S6.

To examine the relationship between neural tuning and behavioral choice, we performed a Receiver Operating Characteristics (ROC) analysis, quantifying the predictive strength of spike-rate regression weights with respect to key decision variables: the *EV* difference between the two offers, the risk difference, and *ATC* (Methods 3.2.1). Predictive performance was assessed using the Area Under the ROC Curve (AUC). Among regressors, the *EV* difference exhibited the strongest choice (median AUC for positive regression weights *β*_1_’ > 0: 0.84, median AUC for negative weights 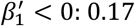, data pooled across the two subjects, Fig. 5D). The risk difference regressor followed (median AUC for 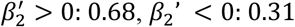, *β*_2_’ < 0: 0.31, Fig. 5D). As we expected, the *ATC* variable itself did not show meaningful choice predictability (median AUC for *β*_3_’ > 0: 0.49, *β*_3_’ < 0: 0.51, Fig. 5D).

Furthermore, spike-rate regression weights for both *EV* and *R* difference showed strong correlations with the respective AUC values (*ρ* = 0.84 for *EV* difference, p<0.001, *ρ* = 0.82 for *R* difference, p<0.001; significance assessed via two-tailed t-tests, Fig. 5D). In line with ROC results, the *ATC* regressor showed modest correlation (*ρ* = 0.05, n.s., Fig. 5D). Importantly, all AUC-related findings and regression weight correlations were consistent and statistically significant across both individual subjects (Supp. Fig. S6).

The second spike-rate model combined value-based variables with accumulated tokens by considering interactions between *EV, R* and *ATC* as in the ‘linear model of choice with *ATC* interactions’ (Methods 2.4.2, 3.1.2). Regressors in this model comprised the main effect *EV* difference between the two offers, the interaction of *EV* difference with *ATC*, the risk *R* difference, a linear interaction term between *R* difference and *ATC*, and a quadratic interaction term between *R* and *ATC*. This approach was motivated by prior observations indicating that the influence of *EV* difference on choice increases with *ATC* (Fig. 4C, Supp. Fig. S1C), and that the relationship between risk-seeking behavior and *ATC* follows an inverted-U profile (Fig. 4D, Supp. Fig. S1D). The introduction of interaction terms did not fundamentally alter the overall *SV* encoding profiles: the largest fractions of cells encoding *SV*s remained concentrated around offer presentation epochs (Fig. 6A-B). However, the inclusion of these interaction terms did lead to a modest increase in the fractions of cells encoding *SVs* during later task epochs in subject 2 (Supp. Fig. S7).

**Figure 6.**
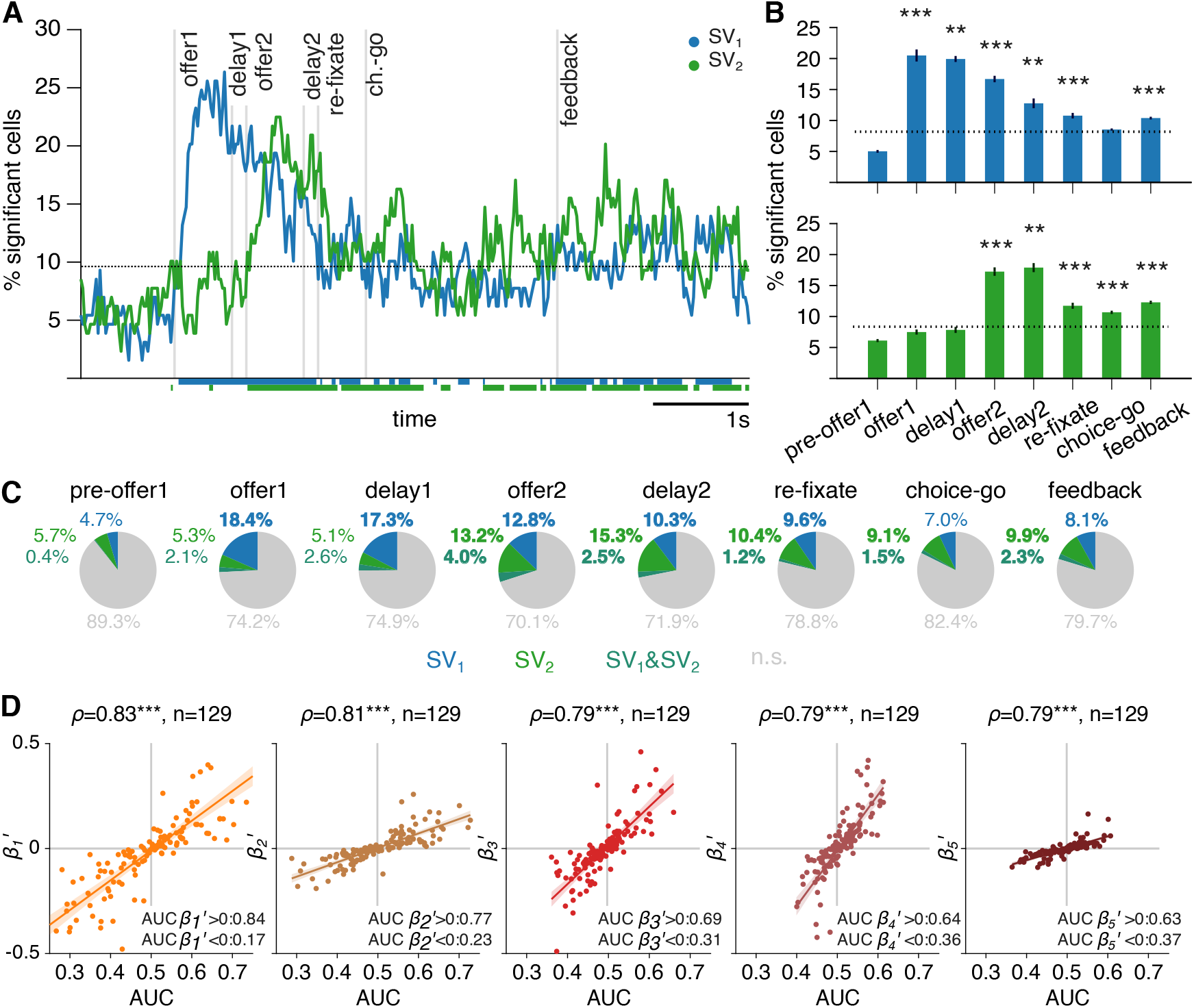
Encoding of *SV*s during task execution in the spike-rate model with *ATC* interactions and neural tuning of *EV* and *R* differences and *ATC* interactions encoding weights to choice prediction accuracy. **A.** Solid lines show the fractions of cells significantly encoding *SV*_1_ and *SV*_2_ in the ‘linear model with *ATC* interaction’ (Methods 3.1.2). The dotted lines indicate trial-order shuffles percentiles and bottom lines show significant time bins as in Fig. 5A. Results are shown for the two subjects separately in Supp. Fig. S7. **B**. Fractions of significant cells for *SV*_1_ (top) and *SV*_2_ (bottom) in task epochs (mean ± s.e.m. across time bins, *n* = 129 total cells). Dotted lines show trial-order shuffles percentiles as in Fig. 5B. Significance is assessed via one-tailed signed rank, FDR corrected via Benjamini-Hochberg procedure, testing that empirical data exceed 95^th^ percentile thresholds (\**p* < 0.05, \*\**p* < 0.01, \*\*\**p* < 0.001, Supp. Table ST10). **C**. Time-averaged fractions of *n* = 129 total cells that exclusively encode *SV*_1_ or *SV*_2_, or jointly *SV*_1_ and *SV*_2_, or are non-significant (n.s.). Results in B represent aggregate categories from C. The fraction of cells encoding *SV*_1_ includes cells exclusively encoding *SV*_1_ as well as cells encoding *SV*_1_ jointly with *SV*_2_, and similarly for *SV*_2_. Fractions in bold exceed the 95^th^ percentile of equivalent fractions from trial-order shuffles. Subject-specific results, percentiles and significance are in Supp. Fig. S8. **D**. Correlations between Area Under the Curve (AUC) from ROC analyses of choice and model weights 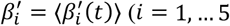 1, … 5, from left to right) time-averaged at times posterior to the offer 2 onset, for the linear model with *ATC* interactions (Methods 3.2.2). Each dot represents a cell (*n* = 129, subjects combined), with AUC and model weights averaged in time bins posterior to offer 2 onset and cross-validation folds. Pearson’s coefficients *ρ* are assessed in significance using two-tailed t-tests (\*\*\**p* < 0.001, *p* = 1.04 · 10^−33^ for 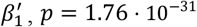, *p* = 1.76 · 10^−31^ for 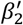, *p* = 2.09 · 10^−28^ for 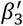, *p* = 3.51 · 10^−28^ for 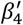, *p* = 2.25 · 10^−28^ for 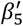). Solid lines show linear regression lines, shaded areas indicate ±CI computed using the standard error of the regression coefficients. The values at the bottom-right of each panel report the median AUC across time bins posterior to offer 2 onset, cells, cross-validation folds for positive (top) and negative (bottom) weights. Results are shown separately for the two subjects in Supp. Fig. S9.

A more detailed analysis of *SV* encoding revealed that significant fraction of cells encoding the two offers either exclusively (*SV*_1_ or *SV*_2_) or synergistically (*SV*_1_ and *SV*_2_) during task epochs following the respective offer presentation, with only exception for the exclusive encoding of *SV*_1_, that was not significant in the choice-go epoch time (Fig. 6C). This pattern was prominent in cells recorded in subject 2 (n=74), and in data combined across the two subjects (n=129). In contrast, cells recorded in subject 1 (n=55) primarily showed significant encoding around offer presentation epochs (Supp. Fig. S8).

In the spike-rate model with *ATC* interactions, we assessed choice predictability using ROC analysis (Methods 3.2.2). Specifically, we assessed the AUC scores for the following regressors: *EV* difference, *EV* difference and *ATC* interaction, *R* difference, *R* difference and *ATC* interaction, *R* difference and *ATC*^2^ interaction (Methods 3.2.1). Consistent with findings from the model without *ATC* interactions, the *EV* difference regressor shows the strongest choice predictability (median AUC for *β*_1_’ > 0: 0.84, 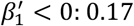, data combined for the two subjects, Fig. 5D). This was followed by the *EV* difference and *ATC* interaction (median AUC for *β*_2_’ > 0: 0.77, *β*_2_’ < 0: 0.23, Fig. 5D), with gradually declining predictability for the Risk difference regressors (median AUC for *β*_3_’ > 0: 0.69, *β*_3_’ < 0: 0.31, Fig. 5D), *R* difference and *ATC* (median AUC for *β*_4_’ > 0: 0.64, *β*_4_’ < 0: 0.36, Fig. 5D), *R* difference and *ATC*^2^ (median AUC for *β*_5_’ > 0: 0.63, *β*_5_’ < 0: 0.37, Fig. 5D).

All spike-rate regression weights showed strong and significant correlations with their respective AUC values, with similar Pearson correlation coefficients across regressors (*ρ* = 0.83 for *EV* difference, p<0.001; *ρ* = 0.81 for *EV* difference and *ATC* interaction, p<0.001; *ρ* = 0.79 for *R* difference, p<0.001; *ρ* = 0.79 for *R* difference and *ATC*, p<0.001; *ρ* = 0.79 for *R* difference and *ATC*^2^; significance assessed via two-tailed t-test). All results from these AUC analyses were significant and consistent in the two subjects (Supp. Fig. S9).

The RDV spike-rate model enabled the classification of offer value variables in gains and losses based on their relationship between a token-dependent reference value, *r*(*ATC*). This reference shifted dynamically: it was set to 0 when jackpot was not achievable and changed to the number of tokens missing to reach the jackpot (6 − *ATC*) once reaching the jackpot was possible. The fraction of values resulting in gains (≈ 43%, Supp. Table ST12) tended to be lower than the number of values resulting in losses (≈ 57%, Supp. Table ST12) as defined by the RDV utility function (Methods 2.4.3). Within each offer, when the value cued by the top portion of offers resulted in a gain, the value of the bottom portion tended to result in a loss. However, there was very little interdependence between the two offers, as confirmed by gains/losses correlations (Supp. Table ST13).

The RDV spike-rate analysis showed that a substantial fraction of dACC cells significantly encoded *SV*s derived from the reference-dependent utility function (Methods 2.4.3, 3.1.3). Like in the other models that we tested, the highest fraction of SV-encoding cells was observed around offer presentation epochs (Fig. 7A-B). Significant *SV* encoding tended to persist in later task epochs for both gains and losses (Fig. 7A-B, mainly in subject 2 cells Supp. Fig. S10). For *SV*_1_ gains, the fractions of significantly encoding cells were significantly higher than for *SV*_1_ losses in almost all task epochs following offer 1 presentation, with exception of the delay 2 epoch (Fig. 7B). For *SV*_2_ gains, the fractions of significantly encoding cells were significantly higher than for *SV*_2_ losses at offer 2, delay 2 and feedback (Fig. 7B).

**Figure 7.**
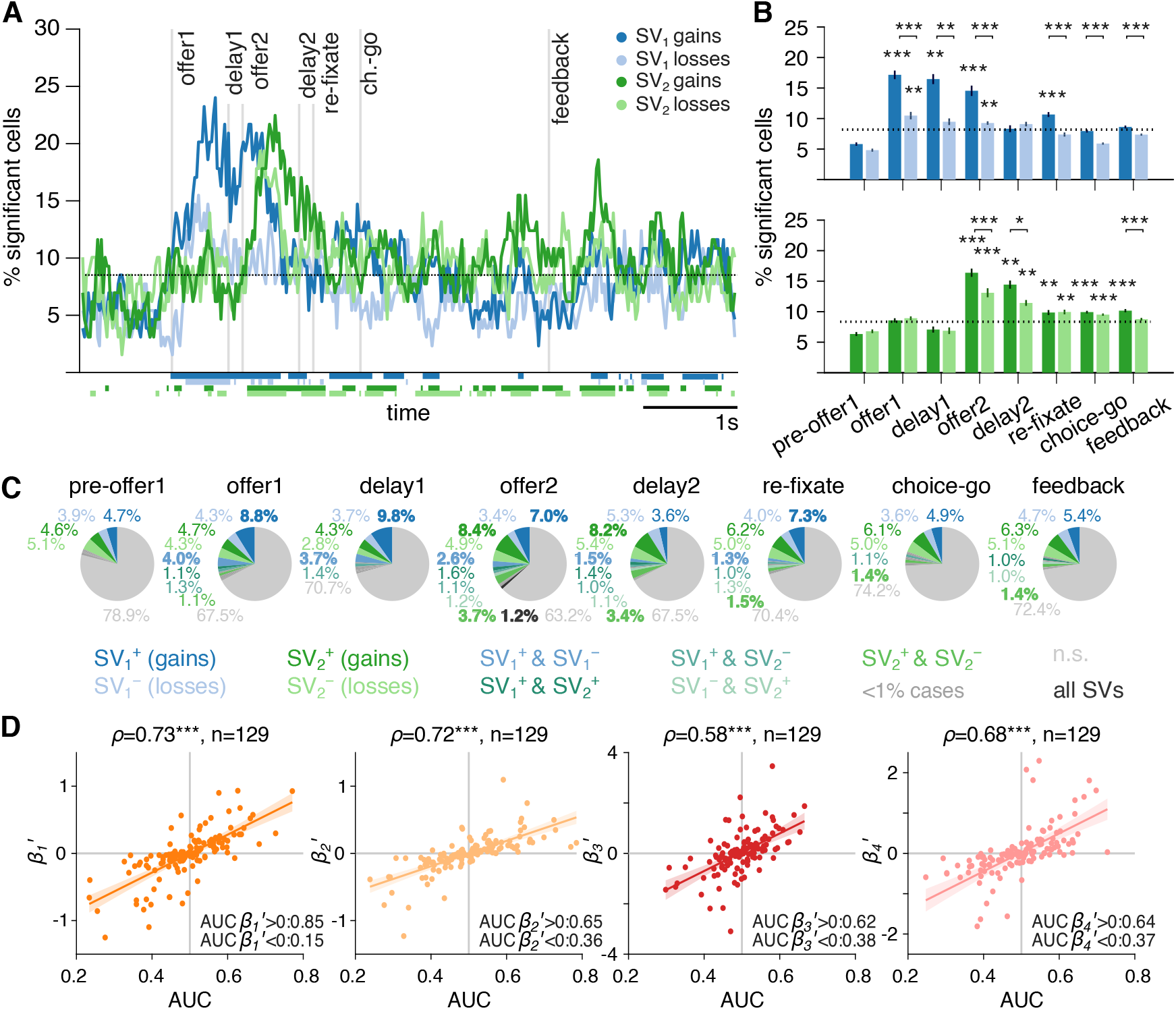
Encoding of *SV*s for relative gains and losses during task execution in the spike-rate RDV model and neural tuning of *EV* and *R* differences encoding weights in gains and losses to choice prediction accuracy. **A.** Solid lines show fractions of cells significantly encoding *SV*_1_ gains, *SV*_1_ losses, *SV*_2_ gains and *SV*_2_ losses in the ‘RDV spike-rate model’ (Methods 3.1.3). The dotted lines indicate trial-order shuffles percentiles and bottom lines indicate significant time bins as in Fig. 5A. Results are shown for the two subjects separately in Supp. Fig. S10. **B**. Fractions of significant cells for *SV*_1_ (top) gains (dark) and losses (light), and for *SV*_2_ (bottom) gains (dark) and losses (light) in task epochs (mean ± s.e.m. across time bins, *n* = 129 total cells). Dotted lines indicate trial-order shuffles percentiles as in Fig. 5B. Significance is assessed via one-tailed signed rank, FDR corrected via Benjamini-Hochberg procedure, testing that empirical data exceed the 95^th^ percentile thresholds (\**p* < 0.05, \*\**p* < 0.01, \*\*\**p* < 0.001, Supp. Table ST11). **C**. Time-averaged fractions of *n* = 129 total cells that encode *SV*_1_ gains 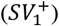, *SV*_1_ losses 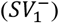, *SV*_2_ gains 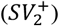, *SV*_2_ losses 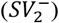 exclusively, jointly encode *SV*s (e.g., 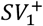 and 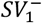), or are non-significant (n.s.). We group joint cases that show significant in minor quantity (< 1% of total cells). Results in B represent aggregate categories from C. For example, the fraction of cells encoding 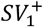 includes cells exclusively encoding 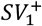 as well as cells encoding it jointly to 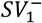 and/or 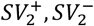. Fractions in bold exceed the 95^th^ percentile of equivalent fractions from trial-order shuffles. Subject-specific results, percentiles and significance are in Supp. Fig. S11. **D**. Correlations between Area Under the Curve (AUC) from ROC analyses of choice and model weights 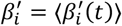 (*i* = 1, … 4, from left to right) time-averaged at time posterior to the offer 2 onset, for the RDV model (Methods 3.2.3). Each dot represents a cell (*n* = 129, subjects combined), with AUC and model weights averaged in time bins posterior to offer 2 onset and cross-validation folds. Pearson’s correlation coefficients *ρ* are assessed in significance via two-tailed t-tests (\*\*\**p* < 0.001, *p* = 1.21 · 10^−22^ for 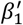, *p* = 1.62 · 10^−21^ for 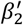, *p* = 7.84 · 10^−13^ for 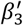, *p* = 7.46 · 10^−15^ for 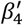). Solid lines show linear regression lines, shaded areas indicate ±CI computed using the standard error of the regression coefficients. The values at the bottom-right of each panel report the median AUC across time bins posterior to offer 2 onset, cells, cross-validation folds for positive (top) and negative (bottom) weights. Results are shown separately for the two subjects in Supp. Fig. S12.

We further analyzed encoding patterns by testing whether cells exclusively or synergistically encoded gains and/or losses for the two *SV*s. Significance was determined by comparing empirical fractions to the 95^th^ percentile of equivalent fractions from trial-order shuffles. We found that as soon as offers were presented, the respective *SV*s were significantly encoded exclusively for gains, or in synergy for gains and losses, but not significantly for losses exclusively (Fig. 7C). During offer 2, we found a modest, though significant fraction of cells encoding all *SV*s, including *SV*_1_ and *SV*_2_, in both gains and losses (Fig. 7C). Consistent with previous analyses, we found that the fractions of significant cells gradually decayed after offer presentation epochs (Fig. 7C).

Finally, we assess how the neural tuning to *SV* in gains and losses related to the behavioral choice, evaluating choice predictability using ROC analysis (Methods 3.2.3). We used as regressors the *EV* and *R* variables defined over reference-dependent values (Methods 2.4.3), by keeping gains and losses in separate regressors. We found that reference-dependent *EV* difference in gains is remarkably more accurate in choice prediction (median AUC for 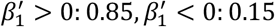, Fig. 7D) than *EV* difference in losses (median AUC for 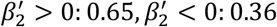, Fig. 7D) or reference-dependent Risk difference in gains (median AUC for 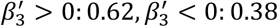, Fig. 7D) or losses (median AUC for 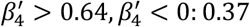, Fig. 7D).

The spike-regression weights had strong and statistically significant correlations with their respective AUC values, with *EV* difference regressors showing higher Pearson correlation coefficients across regressors (*ρ* = 0.73 for *EV* difference in gains, *p* < 0.001; *ρ* = 0.72 for *EV* difference in losses, *p* < 0.001; *ρ* = 0.58 for *R* difference in gains, *p* < 0.001; *ρ* = 0.68 for *R* difference in losses, p<0.001; significance assessed via two-tailed *t*-tests). Importantly, these AUC-based results were consistent in the two subjects (Supplementary Figure S12).

The RDV model results refine previous spike-rate model designs by allowing us to categorize value-based regressors as gains or losses and extract the fraction of cells recruited for value encoding in the two cases. This specific aspect is central to our investigations, as assessing a reference-based method for neural encoding detection can be impacted by data size, in a way that low trial availability can lead to miss the detection of significant encoding, yet lower trial availability does not increase the false alarm rate. As control analysis, we have repeated results in the ‘linear model without *ATC* interactions’ and the ‘linear model with *ATC* interactions’ by considering value respectively as in Methods 3.1.1 or Methods 3.1.2 but by splitting value variables labelled as gains or losses by the RDV utility function defined in Methods 2.4.3. This showed two main results: reducing trial counts by splitting value variables into gains and losses reduced the encoding detection power of spike-rate models (Supplementary Figures S13 and S15); regression weights became correlated or anti-correlated with AUCs depending on whether gain or loss trials were analyzed (Supplementary Figures S14 and S16). Lastly, to rule out the possibility that our findings were artifacts of uneven trial availability in gain versus loss trials, we performed a trial-matching control. We randomly sub-sampled trials (regardless of gain/loss classification) to match the number of gain and loss trials used in the split analyses for both the ‘linear spike-rate model without *ATC* interaction’ (Supp. Fig. S17) and the ‘linear spike-rate model with *ATC* interactions’ (Supp. Fig. S18). This approach yielded similar reductions in encoding detection, confirming that data size alone impacts sensitivity. Importantly, however, in control analyses (Supp. Fig. S17-S18) loss trials consistently showed a higher fraction of significantly encoding cells, consistent with their greater prevalence (≈57% of trials vs. ≈43% gains, Supp. Table ST12). This control supports the robustness of our main finding that gains are encoded by a higher fraction of neurons, despite being less frequent; if the observed effects were driven by trial count, we would expect the opposite predominance pattern seeing losses encoded by higher fractions of cells than gains. Lastly, we also observed that *SV*^+^ always assume positive value since both *EU*^+^ and *VU*^+^ terms are positive, but that *SV*^−^ could be either positive or negative, since *EU*^−^ is negative but *VU*^−^ is positive. This asymmetry fairly follows the RDV utility definition, and is also present in the RDV choice model, which best explains behavioral data. We have checked that our main results suggesting that *SV* encoding in gains entails larger fractions of cells than in losses also holds by flipping the sign of the expected utility *EU* in losses, thus making 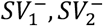 positive 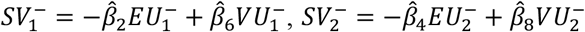 (Methods 3.1.3, Supp. Fig. S19) and by applying neural spike-rate analyses on z-scored 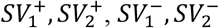 (Supp. Fig. S20).

## Discussion

This study shows that the accumulation of multi-trial tokens toward a final jackpot significantly shapes trial-based decision-making, both behaviorally and at the neural level. The accumulation of virtual tokens strongly influenced the accuracy and speed of subjects solving the task: choices were faster and more accurate for higher *ATC*, particularly when jackpot achievement was more likely. Logistic modeling confirmed a significant increase in the slope of the choice accuracy for high *ATC* conditions, corroborating the idea of stronger engagement when jackpot proximity increased. These effects coincided with a negative trend in errors when more tokens were accumulated, and with a lower propensity for risk-taking whenever the cumulative jackpot approached. Importantly, these results were also modulated by task difficulty, with easier trials showing substantially fewer errors and risky choices.

To capture the mechanisms underlying these behavioral adjustments, we introduced three alternative definitions of *SV*, combining trial-specific, value-based offer attributes *EV* and *R* and *ATC*. We formalized the assessment of our findings by testing three alternative logistic choice models for *ATC* integration and showed that offer evaluation is best modelled by a dynamically adapted reference-dependent value ‘RDV’ that follows cumulative token accumulation. The RDV model allows the decision contingencies to adapt from the comparison of offer values when it is not possible to achieve jackpot, to the comparison of offer value to the jackpot threshold when jackpot is can be achieved, i.e., to dynamically switch from value-based to goal-oriented task solving strategies. Model comparisons revealed that RDV best explains behavior, suggesting that decisions are best modelled as dynamic, reference-dependent processes tied to cumulative token progress rather than trial-specific value differences alone, even when including relevant interaction terms between offer values and *ATC*.

In addition to token-dependent behavioral effects, we also included model comparisons for temporal discounting models (Supp. Materials, Supp. Tables ST14-ST15, Supp. Figure S21). In temporal discounting, the SVs scale with time delay *D* to reward achievement, following an exponential factor *e*^−*kD*^. In our formulation, we propose that temporal discounting relates to token accumulation through the exponent *kD* ≈ (6 − *ATC*)/*τ* (Supp. Materials), with *τ* ≈ 3.4 (Supp. Table ST12), fit via maximum likelihood estimation. Inverting the above equation, we estimate effective time-discount rates of *k* ≈0.88 Hz for *ATC* = 0, and ≈0.15 Hz for *ATC* = 5, comparable to reports from previous studies^49,50^. However, in line with works showing that conventional intertemporal choice tasks often overestimate discounting and may capture heuristic, reward-rate maximizing strategies rather than genuine devaluation of delayed outcomes^51,52^, adding time-discounting terms to our models did not improve predictive performance beyond RDV. Crucially, the regression weight magnitudes for RDV coefficients remained stable across combined variants (Supp. Table ST12), indicating that RDV captures the substantial and behaviorally meaningful component of subjective decisions based on expected outcomes (gains/losses) observed across token accumulation (Supplementary Methods, Supp. Tables ST14-ST15, Supp. Fig. S21).

Overall, these findings extend previous work on token-based paradigms, where progressive, multi-trial reference effects did not previously emerge^43,45^. While our results align with previous behavioral theories for subjective preferences^1,53,54^, our data demonstrate that reference-dependent dynamics strongly influence accuracy and response time at cumulative stages before jackpot attainment, with measurable effects on best choice discrimination accuracy and risk-taking attitude. The integration of positive or negative tokens as reinforcers refines the understanding of how virtual rewards and punishments modulate risk-taking and learning, aspects of investigation typically only studied at a coarser level^55,56^.

At the neural level, we tracked the encoding of *SV*s defined by the three models that were used for behavioral choice assessment. Across all models, we found that the encoding of offer *SV* involved the most prominent portion of dACC cells during the respective offer presentation and gradually faded across subsequent task epochs. The first neural encoding model that we considered did not include *ATC* in the *SV* definition but included it as independent regressor. This showed that *ATC* is encoded by large fractions of cells in dACC, often beyond the extent of *SV* encoding fractions, especially outside offer presentation epochs. By correlating the neural encoding weights for the *EV* and *R* differences and the AUC computed via ROC analyses for choice prediction, we found that *EV* difference weights were most aligned in tuning with choices than other variables considered. Crucially, this also revealed that, as expected, the encoding tuning of *EV* and *R* differences variables is aligned to choices, whereas the encoding tuning of *ATC* is not aligned to choices. The second neural encoding model that we considered included *ATC* interaction terms in the definition of *SV*s. Comparing results to first model, we observed that including *ATC* in *SV*s led to higher fractions of cells significantly encoding *SV*s at later task times, and that correlating the encoding weights of *EV* and *R* differences to the AUC of choice predictions results in lower alignment in encoding tuning when *EV* and *R* difference terms interact with *ATC*. This latter result indicates that *ATC* interaction with *EV* and *R* does not provide choice-related tuning adjustment at the neural level. By using a token-dependent reference, we introduced a third neural encoding model that followed the ‘RDV’ choice model best explaining behavioral data that allowed us to consider value-based variables as gains or losses based on their relationship with an *ATC*-dependent reference. Notably, this revealed that significantly larger fractions of dACC cells encode *SV*s resulting from gains. By analyzing the fractions of cells exclusively or synergistically encoding gains- and/or losses-related variables, we found that that differently from gains, the losses-related variables were not exclusively encoded by significant fractions of cells, but that they were always encoded in synergy with gain-related variables. In addition, we found that the neural encoding of *EV* in gains was robustly aligned to choices, whereas this was not the case in losses. In contrast to behavioral insights suggesting that ‘losses hurt more than gains’, neural analyses result indicate that dACC possibly follows a reference-dependent neural tuning that aligns value encoding weights to choice mainly for gains, and less for losses. We further verified that adopting a model combining reference-dependency with time-discounting yields a qualitative match to the neural results (Supp. Fig. S22), supporting the evidence that reference-dependent valuation provides an explanatory framework for both behavioral adjustments and dACC encoding in our task.

Converging evidence suggests that expected rewards are updated by experience and estimated in the frontal cortex and basal ganglia^57^. Our evidence that dACC encoding follows reference-dependent principles aligns with previous theories by which dACC supports the selection and maintenance of context-specific sequences of behavior directed toward delayed goals^58. 5110^ Our results extend previous findings relating ACC signals to post-decisional variables related to gains or losses ^42,55,59^, as well as to context-dependent attentional drive^60,61^, elaborating on previous interpretations about dACC not being directly involved in ongoing decisions but in outcome evaluations^44^. Remarkably, we find that dACC recruits larger fractions of cells for RDV gains than for losses, and that their encoding tuning aligns with upcoming choices, linking gains to reward anticipation ^16^, and connecting to theories for distinct specialized circuitry for reward anticipation and punishment across brain structures^62,63^. This may have notable implications, linking the observed reduction in risk-taking and faster responses near jackpot completion to motivational shifts possibly mediated by dopaminergic modulation^64^. This reinforcement learning perspective suggests prioritization of rewards maximization^65–68^, by enhancing of neural encoding for positive expected outcomes.

Furthermore, the involvement of the dACC in token accumulation reinforcement is particularly noteworthy. Previous research has implicated the dACC in cognitive control^7,69^, conflict monitoring^23–25^, and value-based decision-making^12,14,26,29^. Our findings further support its role in encoding the subjective value of outcomes in a dynamic, multi-trial context, adjusting its representation as the likelihood of jackpot attainment increases. This suggests that dACC supports ongoing monitoring of a probabilistic, history-dependent reference frame, an often overlooked dimension of the putative functional role of value encoding in dACC. The dynamic influence of tokens accumulation towards a jackpot reward provides a critical advancement for realistic decision-making models that incorporate history-dependent value signals and reinforcement-driven, reference-dependent shaping in reward probability^2–5,70,71^.

In conclusion, this study underscores the importance of considering accumulated rewards in decision-making frameworks. By showing how both behavior and dACC encoding adapt to reference-dependent, multi-trial dynamics, we extend current understanding of the neural computations supporting value-based and goal-directed behavior in environments where reward attainability context unfolds over time.

## Methods

### 1. Experimental settings and reproducibility

The data includes n = 227 behavioral sessions from two subjects (109 sessions, 433.28 ± 8.70 mean ± s.e.m. trials per session for subject 1, 118 sessions, 500.68 ± 10.97 trials for subject 2), of which n = 108 sessions (65 sessions, 479.16 ± 14.57 mean ± s.e.m. trials per session for subject 1, 43 sessions, 519.35 ± 14.62 trials for subject 2) include extracellular activity of dACC (Area 24) recorded using single electrodes (Frederick Haer & Co., impedance range 0.8 ± 4 MΩ), while monkeys performed the task. The electrode probes were lowered using a microdrive (NAN Instruments) until well-isolated activity was encountered. Data covered 1-3 simultaneous single neurons, Plexon system (Plexon, Inc.) was used to isolate the action potential waveforms. Cells are included based on quality of isolation but not on task–related response properties. Here we present neuronal data consisting of *n* = 129 single units (55 for subject 1 and 74 for subject 2). These data were published in the context of different research questions^39,40^.

#### 1.1 Experimental Procedures and Neural Recordings

All procedures were approved by the University Committee on Animal Resources at the University of Rochester and were designed and conducted in compliance with the Guide for the Care and Use of Animals of the Public Health Service’s Guide for the Care and Use of Animals (protocol UCAR-2010-169). Two male macaques (*Macaca mulatta*; subject 1: age, 8 years, 11 months; subject 2: age, 10 years, 9 months) served as subjects. Recorded brain regions covered dACC, using single-contact probes. Data include 47228 behavioral trials (433.28 ± 8.70, mean ± sem across sessions) from subject 1 and 59080 (500.68 ± 10.97) from subject 2. Of these, 20604 (479.16 ± 14.57) from subject 1, and 33758 (519.35 ± 14.62) from subject 2 were simultaneous to dACC data recordings.

#### 1.2 Behavioral Task

The token-based reward gambling task consists of the sequential presentation of two visual stimuli at the opposite sides of the screen, providing reward offers to be subsequently chosen by performing saccade to either target location (Fig. 1A)^21,38–40^. The task starts with a first offer presentation (*offer 1*, 600 ms), followed by a first delay time (*delay 1*, 150 ms), a second offer presentation (*offer 2*, 600 ms), respectively followed by a second delay time (*delay 2*, 150 ms). After fixating a central cross (*re-fixate*), for at least 100 ms, the choice can be reported upon *choice-go* cue onset, consisting of the simultaneous presentation of both offer stimuli previously presented. Choice is reported by fixation on target offer cue for at least 200 ms (*choice*). A successful choice selection fixation was followed by 750 ms delay period after which the gamble was resolved. Regardless of the outcome, within the next 300 ms a small juice reward (100 μL) was delivered to keep subjects engaged in the task. If, after the outcome resolution less than six tokens are collected, the trial ends. If six or more tokens are collected, after a 500 ms delay period a “jackpot” large liquid reward (300 μL) is delivered, the token count is reset, and the trial ends. Trials were separated by a random inter–trial interval ranging between 0.5 and 1.5 s. Circular indicators at the bottom of the screen report the token count through task time as empty circles, filled according to token count. The visual offer stimuli are split horizontally into top and bottom parts of different colors, with associated reward magnitude and probability. The height of the top/bottom part is informative of the associated reward probability, while the color is informative of the magnitude. Thanks to the split, the probability of the outcomes color-cued by the top and bottom parts of the stimuli is always complementary. The probabilities are discretized as 10%, 30%, 50%, 70% and 90%, the magnitudes consist of reward counts (−2, −1,0, +1, +2, +3 tokens) including negative values. In addition, offers include safe options where 0 (red) or 1 (blue) token are achieved with 100% probability. Each gamble includes at least one positive or zero outcome. This allows a less trivial level of choice computation and keeps the subjects motivated throughout behavioral execution.

### 2. Behavioral Data Analyses

We analyzed behavioral data at multiple levels. First, we assessed the factors that influenced the subjects’ choices. The visual offers cued at magnitude *v* and probability *p* of the top (*t*) and bottom (*b*) parts of the offers, e.g., in Fig. 1A we show a sample configuration where the left offer is of *v*^*t*^ = +3 tokens, and *v*^*b*^ = −2 tokens, while the right offer has *v*^*t*^= +1 tokens, *v*^*b*^ = −1 tokens. We defined the expected value as: *EV* = *p*^*t*^*v*^*t*^ + *p*^*b*^*v*^*b*^, where *p*^*t*^ and *p*^*b*^ = 1 − *p*^*t*^ are indicated by the height fraction of the top or bottom offer segments respectively. Similarly, we defined the risk associated with the offers as: *R* = *p*^*t*^(*v*^*t*^ − *EV*)^2^ + (1 − *p*^*t*^)(*v*^*b*^ − *EV*)^2^. In behavioral data analyses, we stratified the analyses in ranges of accumulated tokens: ‘Low’, ‘Medium’, ‘High, and in levels of difficulty in detecting the offer with best *EV*: ‘Easy’ and ‘Hard’. As accumulated tokens count ranges, we use ‘Low’ *ATC* = [0, 1], ‘Medium’ *ATC* = [2, 3], and ‘High’ *ATC* = [4, 5] (Supp. Table ST1).We defined difficulty as inversely related to the absolute *EV* difference Δ_*EV*_= |*EV*_1_ − *EV*_2_|, i.e., larger Δ_*EV*_ indicate lower difficulty. We used the median value of Δ_*EV*_, *median*(Δ_*EV*_) = 1 as discriminant for ‘Easy’ (Δ_*EV*_≥1) and ‘Hard’ (Δ_*EV*_< 1) trials (Supp. Table ST2).

#### 2.1 Generalized linear model of choice

To assess the choice performance, in Fig. 2A we used a logistic model for the ‘correct choice’ *CC*, defined as choice for the offer with best *EV*. In symbols, *CC* = 1 if choice is for offer 1 when *EV*_1_ > *EV*_2_, and for offer 2 when *EV*_2_ > *EV*_1_; *CC* = 0 in all other cases. The logistic model assumes a logistic relationship between the correct choice *CC* and the above variables: logit(*CC*) = *β*_0_ + *β*_1_Δ_*EV*_ + *β*_2_*M*_*EV*_ + *β*_3_*ORL* + *β*_4_*ATC* + *β*_5_*TSLR* + *β*_6_*OPT* + *β*_7_*JPT*. We considered as regressors: the absolute *EV* difference between the two offers (Δ_*EV*_); the ‘Mean expected value’ as the average *EV* in each trial: *M*_*EV*_ = (*EV*_1_ + *EV*_2_)/2; the ‘Offer Risk Level’ *ORL* = *R*_*Best*_ − *R*_*Worst*_ where *R*_*Best*_ or *R*_*Worst*_ indicates the risk associated to the offer with best or worst *EV*, respectively; the ‘Accumulated Tokens Count’ *ATC*, i.e., the token count at the start of the trial; the ‘Trials Since Last Reward’ *TSLR* that is, the count, in number of trials, since last jackpot reward; the ‘Outcome of Previous Trial’ *OPT*, that is the number of tokens achieved in previous trial; and the ‘Jackpot on Previous Trial’ *JPT*, a binary variable = 1 if previous trial ended with jackpot, = 0 otherwise. Due to using previous trial variables, the first trial in each session was removed in this analysis (*n* = 47030 total trials in subject 1, *n* = 58911 total trials in subject 2). Prior to least squares estimation of the weights *β* _*i*= [0,7]_, the regressed variables are normalized to their absolute maximum value so that (Δ_*EV*_, *ATC, TSLR, JPT*) are in the range [0, 1], and (*M*_*EV*_, *ORL, OPT*) are in the rage [-1, 1], and assessed for significance via two-tailed *F*-Statistics tests.

#### 2.2 Behavioral choice accuracy and execution time

In Fig. 2B we compute the fraction of trials with correct choice (i.e., choice for the offer with best *EV*) for ‘*No Jackpot*’ (JPT = 0) or ‘*Jackpot*’ (JPT = 1) on previous trial, for ‘Low’ *ATC* = [0, 1], ‘Medium’ *ATC* = [2, 3], and ‘High’ *ATC* = [4, 5] accumulated tokens count (Supp. Table ST1). The same conditions are used in Fig. 2C to analyze the fraction of errors (i.e., trials where the subjects did not choose the offer with best *EV*), as well as for binned levels of Δ_*EV*_, inversely related to the difficulty in detecting the best offer (Supp. Table ST2). Finally, in Fig. 2D we show the logistic relationship between expected value difference *EV*_1_ − *EV*_2_ and the choice logit(*p*(*ch* = 1)) = *β*_0_ + *beta*_1_(*EV*_1_ − *EV*_2_) for ‘Low’ and ‘High’ accumulated tokens count (*ATC* = [0,1] or *ATC* = [4,5] respectively). We assess the difference in *β*_0_, *β*_1_ for low and high *ATC* in Fig. 2E.

#### 2.3 Risk propension

The Markowitz risk return model is defined via the weights of the logistic regression of offer choice (*ch1*= 1 for first offer choice, 0 otherwise) as logit(*ch*1) = *β*_0_ + *β*_1_(*EV*_1_ − *EV*_2_) + *β*_2_(*R*_1_ − *R*_2_). The subject’s utility for the *i*^*th*^ offer is modelled as a trade-off between *EV* and risk, defined as *U*_*i*_ = *EV*_*i*_ − *θR*_*i*_. The model parameter *θ* = −*β*_2_/*β*_1_, describes the behavioral attitude as risk aversion (*θ* > 0) or risk seeking (*θ* < 0).

#### 2.4 Choice models and Subjective Values

The definition of Subjective Value (*SV*) followed the assessment of alternative models of choice prediction, including task variables such as value of the top and bottom part of the offers (*v*^*t*^, *v*^*b*^), expected values (*EV*_1_, *EV*_2_), risks (*R*_1_, *R*_2_) and *ATC*. We considered three alternatives:’linear model of choice without *ATC* interaction’, ‘linear model of choice with *ATC* interaction’, and a ‘reference-dependent model of choice’.

##### 2.4.1 Linear model of choice without *ATC* interaction

We define a linear model the choice as:

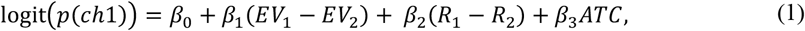

with *ch*1 = 1 indicating choice for the first offer, *ch*1 = 0 for the second offer. The model includes separate terms for *EV, R*, and *ATC*. The model weights 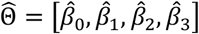 are estimated via ordinary least squares regression, i.e., by minimizing the negative log-likelihood of the predicted variable

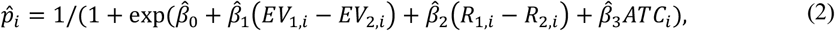

for test data *y*_*i*_ = 1 if first option is chosen, 0 otherwise. We use *k* = 4 fold cross-validation to estimate parameters over train subsets and test predictions using estimated parameters over test subsets. We apply a Ridge penalty to the negative log-likelihood, i.e., we minimize

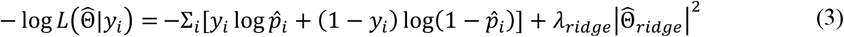

with *λ*_*ridge*_ = 0.1 and 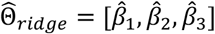.

Following this model of choice, we define subjective values as:

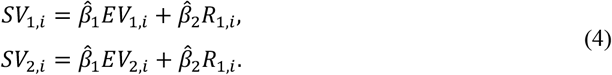

We include an *ATC* term in this logistic model for two main reasons. First, it allows us to explicitly isolate the contribution of *ATC* without conflating its effect with offer *EV* and *R*. Second, it controls for potential *ATC*-related variability that could otherwise bias the estimation of *β*_1_ and *β*_2_, ensuring that *EV* and *R* related weights remain interpretable. While this model does not include interaction terms between *ATC* and the other predictors, the estimated effect of *ATC* was very weak ( 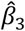 subject 1: 0.03 ± 0.01; subject 2: 0.01 ± 0.01, Supp. Fig S2, Supp. Table ST7).

##### 2.4.2 Linear model of choice with *ATC* interaction

We extend the linear model of the choice to include *ATC* interaction terms:

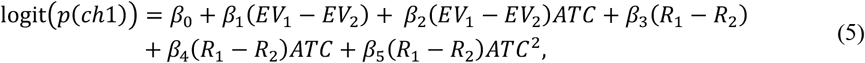

combining a linear interaction for *EV* with linear and quadratic interaction terms for *R*. This design was inspired by results in Fig. 4 and Supp. Fig. S1, showing linear increase in *EV* with *ATC*, and inverted-u relation between *R* and *ATC*.

Also in this case, we estimate the model weights 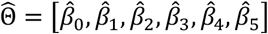 via ordinary least squares regression, i.e., by minimizing the negative log-likelihood of the predicted variable

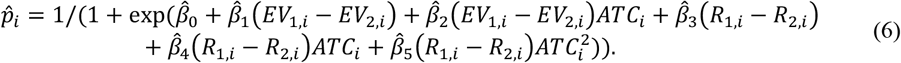

We use *k* = 4 fold cross-validation and Ridge penalty with *λ*_*ridge*_ = 0.1 and 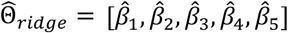. Following this model of choice, we define subjective values as:

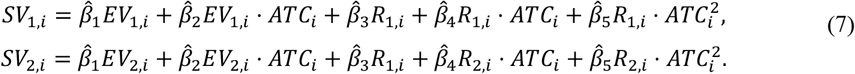

#### 2.4.3 Reference-dependent value (RDV) model of choice

We define a reference-dependent value ‘RDV’ model based on Prospect Theory principles. This model design relies on a token-based reference-dependent utility function for offer values 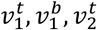, and 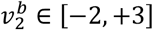 :

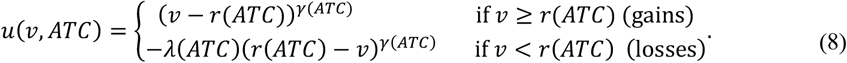

Here, the reference point *r*(*ATC*) is token-dependent and defined as:

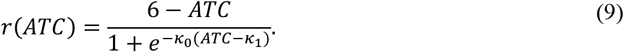

The *r*(*ATC*) function tends toward 6 − *ATC* when *ATC* ≥ *κ*_1_, and toward 0 for *ATC* < *κ*_1_. The formulation is grounded in the intuition that subjects consider offered values as potential “gains” or “losses” relative to a “missing tokens to jackpot” threshold (6 − *ATC*) when jackpot is achievable, or they consider offer values relative to zero when jackpot is not achievable. In our data, *κ*_1_ is numerically fit to data in each subject: 3.29±0.05 in subject 1, 3.08±0.06 in subject 2 (mean ± s.e.m). These values are consistent with using 6 − *ATC* as reference when jackpot is reachable (i.e., *ATC* ≥ 3, for *v* ∈ [−2, +3] allows *ATC* + *v* ≥ 6), and 0 when *ATC* jackpot is not reachable (*ATC* < 3).

The utility function *u*(*v, ATC*) transforms the reference-dependent value |*v* − *r*(*ATC*)| via a gain-sensitive parameter *γ*(*ATC*) = *γ*_0_ + *γ*_1_*ATC*, increasing the steepness of the value-to-choice mapping in the gain domain. Losses are penalized by scaling factor *λ*(*ATC*) = *λ*_0_ + *λ*_1_*ATC* + *λ*_2_*ATC*^2^, with *λ*_0_, *λ*_1_, *λ*_2_ reflecting the Prospect Theory asymmetry principle by which “losses hurt more than gains”. The quadratic penalty also captures the inverted-U pattern in risk-taking (Fig. 4 and Supp. Fig. S1), as losses correspond with riskiest choices in our task.

Finally, the reference-dependent utilities are used in a logistic regression predicting choice:

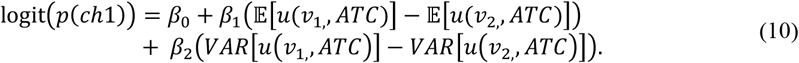

Regression was performed via ordinary least squares, using

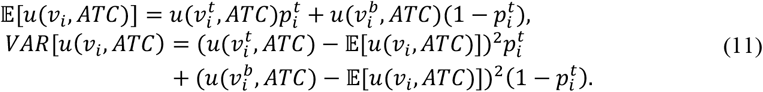

We use *k* = 4 cross-validation folds, and apply Ridge penalty to the negative log likelihood 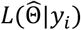 as in the linear model, estimate 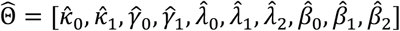, with 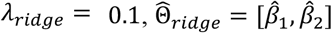, and

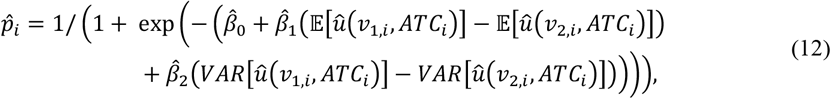

where 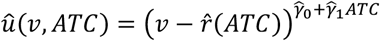 if 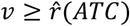 and 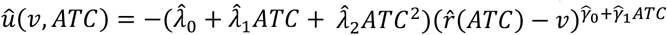 if 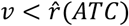, with 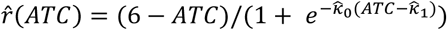.

In the RDV model, we define Subjective Values (SV) as:

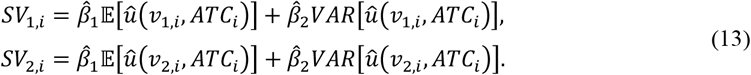

The SVs are computed estimating 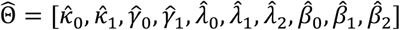 on train data and 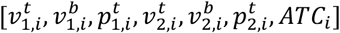 from the *i*^*th*^ test trial, using *k* = 4 cross-validation folds.

### 3. Neural data analyses

We analyzed the spiking activity of neurons in dACC to compute the fraction of cells that show significant encoding of the *SV*. Following the *SV* computation (Methods 2.4), we run a time-resolved analysis of the spike rate *η*_*i*_(*t*) of *i*^*th*^ trial at time *t* in 200 ms boxcar time windows, sliding at 10 ms offset bins during task time. We used a linear model of the spike rate, and tracked the fraction of cells showing significant encoding of the two *SV*s. The significance of *SV* interaction slope is assessed to be different from zero via two-tailed *F*-statistics tests on the sum of squared errors (*SSE*) of the full model 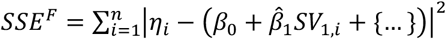, and of the reduced model 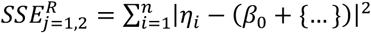, including only 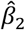 and other eventual regressors (indicated here with {…}) for the assessment of 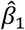, and, respectively, only 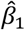 and other eventual regressors for the assessment of 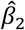. The *F*-value for *β*_*j*=1,2_ (*t*) is defined as 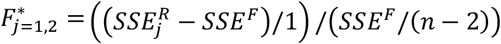, with *n* being the number of trials observations. The *p*-values are computed by comparing empirical *F*-values with the *F*-distribution. The significance of the fractions of cells showing significant encoding is further assessed via the comparison of the significant fractions of cells with surrogate results run over *n* = 1000 trial order randomizations, independently generated across variables. Lastly, consecutive runs of significant bins are assessed in length via run-length, cluster-based significance tests^20^.

#### 3.1 Spike-rate models for the encoding of value in the dACC

We investigated three models of the spike-rate derived from choice prediction frameworks outlined in Methods 2.4. The first model consisted of a linear combination of *EV* and risk variables alongside *ATC*, which we include to assess the neural encoding of such variable independently of its interactions with value-based parameters, as by the ‘choice model without *ATC* interactions’ introduced in Methods 2.4.1. The second model extended the linear approach by introducing interaction terms between *ATC* and value variables as by the ‘choice prediction model with *ATC* interactions’ detailed in Methods 2.4.2. The third model adopted a reference-dependent definition of value-based parameters, consistent with the ‘RDV model of the choice’ described in Methods 2.4.3.

##### 3.1.1 Linear spike-rate model without *ATC* interactions

To dissect the respective contribution of *EV, R*, and *ATC*, we defined a model of subjective value starting with a simplified linear model of choice, considering *SV* as a weighted sum of *EV* and *R*, and regressing the two *SV*s in a spike model with *SV*_1_, *SV*_2_ and *ATC* as in the ‘linear model of the choice without *ATC* interactions’ (Methods 2.4.1). To ensure independence between the estimation of *SV* value rates and the neural rate regression, trials were split into two disjoint subsets S1 and S2. First, we used S1 trials to compute *SV* weights via a logistic fit of the choice 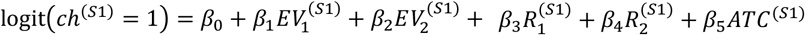. Then, weights 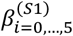, are combined with S2 data to define 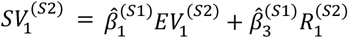. Following *SV* computation on S2 trial data, the subsets S1 and S2 are swapped, to compute *SV*s on S1, 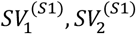. This procedure allows cross-validation by which choice-relevant *SV* weights are computed on a first subset, then combined with independent *EV, R* and *ATC* from a second, disjoint subset. Finally, SVs from the two disjoint subsets are pooled together in the set *S* and regressed against spiking data. The 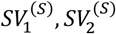 are regressed with rate model 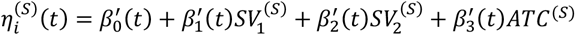, for the *i*^*th*^ cell, and time bin *t*.

##### 3.1.2 Linear spike-rate model with *ATC* interactions

We consider a spike-rate model where *EV* and *R* interact with the variable *ATC* as in the ‘linear model of the choice with *ATC* interactions’ (Methods 2.4.2). We split trials into two disjoint subsets S1 and S2 as in the ‘linear spike-rate model without *ATC* interactions’ (Methods 3.1.1). In this case the *SV* weights are computed via logistic fit of the choice model 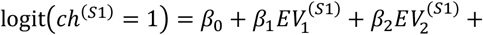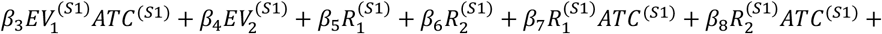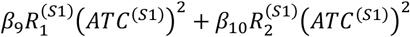. We estimate the parameters 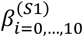, and define 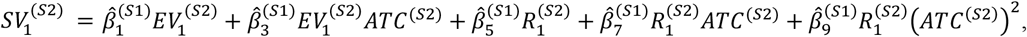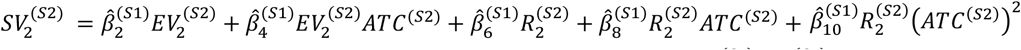. The subsets S1 and S2 are swapped to compute 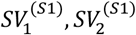. The SVs are then pooled in the set *S* as 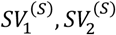 and regressed using the rate model 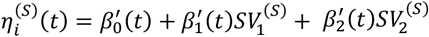, for the *i*^*th*^ cell, and time bin *t*.

##### 3.1.3 RDV spike-rate model

We use reference-dependent SVs to define RDV spike-rates models following methods in ‘Reference-dependent model of the choice’ (Methods 2.4.3). Prior to computing the SV weights, we compute the factors 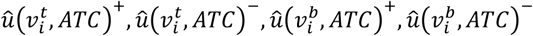, for both first and second offer *i* = 1,2, separating gains (+) from losses (-). Although offer values 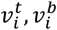 are anti-correlated by task design (*ρ* = −0.74, *p* < 0.001 for both first and second offer), gains and losses could vary in top/bottom parts (*t, b*) of the offers. We define the gains Expected Utility

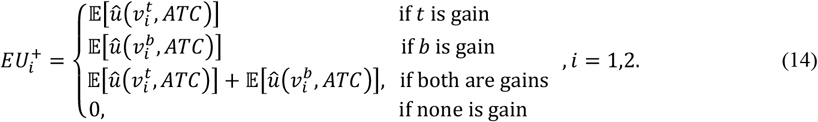

The same definition applies to losses Expected Utility 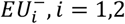, *i* = 1,2, and to the respective *VAR* utility 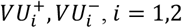, *i* = 1,2. The SV weights are computed via logistic fit of the choice model 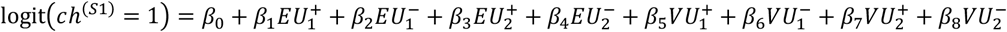, by estimating the 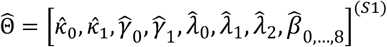 (Methods 2.4.3), and defining:

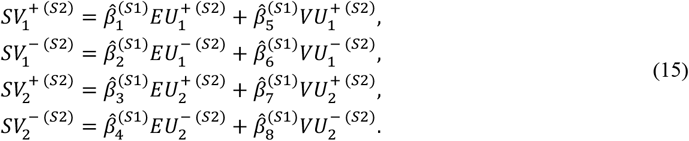

Following *SV* computation on S2, we swap S1 and S2 to compute 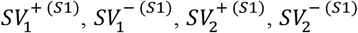. The SVs from the two subsets are pooled in the set *S* and regressed separately for gains and losses as 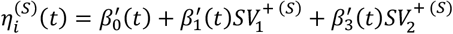 for gains, and 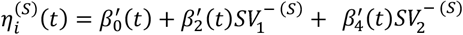 for losses, for the *i*^*th*^ cell, and time bin *t*.

#### 3.2 Receiver Operating Characteristics analysis on spike-rate models choice predictions

We refined the study of spike-rate models to study their relationship with the choice, correlating choice predicted using the spike-rate weights (*β*′ in the respective models) in a Receiver Operating Characteristics (ROC) analysis, estimating the Area Under the Curve (AUC) Importantly, this analysis allowed to detect behavioral readout^72^ from neural signals.

##### 3.2.1 ROC analysis in the linear spike-rate model without *ATC* interaction

First, we compute the weights 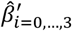 using trials in the subset S1 (Methods 3.1.1) in the model 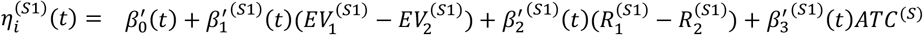. We compute predictive variables

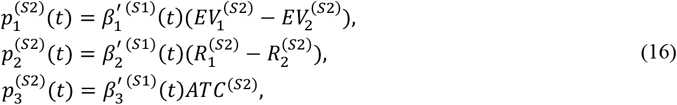

We predict the choice by sorting the variable 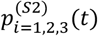, and sweeping a threshold assuming all possible values of 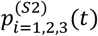. At each threshold value, we compute the true positive rate (TPR) and false positive rate (FPR) comparing the prediction variable the choice variable 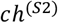, and constructing the ROC curve by tracing FPR versus TPR. The 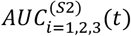 for the respective 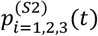 is computed as the discrete integral of the ROC curve. Lastly, we correlate the 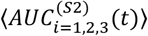 and 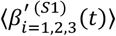 using Pearson’s correlation, ⟨⋅⟩ is the time-average across time bins posterior to offer 2 presentation and up to the end of the trial. We only used time-averages posterior to offer 2 onset time to consider time bins where subjects had access to second offer information, thus *EV* and *R* difference could be possibly encoded at the neural level.

##### 3.2.2 ROC analysis in the linear spike-rate model with *ATC* interaction

In this case we compute predictive variables based on the *ATC* interaction model (Methods 3.1.2)

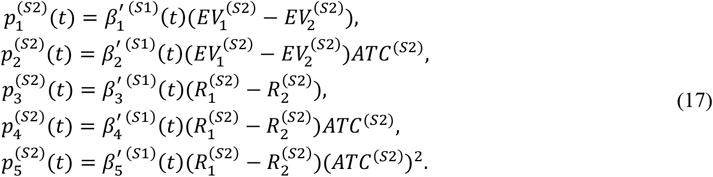

As in 3.2.1, 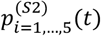 is a predictor of choice *ch*^(*S*2)^ used to compute 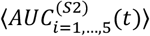 and we correlate it to 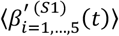.

##### 3.2.3 ROC analysis in the RDV spike-rate model

In this case, since the model needs to compute parameter estimates 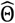 (Methods 3.1.3), we split the trials into three separate subsets. We estimate parameters 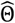 on a first subset S1, and use them to define 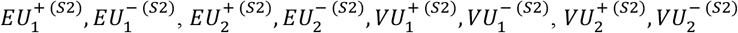 on a second subset S2. We fit the spike-rate

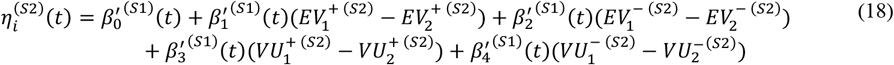

and compute:

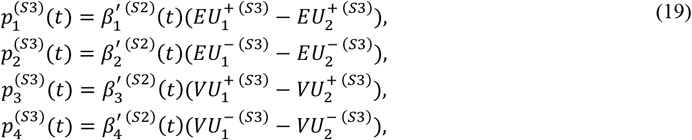

to predict the choice variable *ch*^(*S*3)^. Based on choice predictions 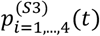, we compute the 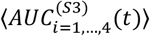 as in 3.2.1 and correlate it to 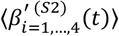.

## Data availability

Data used in this study is available at the DOI: 10.12751/g-node.1kkrw6.

## Code availability

Code developed for the presented analyses is available at the DOI: 10.12751/g-node.1kkrw6.

## Acknowledgements

This project was supported by grants funded by the Spanish Ministry of Science, Innovation and Universities (MICIU/AEI/10.13039/501100011033) and by “FEDER A way of making Europe” (ref: PID2023-146524NB), and by ICREA ACADÈMIA (2022) funded by the Catalan Institution for Research and Advanced Studies to R.M.B.

## Author contributions

D.F., R.M.B. conceptualized and designed the analyses. B.H. ideated the task. D.F. analyzed the data. B.H., H.A. collected the data. D.F., B.H., R.M.B. wrote the manuscript.

## References

1. Kahneman, D. & Tversky, A. Prospect Theory: An Analysis of Decision under Risk. vol. 47 https://about.jstor.org/terms (1979).

2. Anke Braun, X., Urai, A. E., Tobias, X. & Donner, H. Adaptive History Biases Result from Confidence-Weighted Accumulation of past Choices. Journal of Neuroscience 38, (2018).

3. Nogueira, R. et al. Lateral orbitofrontal cortex anticipates choices and integrates prior with current information. Nat Commun 8, (2017).

4. Mochol, G., Kiani, R. & Moreno-Bote, R. Prefrontal cortex represents heuristics that shape choice bias and its integration into future behavior. Current Biology 31, 1234–1244.e6 (2021).

5. Hermoso-Mendizabal, A. et al. Response outcomes gate the impact of expectations on perceptual decisions. Nat Commun 11, (2020).

6. Bartra, O., McGuire, J. T. & Kable, J. W. The valuation system: A coordinate-based meta-analysis of BOLD fMRI experiments examining neural correlates of subjective value. Neuroimage 76, 412–427 (2013).

7. Ebitz, R. B. & Hayden, B. Y. Dorsal anterior cingulate: A Rorschach test for cognitive neuroscience. Nature Neuroscience vol. 19 1278–1279 Preprint at 10.1038/nn.4387 (2016).

8. Haber, S. N. & Behrens, T. E. J. The Neural Network Underlying Incentive-Based Learning: Implications for Interpreting Circuit Disruptions in Psychiatric Disorders. Neuron vol. 83 1019–1039 Preprint at 10.1016/j.neuron.2014.08.031 (2014).

9. Heilbronner, S. R. & Hayden, B. Y. Dorsal Anterior Cingulate Cortex: A Bottom-Up View. Annu Rev Neurosci 39, 149–170 (2016).

10. Kable, J. W. & Glimcher, P. W. The neural correlates of subjective value during intertemporal choice. Nat Neurosci 10, 1625–1633 (2007).

11. Camille, N., Griffiths, C. A., Vo, K., Fellows, L. K. & Kable, J. W. Ventromedial frontal lobe damage disrupts value maximization in humans. Journal of Neuroscience 31, 7527–7532 (2011).

12. Rushworth, M. F. S. & Behrens, T. E. J. Choice, uncertainty and value in prefrontal and cingulate cortex. Nat Neurosci 11, 389–397 (2008).

13. Rushworth, M. F. S., Noonan, M. A. P., Boorman, E. D., Walton, M. E. & Behrens, T. E. Frontal Cortex and Reward-Guided Learning and Decision-Making. Neuron vol. 70 1054–1069 Preprint at 10.1016/j.neuron.2011.05.014 (2011).

14. Kennerley, S. W., Behrens, T. E. J. & Wallis, J. D. Double dissociation of value computations in orbitofrontal and anterior cingulate neurons. Nat Neurosci 14, 1581–1589 (2011).

15. Padoa-Schioppa, C. & Assad, J. A. Neurons in the orbitofrontal cortex encode economic value. Nature 441, 223–226 (2006).

16. Wallis, J. D. Orbitofrontal cortex and its contribution to decision-making. Annu Rev Neurosci 30, 31–56 (2007).

17. Rich, E. L. & Wallis, J. D. Decoding subjective decisions from orbitofrontal cortex. Nat Neurosci 19, 973–980 (2016).

18. Padoa-Schioppa, C. Neuronal origins of choice variability in economic decisions. Neuron 80, 1322–1336 (2013).

19. Strait, C. E., Blanchard, T. C. & Hayden, B. Y. Reward value comparison via mutual inhibition in ventromedial prefrontal cortex. Neuron 82, 1357–1366 (2014).

20. Ferro, D., Cash-Padgett, T., Wang, M. Z., Hayden, B. Y. & Moreno-Bote, R. Gaze-centered gating, reactivation, and reevaluation of economic value in orbitofrontal cortex. Nat Commun 15, (2024).

21. Maisson, D. J. N. et al. Choice-relevant information transformation along a ventrodorsal axis in the medial prefrontal cortex. Nat Commun 12, (2021).

22. Juechems, K., Balaguer, J., Ruz, M. & Summerfield, C. Ventromedial Prefrontal Cortex Encodes a Latent Estimate of Cumulative Reward. Neuron 93, 705–714.e4 (2017).

23. Botvinick, M., Nystrom, L. E., Fissell, K., Carter, C. S. & Cohen, J. D. Conflict monitoring versus selection-for-action in anterior cingulate cortex. Nature 1999 402:6758 402, 179–181 (1999).

24. Van Veen, V., Cohen, J. D., Botvinick, M. M., Stenger, V. A. & Carter, C. S. Anterior cingulate cortex, conflict monitoring, and levels of processing. Neuroimage 14, 1302–1308 (2001).

25. Botvinick, M. M., Cohen, J. D. & Carter, C. S. Conflict monitoring and anterior cingulate cortex: An update. Trends Cogn Sci 8, 539–546 (2004).

26. Shenhav, A., Botvinick, M. M. & Cohen, J. D. The expected value of control: An integrative theory of anterior cingulate cortex function. Neuron 79, 217–240 (2013).

27. Hadland, K. A., Rushworth, M. F. S., Gaffan, D. & Passingham, R. E. The anterior cingulate and reward-guided selection of actions. J Neurophysiol 89, 1161–1164 (2003).

28. Vassena, E., Holroyd, C. B. & Alexander, W. H. Computational models of anterior cingulate cortex: At the crossroads between prediction and effort. Front Neurosci 11, (2017).

29. Vassena, E., Deraeve, J. & Alexander, W. H. Surprise, value and control in anterior cingulate cortex during speeded decision-making. Nat Hum Behav 4, 412–422 (2020).

30. Vassena, E. et al. Overlapping neural systems represent cognitive effort and reward anticipation. PLoS One 9, (2014).

31. Kurniawan, I. T., Guitart-Masip, M., Dayan, P. & Dolan, R. J. Effort and valuation in the brain: The effects of anticipation and execution. Journal of Neuroscience 33, 6160–6169 (2013).

32. Goh, A. X. A., Bennett, D., Bode, S. & Chong, T. T. J. Neurocomputational mechanisms underlying the subjective value of information. Commun Biol 4, (2021).

33. Amiez, C., Joseph, J. P. & Procyk, E. Anterior cingulate error-related activity is modulated by predicted reward. European Journal of Neuroscience 21, 3447–3452 (2005).

34. Blanchard, T. C., Strait, C. E. & Hayden, B. Y. Ramping ensemble activity in dorsal anterior cingulate neurons during persistent commitment to a decision. J Neurophysiol 114, 2439–2449 (2015).

35. Bush, G. et al. Dorsal Anterior Cingulate Cortex: A Role in Reward-Based Decision Making. vol. 99 10.1073pnas.012470999 (2002).

36. Vassena, E., Holroyd, C. B. & Alexander, W. H. Computational models of anterior cingulate cortex: At the crossroads between prediction and effort. Front Neurosci 11, (2017).

37. Aarts, E. & Roelofs, A. Attentional Control in Anterior Cingulate Cortex Based on Probabilistic Cueing. http://mitprc.silverchair.com/jocn/article-pdf/23/3/716/1774802/jocn.2010.21435.pdf (2010).

38. Strait, C. E. et al. Neuronal selectivity for spatial positions of offers and choices in five reward regions. J Neurophysiol 115, 1098–1111 (2016).

39. Azab, H. & Hayden, B. Y. Correlates of decisional dynamics in the dorsal anterior cingulate cortex. PLoS Biol 15, (2017).

40. Azab, H. & Hayden, B. Y. Correlates of economic decisions in the dorsal and subgenual anterior cingulate cortices. European Journal of Neuroscience 47, 979–993 (2018).

41. Farashahi, S., Azab, H., Hayden, B. & Soltani, A. On the flexibility of basic risk attitudes in monkeys. Journal of Neuroscience 38, 4383–4398 (2018).

42. Blanchard, T. C. & Hayden, B. Y. Neurons in dorsal anterior cingulate cortex signal postdecisional variables in a foraging task. Journal of Neuroscience 34, 646–655 (2014).

43. Shidara, M. & Richmond, B. J. Anterior Cingulate: Single Neuronal Signals Related to Degree of Reward Expectancy. Science (1979) 296, 1709–1711 (2002).

44. Cai, X. & Padoa-Schioppa, C. Neuronal activity in dorsal anterior cingulate cortex during economic choices under variable action costs. Elife 10, (2021).

45. Hayden, B. Y., Pearson, J. M. & Platt, M. L. Fictive reward signals in the anterior cingulate cortex. Science (1979) 324, 948–950 (2009).

46. Kerns, J. G. et al. Anterior Cingulate Conflict Monitoring and Adjustments in Control. Science (1979) 303, 1023–1026 (2004).

47. Hayden, B. Y., Heilbronner, S. R., Pearson, J. M. & Platt, M. L. Surprise signals in anterior cingulate cortex: Neuronal encoding of unsigned reward prediction errors driving adjustment in behavior. Journal of Neuroscience 31, 4178–4187 (2011).

48. Bos, W. Van Den & McClure, S. M. Towards a general model of temporal discounting. J Exp Anal Behav 99, 58–73 (2013).

49. Kim, S., Hwang, J. & Lee, D. Prefrontal Coding of Temporally Discounted Values during Intertemporal Choice. Neuron 59, 161–172 (2008).

50. Louie, K. & Glimcher, P. W. Separating value from choice: Delay discounting activity in the lateral intraparietal area. Journal of Neuroscience 30, 5498–5507 (2010).

51. Blanchard, T. C., Pearson, J. M. & Hayden, B. Y. Postreward delays and systematic biases in measures of animal temporal discounting. Proc Natl Acad Sci U S A 110, 15491–15496 (2013).

52. Hayden, B. Y. Time discounting and time preference in animals: A critical review. Psychon Bull Rev 23, 39–53 (2016).

53. Tversky, A. & Kahneman, D. Judgment under Uncertainty: Heuristics and Biases Biases in judgments reveal some heuristics of thinking under uncertainty. Science (1979) 185, 1124–1131 (1974).

54. Tversky, A. & Kahneman, D. Advances in Prospect Theory: Cumulative Representation of Uncertainty. J Risk Uncertain 5, 297–323 (1992).

55. O’Doherty, J., Kringelbach, M. L., Rolls, E. T., Hornak, J. & Andrews, C. Abstract reward and punishment representations in the human orbitofrontal cortex. Nat Neurosci 4, 95–102 (2001).

56. Tobler, P. N., O’Doherty, J. P., Dolan, R. J. & Schultz, W. Reward value coding distinct from risk attitude-related uncertainty coding in human reward systems. J Neurophysiol 97, 1621–1632 (2007).

57. Lee, D., Seo, H. & Jung, M. W. Neural basis of reinforcement learning and decision making. Annual Review of Neuroscience vol. 35 287–308 Preprint at 10.1146/annurev-neuro-062111-150512 (2012).

58. Holroyd, C. B. & Yeung, N. Motivation of extended behaviors by anterior cingulate cortex. Trends in Cognitive Sciences vol. 16 122–128 Preprint at 10.1016/j.tics.2011.12.008 (2012).

59. Sallet J. et al. Expectations, gains, and losses in the anterior cingulate cortex. Cogn Affect Behav Neurosci 7, 327–336 (2007).

60. Hajnal, M. A. et al. Shifts in attention drive context-dependent subspace encoding in anterior cingulate cortex in mice during decision making. Nat Commun 15, (2024).

61. Weissman, D. H., Gopalakrishnan, A., Hazlett, C. J. & Woldorff, M. G. Dorsal anterior cingulate cortex resolves conflict from distracting stimuli by boosting attention toward relevant events. Cerebral Cortex 15, 229–237 (2005).

62. Lake, J. I. et al. Reward anticipation and punishment anticipation are instantiated in the brain via opponent mechanisms. Psychophysiology 56, (2019).

63. Aarts, E., Roelofs, A. & Van Turennout, M. Anticipatory activity in anterior cingulate cortex can be independent of conflict and error likelihood. Journal of Neuroscience 28, 4671–4678 (2008).

64. Meder, D., Herz, D. M., Rowe, J. B., Lehéricy, S. & Siebner, H. R. The role of dopamine in the brain - lessons learned from Parkinson’s disease. NeuroImage vol. 190 79–93 Preprint at 10.1016/j.neuroimage.2018.11.021 (2019).

65. Berridge, K. C. & Robinson, T. E. Parsing reward. Trends Neurosci 26, 507–513 (2003).

66. Glimcher, P. W. Understanding dopamine and reinforcement learning: The dopamine reward prediction error hypothesis. Proc Natl Acad Sci U S A 108, 15647–15654 (2011).

67. Westbrook, A. et al. Dopamine promotes cognitive effort by biasing the benefits versus costs of cognitive work. Science (1979) https://www.science.org (2020).

68. Chakroun, K. et al. Dopamine regulates decision thresholds in human reinforcement learning in males. Nat Commun 14, (2023).

69. Kennerley, S. W., Walton, M. E., Behrens, T. E. J., Buckley, M. J. & Rushworth, M. F. S. Optimal decision making and the anterior cingulate cortex. Nat Neurosci 9, 940–947 (2006).

70. Morcos, A. S. & Harvey, C. D. History-dependent variability in population dynamics during evidence accumulation in cortex. Nat Neurosci 19, 1672–1681 (2016).

71. Abrahamyan, A., Silva, L. L., Dakin, S. C., Carandini, M. & Gardner, J. L. Adaptable history biases in human perceptual decisions. Proc Natl Acad Sci U S A 113, E3548–E3557 (2016).

72. McGinty, V. B. & Lupkin, S. M. Behavioral read-out from population value signals in primate orbitofrontal cortex. Nat Neurosci 26, 2203–2212 (2023).

