## Supplementary Information for "Accumulation of virtual tokens towards a jackpot reward enhances performance and value encoding in dorsal anterior cingulate cortex"

| | No Jackpot $JPT = 0$ | Jackpot $JPT = 1$ | Low $ATC = [0, 1]$ | Medium $ATC = [2, 3]$ | High $ATC = [4, 5]$ |
|---|---|---|---|---|---|
| Subject 1 | 38606 (82.09%) | 8424 (17.91%) | 20674 (43.96%) | 15222 (32.37%) | 11134 (23.67%) |
| Subject 2 | 48910 (83.02%) | 10001 (16.98%) | 26889 (45.64%) | 19057 (32.35%) | 12965 (22.01%) |

| | No Jackpot $JPT = 0$ | Jackpot $JPT = 1$ | Low $ATC = [0, 1]$ | Medium $ATC = [2, 3]$ | High $ATC = [4, 5]$ |
|---|---|---|---|---|---|
| Subject 1 Easy | 19504 (41.57%) | 4170 (9.05%) | 10289 (22.08%) | 7388 (16.66%) | 5547 (11.88%) |
| Hard | 19552 (40.51%) | 4254 (8.87%) | 10385 (21.88%) | 7834 (15.71%) | 5587 (11.79%) |
| Subject 2 Easy | 24628 (41.22%) | 4977 (8.53%) | 13588 (22.58%) | 9500 (16.22%) | 6517 (10.95%) |
| Hard | 24282 (41.81%) | 5024 (8.45%) | 13301 (23.07%) | 9557 (16.13%) | 6448 (11.06%) |

**Supplementary Table ST1: Trials available for jackpot and accumulated token counts ranges.** Numbers of trials (fractions of the total) available for different token counts indexed by $JPT$ and $ATC$ ranges. The variable $JPT$ indicates the presence ($JPT = 1$) or absence ($JPT = 0$) of jackpot reward on previous trial. The variable $ATC$ is the accumulated tokens count as of the start of the current trial. The number of trials and fractions of the total are also shown for Easy vs Hard trials. The median value of $\Delta_{EV}$ is $median(\Delta_{EV}) = 1$ and is used to split data in 'Hard' and 'Easy' trials in behavioral analyses. These $ATC$ settings are used in behavioral analyses. In neural analyses, we use 'Low' ($ATC = [0, 1]$) and 'High' ($ATC = [2, 5]$) as this allows us to improve the trial availability per condition.

| | $\Delta_{EV} \leq 0.5$ | $0.5 < \Delta_{EV} \leq 1.5$ | $1.5 < \Delta_{EV} \leq 2.5$ | $2.5 < \Delta_{EV} \leq 3.5$ | $\Delta_{EV} > 3.5$ |
|---|---|---|---|---|---|
| Subject 1 | 13672 (29.07%) | 19412 (41.28%) | 10282 (21.86%) | 3235 (6.88%) | 429 (0.91%) |
| Subject 2 | 17152 (29.12%) | 23945 (40.65%) | 13298 (22.57%) | 3996 (6.78%) | 520 (0.88%) |

| | Hard: ($\Delta_{EV} < 1$) | Easy: ($\Delta_{EV} \geq 1$) |
|---|---|---|
| Subject 1 | 23806 (50.62%) | 23224 (49.38%) |
| Subject 2 | 29306 (49.75%) | 29605 (50.25%) |

**Supplementary Table ST2: Trials available for different difficulty levels.** Numbers of trials (fractions of the total) available for different difficulty levels indexed by $\Delta_{EV}$ ranges. Easier trials have larger $\Delta_{EV}$. The median value of $\Delta_{EV}$ is $median(\Delta_{EV}) = 1$ and is used to split data in 'Hard' and 'Easy' trials in behavioral analyses. Note that in neural analyses we use $\Delta_{SV}$ based on $SV$.

| Subject | $\beta_0$ | $\beta_{\Delta_{EV}}$ | $\beta_{M_{EV}}$ | $\beta_{ORL}$ | $\beta_{ATC}$ | $\beta_{JPT}$ | $\beta_{OPT}$ | $\beta_{TSLR}$ |
|---|---|---|---|---|---|---|---|---|
| 1 | -0.18±0.04<br>p=5.8·10$^{-51}$ | 1.79±0.03<br>p<10$^{-308}$ | 0.42±0.02<br>p=3.6·10$^{-103}$ | 0.49±0.01<br>p<10$^{-308}$ | 0.08±0.01<br>p=5.96·10$^{-11}$ | 0.19±0.06<br>p=4.4·10$^{-4}$ | 0.000018±0.01<br>p=0.99 | -0.03±0.01<br>p=7.5·10$^{-7}$ |
| 2 | -0.46±0.04<br>p=1.7·10$^{-6}$ | 1.28±0.03<br>p<10$^{-308}$ | 0.57±0.02<br>p=1.26·10$^{-274}$ | 0.48±0.01<br>p<10$^{-308}$ | 0.15±0.01<br>p=1.2·10$^{-49}$ | 0.22±0.06<br>p=9.2·10$^{-7}$ | -0.03±0.01<br>p=6.8·10$^{-3}$ | -0.01±0.01<br>p=0.0502 |

**Supplementary Table ST3:** Numerical mean ± SE and p-values for Figure 2A.

| | Subject | No Jackpot | Jackpot | Low | Medium | High |
|---|---|---|---|---|---|---|
| **Prob. Correct choice** | 1 | 79.8±0.5% | 80.5±0.6% | 79.8±0.6% | 79.8±0.6% | 81.5±0.6% |
| | 2 | 74.7±0.4% | 72.2±0.5% | 72.1±0.5% | 74.5±0.5% | 78.6±0.4% |
| **Execution time (s)** | 1 | 2.12±0.06 | 2.27±0.11 | 2.31±0.10 | 2.03±0.05 | 2.01±0.07 |
| | 2 | 2.00±0.05 | 2.17±0.11 | 2.26±0.10 | 1.88±0.03 | 1.73±0.01 |

**Supplementary Table ST4:** Numerical mean ± s.e.m values for Figure 2B-C.

| Subject | No Jackpot | Jackpot | Low | Medium | High |
|---|---|---|---|---|---|
| **1 Easy** | 08.5±0.5% | 08.0±0.6% | 09.3±0.6% | 07.9±0.6% | 07.3±0.5% |
| **1 Hard** | 31.1±0.5% | 30.2±0.8% | 31.6±0.6% | 31.4±0.8% | 28.9±0.8% |
| **2 Easy** | 14.2±0.5% | 19.5±0.7% | 18.1±0.6% | 14.7±0.7% | 09.3±0.4% |
| **2 Hard** | 35.9±0.4% | 35.5±0.7% | 37.3±0.5% | 35.7±0.7% | 33.4±0.6% |
| | $\Delta_{EV} = 0$ | $\Delta_{EV} = 1$ | $\Delta_{EV} = 2$ | $\Delta_{EV} = 3$ | $\Delta_{EV} = 4$ |
| **1 Low** | 38.6±0.7% | 21.2±0.7% | 07.6±0.6% | 03.6±0.6% | 01.3±0.8% |
| **1 High** | 36.7±1.0% | 18.4±0.7% | 04.8±0.6% | 02.3±0.5% | 00.0±0.0% |
| **2 Low** | 41.2±0.7% | 30.2±0.6% | 16.7±0.8% | 10.3±0.8% | 03.0±1.0% |
| **2 High** | 38.7±1.0% | 23.6±0.7% | 7.00±0.6% | 02.3±0.4% | 01.1±0.8% |

| | No J. vs. J. | L. vs. M. | L. vs. H. | M. vs. H. |
|---|---|---|---|---|
| **1 Easy** | p=0.067 | $p=1.51\cdot10^{-4}$ | $p=1.51\cdot10^{-4}$ | p=0.18 |
| **1 Hard** | p=0.24 | p=0.91 | $p=6.04\cdot10^{-3}$ | p=0.01 |
| **2 Easy** | $p=1.21\cdot10^{-10}$ | $p=1.16\cdot10^{-7}$ | $p=1.29\cdot10^{-19}$ | $p=3.54\cdot10^{-15}$ |
| **2 Hard** | p=0.31 | p=0.03 | $p=1.42\cdot10^{-6}$ | p=0.02 |

| | | | **Low vs High** | | |
|---|---|---|---|---|---|
| | $\Delta_{EV} = 0$ | $\Delta_{EV} = 1$ | $\Delta_{EV} = 2$ | $\Delta_{EV} = 3$ | $\Delta_{EV} = 4$ |
| **1 Low vs High** | p=0.12 | p=0.022 | $p=2.76\cdot10^{-4}$ | p=0.015 | p=0.14 |
| **2 Low vs High** | p=0.039 | $p=1.74\cdot10^{-10}$ | $p=8.93\cdot10^{-19}$ | $p=7.83\cdot10^{-17}$ | p=0.035 |

**Supplementary Tabe ST5:** Numerical mean ± s.e.m and p-values for Figure 3.

| | Subject | No Jackpot | Jackpot | Low | Medium | High |
|---|---|---|---|---|---|---|
| **Faction risky choices** | 1 Easy | 51.0±0.4% | 49.4±0.8% | 51.5±0.5% | 49.8±0.6% | 50.3±0.7% |
| | 1 Hard | 68.8±0.3% | 66.7±0.7% | 68.1±0.5% | 69.4±0.5% | 67.7±0.6% |
| | 2 Easy | 54.5±0.3% | 56.4±0.7% | 56.5±0.4% | 55.9±0.5% | 49.8±0.6% |
| | 2 Hard | 70.2±0.3% | 67.8±0.7% | 69.3±0.4% | 72.4±0.5% | 67.0±0.6% |
| $\boldsymbol{\theta = -\beta_2/\beta_1}$ | 1 Easy | -0.23±0.00706 | -0.21±0.01507 | -0.24±0.00953 | -0.22±0.01134 | -0.19±0.01333 |
| | 1 Hard | -0.25±0.00002 | -0.21±0.00131 | -0.24±0.00003 | -0.25±0.00006 | -0.22±0.00077 |
| | 2 Easy | -0.33±0.00581 | -0.30±0.01387 | -0.38±0.00846 | -0.39±0.00809 | -0.23±0.01146 |
| | 2 Hard | -0.33±0.00197 | -0.39±0.00380 | -0.34±0.00394 | -0.37±0.00398 | -0.24±0.00033 |
| $\boldsymbol{\beta_1}$ | 1 Easy | 1.68±0.04 | 1.77±0.08 | 1.62±0.05 | 1.74±0.06 | 1.80±0.07 |
| | 1 Hard | 2.30±0.06 | 2.38±0.14 | 2.17±0.09 | 2.41±0.11 | 2.47±0.12 |
| | 2 Easy | 1.31±0.02 | 1.04±0.04 | 1.07±0.03 | 1.38±0.04 | 1.69±0.06 |
| | 2 Hard | 1.68±0.05 | 1.54±0.11 | 1.44±0.07 | 1.81±0.09 | 1.93±0.10 |
| $\boldsymbol{\beta_2}$ | 1 Easy | 0.38±0.02 | 0.37±0.04 | 0.39±0.03 | 0.39±0.03 | 0.34±0.04 |
| | 1 Hard | 0.57±0.02 | 0.49±0.03 | 0.52±0.02 | 0.61±0.03 | 0.56±0.03 |
| | 2 Easy | 0.44±0.02 | 0.40±0.03 | 0.40±0.02 | 0.54±0.03 | 0.39±0.03 |
| | 2 Hard | 0.55±0.01 | 0.47±0.03 | 0.49±0.02 | 0.66±0.02 | 0.47±0.02 |

**Supplementary Table ST6:** Numerical mean ± s.e.m. for the Fraction of risky choices and mean ± CI values for $\theta, \beta_1, \beta_2$ shown in Figure 4.

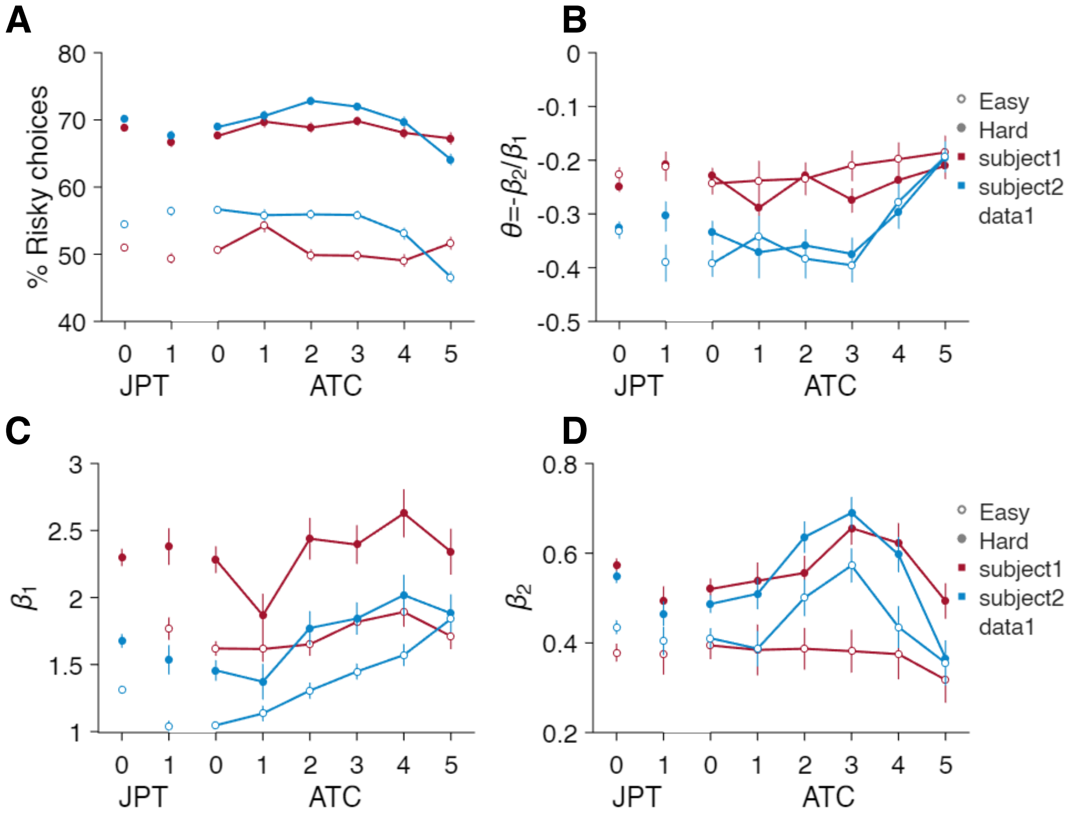


**Supplementary Figure S1. Risk seeking attitude vs accumulated reward and difficulty.**

**A)** Fraction of trials (mean ± s.e.m.) with choice for the offer with higher risk. The data are split in 'No Jackpot' ($JPT = 0$), 'Jackpot' ($JPT = 1$), and for accumulated tokens count ($ATC = 0 - 5$) for subject 1 (red) and subject 2 (blue), in Easy ($\Delta_{EV} \geq 1$, empty markers) and Hard trials ($\Delta_{EV} < 1$, filled markers). **B)** Markowitz risk return model for the offer utility based on the mean value ($EV$) and risk ($R$) of the offers. The model parameter ($\theta$) describes risk attitude ($\theta < 0$ risk seeking, $\theta > 0$ risk avoiding) for jackpot cases and for values of accumulated token counts. Data is split in difficulty and across subjects as in A. **C)** Markowitz risk return model, parameter $\beta_1$ relative to $EV$ difference weights for jackpot cases and for values of accumulated token counts. Data is split as in A. **D)** Markowitz risk return model, parameter $\beta_2$ relative to risk $R$ difference weights for jackpot cases and for values of accumulated token counts. Data is split as in A.

| Subject | Model | Parameters |
|---|---|---|
| 1 | Linear (no ATC int.) | $\beta_0$:−0.30±0.03$^{***}$; $\beta_1$:2.23±0.06$^{***}$; $\beta_2$:0.55±0.02$^{***}$; $\beta_3$:0.03±0.01$^{*}$<br>$p=8.51\cdot10^{-15}$; $p=1.74\cdot10^{-19}$; $p=1.74\cdot10^{-19}$; $p=2.15\cdot10^{-2}$ |
| 1 | Linear (ATC int.) | $\beta_0$:−0.24±0.02$^{***}$; $\beta_1$:2.08±0.07$^{***}$; $\beta_2$:0.51±0.02$^{***}$; $\beta_3$:0.12±0.02$^{***}$;<br>$\beta_4$:0.07±0.01$^{***}$; $\beta_5$:−0.01±0.00$^{***}$<br>$p=5.99\cdot10^{-18}$; $p=1.74\cdot10^{-19}$; $p=1.74\cdot10^{-19}$; $p=2.73\cdot10^{-10}$;<br>$p=2.74\cdot10^{-6}$; $p=1.17\cdot10^{-4}$ |
| 1 | RDV | $\kappa_0$:2.61±0.25$^{***}$; $\kappa_1$:3.28±0.05$^{***}$; $\gamma_0$:0.79±0.02$^{***}$; $\gamma_1$:0.02±0.00$^{***}$;<br>$\lambda_0$:0.49±0.03$^{***}$; $\lambda_1$:0.18±0.02$^{***}$; $\lambda_2$:−0.02±0.00$^{***}$;<br>$\beta_0$:−0.25±0.02$^{***}$; $\beta_1$:3.37±0.08$^{***}$; $\beta_2$:0.72±0.04$^{***}$<br>$p=1.74\cdot10^{-19}$; $p=1.74\cdot10^{-19}$; $p=1.74\cdot10^{-19}$; $p=1.74\cdot10^{-19}$;<br>$p=1.74\cdot10^{-19}$; $p=1.74\cdot10^{-19}$; $p=3.23\cdot10^{-5}$;<br>$p=5.80\cdot10^{-18}$; $p=1.74\cdot10^{-19}$; $p=1.74\cdot10^{-19}$ |
| 2 | Linear (no ATC int.) | $\beta_0$:−0.45±0.04$^{***}$; $\beta_1$:1.50±0.04$^{***}$; $\beta_2$:0.53±0.02$^{***}$; $\beta_3$:0.01±0.01<br>$p=9.25\cdot10^{-18}$; $p=9.02\cdot10^{-21}$; $p=9.02\cdot10^{-21}$; $p=2.54\cdot10^{-1}$ |
| 2 | Linear (ATC int.) | $\beta_0$:−0.44±0.03$^{***}$; $\beta_1$:1.25±0.04$^{***}$; $\beta_2$:0.48±0.02$^{***}$; $\beta_3$:0.19±0.02$^{***}$;<br>$\beta_4$:0.17±0.01$^{***}$; $\beta_5$:−0.04±0.00$^{***}$<br>$p=1.41\cdot10^{-20}$; $p=9.02\cdot10^{-21}$; $p=9.02\cdot10^{-21}$; $p=1.74\cdot10^{-19}$;<br>$p=1.96\cdot10^{-17}$; $p=1.68\cdot10^{-16}$ |
| 2 | RDV | $\kappa_0$:0.97±0.12$^{***}$; $\kappa_1$:3.09±0.06$^{***}$; $\gamma_0$:0.66±0.02$^{***}$; $\gamma_1$:0.03±0.00$^{***}$;<br>$\lambda_0$:0.27±0.02$^{***}$; $\lambda_1$:0.16±0.02$^{***}$; $\lambda_2$:−0.01±0.00$^{*}$;<br>$\beta_0$:−0.47±0.03$^{***}$; $\beta_1$:3.28±0.06$^{***}$; $\beta_2$:0.64±0.05$^{***}$<br>$p=9.02\cdot10^{-21}$; $p=9.02\cdot10^{-21}$; $p=9.02\cdot10^{-21}$; $p=9.02\cdot10^{-21}$;<br>$p=9.02\cdot10^{-21}$; $p=9.02\cdot10^{-21}$; $p=3.53\cdot10^{-2}$;<br>$p=1.48\cdot10^{-20}$; $p=9.02\cdot10^{-21}$; $p=2.06\cdot10^{-16}$ |
| 1,2 | Linear (no ATC int.) | $\beta_0$:−0.37±0.02$^{***}$; $\beta_1$:1.85±0.04$^{***}$; $\beta_2$:0.54±0.01$^{***}$; $\beta_3$:0.02±0.01$^{*}$<br>$p=1.46\cdot10^{-30}$; $p=2.95\cdot10^{-38}$; $p=2.95\cdot10^{-38}$; $p=1.56\cdot10^{-2}$ |
| 1,2 | Linear (ATC int.) | $\beta_0$:−0.35±0.02$^{***}$; $\beta_1$:1.65±0.05$^{***}$; $\beta_2$:0.49±0.01$^{***}$; $\beta_3$:0.15±0.01$^{***}$;<br>$\beta_4$:0.12±0.01$^{***}$; $\beta_5$:−0.02±0.00$^{***}$<br>$p=7.40\cdot10^{-37}$; $p=2.95\cdot10^{-38}$; $p=2.95\cdot10^{-38}$; $p=1.59\cdot10^{-28}$;<br>$p=1.06\cdot10^{-21}$; $p=1.74\cdot10^{-19}$ |
| 1,2 | RDV | $\kappa_0$:1.76±0.14$^{***}$; $\kappa_1$:3.18±0.04$^{***}$; $\gamma_0$:0.72±0.01$^{***}$; $\gamma_1$:0.03±0.00$^{***}$;<br>$\lambda_0$:0.38±0.02$^{***}$; $\lambda_1$:0.17±0.01$^{***}$; $\lambda_2$:−0.01±0.00$^{***}$;<br>$\beta_0$:−0.37±0.02$^{***}$; $\beta_1$:3.33±0.05$^{***}$; $\beta_2$:0.68±0.03$^{***}$<br>$p=2.95\cdot10^{-38}$; $p=2.95\cdot10^{-38}$; $p=2.95\cdot10^{-38}$; $p=2.95\cdot10^{-38}$;<br>$p=2.95\cdot10^{-38}$; $p=2.95\cdot10^{-38}$; $p=7.57\cdot10^{-6}$;<br>$p=7.88\cdot10^{-37}$; $p=2.95\cdot10^{-38}$; $p=4.13\cdot10^{-34}$ |

**Supplementary Table ST7.** Parameter estimates (mean ± s.e.m) across sessions and $k = 4$ cross-validation folds for the models tested. Subjects are combined by pooling session by session results ($n = 109$ in subject 1, 118 in subject 2). All parameters show significant magnitude (***p<0.001, *p<0.05) as assessed via two-sided signed-rank tests, FDR corrected via Benjamini-Hochberg procedure.

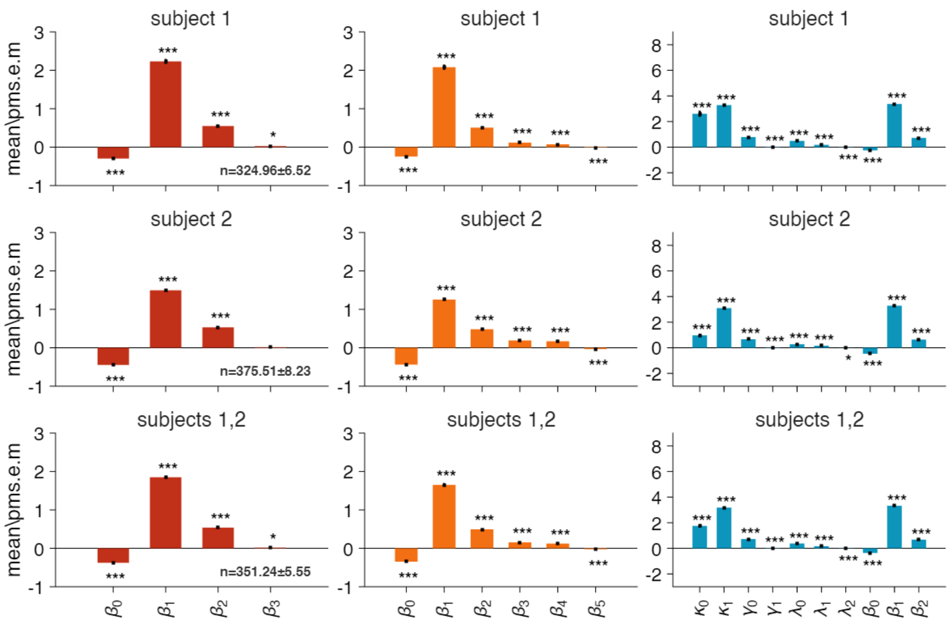


**Supplementary Figure S2.** Parameter estimates choice models. Left: linear model with no ATC interactions, middle: linear model with ATC interactions, right: RDV model. The estimations are run session by session and then averaged over $k = 4$ folds cross-validation stages ($n$ values at the bottom right of the first column refer to mean ± s.e.m. number of trials per session used in cross-validation training sets). P-values computed across sessions (*** p<0.001, * p<0.05) via signed-rank tests, pooled and FDR corrected via Benjamini-Hochberg procedure. Results for subjects combined are combined across all sessions ($n = 109$ in subject 1, 118 in subject 2).

| Subject | Model | $n_{\hat{\Theta}}$ | Accuracy (%) | $R^2_{adj}$ | -log(L) | AIC | BIC |
|---|---|---|---|---|---|---|---|
| 1 | Linear (no ATC int.) | 4 | 92.15 ± 0.32 | 0.69 ± 0.01 | 176.6 ± 6.11 | 361.3 ± 12.22 | 380.3 ± 12.22 |
| 1 | Linear (w. ATC int.) | 6 | 92.12 ± 0.32 | 0.69 ± 0.01 | 178.3 ± 6.09 | 368.7 ± 12.18 | 397.2 ± 12.18 |
| **1** | **RDV** | **10** | **92.26 ± 0.32** | **0.70 ± 0.01** | **148.9 ± 4.86** | **317.8 ± 9.72** | **365.2 ± 9.72** |
| 2 | Linear (no ATC int.) | 4 | 88.81 ± 0.34 | 0.58 ± 0.01 | 226.5 ± 6.15 | 460.9 ± 12.29 | 479.9 ± 12.29 |
| 2 | Linear (w. ATC int.) | 6 | 88.84 ± 0.33 | 0.59 ± 0.01 | 224.5 ± 6.07 | 461.0 ± 12.14 | 489.5 ± 12.14 |
| **2** | **RDV** | **10** | **89.27 ± 0.33** | **0.61 ± 0.01** | **195.2 ± 5.27** | **410.4 ± 10.53** | **457.8 ± 10.53** |
| 1, 2 | Linear (no ATC int.) | 4 | 90.41 ± 0.26 | 0.64 ± 0.01 | 202.5 ± 4.63 | 413.1 ± 9.27 | 432.0 ± 9.27 |
| 1,2 | Linear (w. ATC int.) | 6 | 90.41 ± 0.25 | 0.63 ± 0.01 | 202.3 ± 4.56 | 416.7 ± 9.12 | 445.1 ± 9.12 |
| **1, 2** | **RDV** | **10** | **90.71 ± 0.25** | **0.65 ± 0.01** | **173.0 ± 3.91** | **365.9 ± 7.81** | **413.4 ± 7.81** |

**Supplementary Table ST8.** Performances for model comparisons (mean ± s.e.m across sessions). In $n_{\hat{\Theta}}$ we show the number of parameters fit in each model. Subjects are combined be pooling session by session results ($n = 109$ in subject 1, 118 in subject 2).

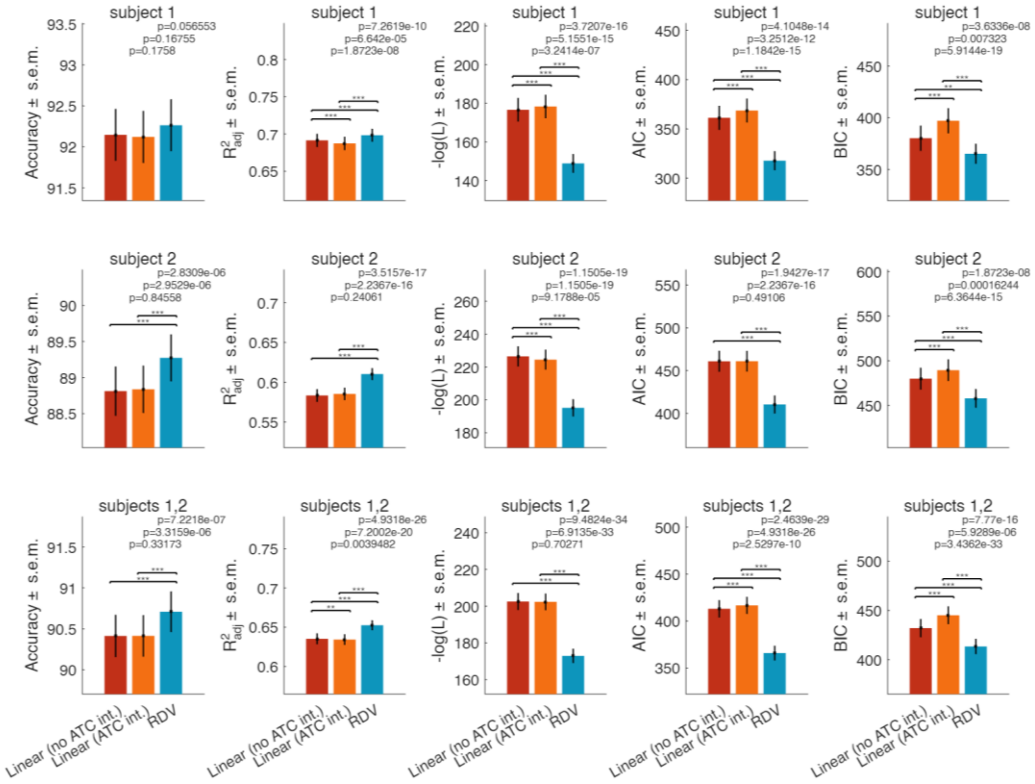


**Supplementary Figure S3.** Comparison of qualitative parameters for the three choice models. Linear no ATC interaction (red), Linear with ATC interaction (orange), and RDV (blue). Accuracy is computed as fraction of choice prediction $\hat{p}_i$ matching test set choices $y_i = 1$ if first offer is chosen and 0 otherwise. The predictions $\hat{p}_i$ are obtained by combining test data $[v^t_{1,i}, v^b_{1,i}, p^t_{1,i}, v^t_{2,i}, v^b_{2,i}, p^t_{2,i}, ATC_i]$ and $\widehat{\Theta}$ fits on training data, with $\widehat{\Theta} = [\hat{\beta}_0, \hat{\beta}_1, \hat{\beta}_2, \hat{\beta}_3]$ in 'linear no ATC interactions', $\widehat{\Theta} = [\hat{\beta}_0, \hat{\beta}_1, \hat{\beta}_2, \hat{\beta}_3, \hat{\beta}_4, \hat{\beta}_5]$ in 'linear with ATC interactions', $\widehat{\Theta} = [\hat{\kappa}_0, \hat{\kappa}_1, \hat{\gamma}_0, \hat{\gamma}_1, \hat{\lambda}_0, \hat{\lambda}_1, \hat{\lambda}_2, \hat{\beta}_0, \hat{\beta}_1, \hat{\beta}_2]$ in 'RDV'. The coefficient of determination is $R^2_{adj} = 1 - (\Sigma_i (y_i - \hat{p}_i)^2 / \Sigma_i (y_i - \bar{y}_i)^2)(n-1)/(n - n_{\widehat{\Theta}} - 1)$, with $n$ the number of total trials, and $n_{\widehat{\Theta}}$ the number of parameters fit. The negative log likelihood $-\log L(\widehat{\Theta}|y_i)$ includes Ridge penalty and is used to compute the Akaike Information Criteria $AIC = 2n_{\widehat{\Theta}} + 2\left(-\log(L) + \lambda_{\text{ridge}}\left|\widehat{\Theta}_{\text{ridge}}\right|^2\right)$, and the Bayesian Information Criteria $BIC = \log(n)\, n_{\widehat{\Theta}} + 2\left(-\log(L) + \lambda_{\text{ridge}}\left|\widehat{\Theta}_{\text{ridge}}\right|^2\right)$. All metrics consistently indicate the reference-dependent model as significantly better than other linear models for choice prediction, with improvement in both subjects (***p<0.001, **p<0.01, signed-rank assessment on session-by-session results, FDR corrected via Benjamini-Hochberg procedure; comparisons in subject 1 showed p<0.05 in Accuracy prior to FDR correction).

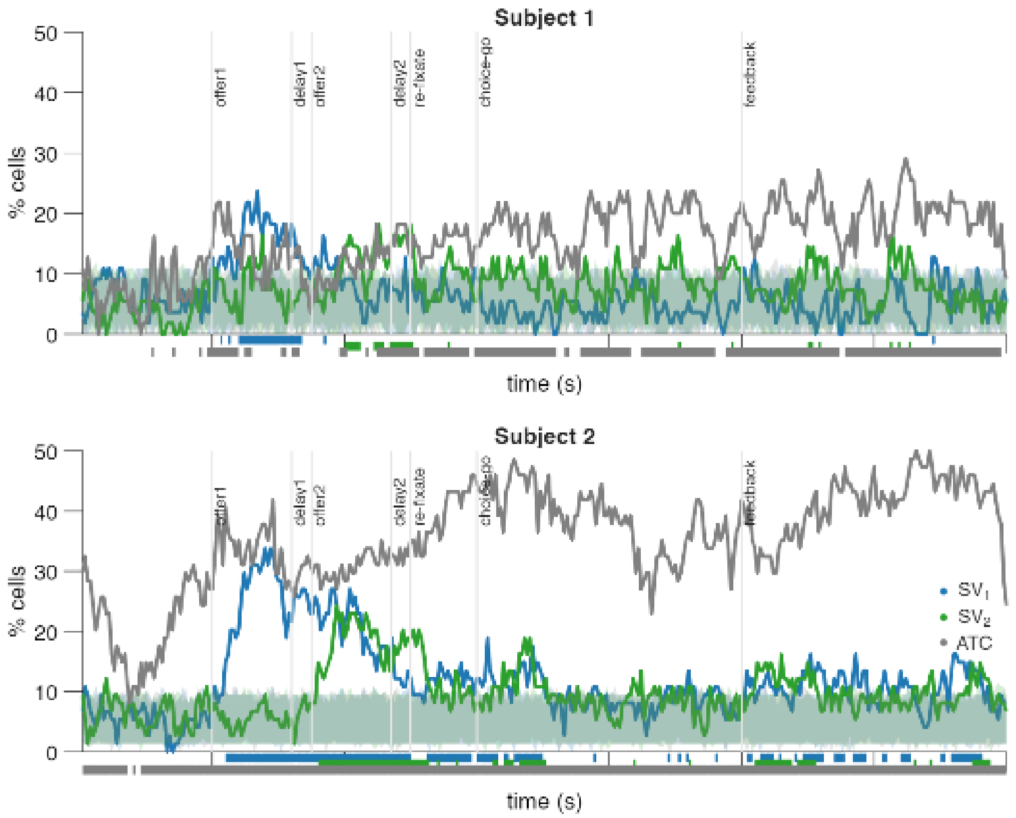


**Supplementary Figure S4.** Same as Fig. 5A, for subject 1 (top, n=55 cells) and subject 2 (bottom, n=74). The shaded areas show the 5th to 95th percentile of equivalent results from n=1000 trial order shuffled data.

| | **Pre-offer1** | **Offer 1** | **Delay 1** | **Offer 2** | **Delay 2** | **Re-fixate** | **Choice-go** | **Feedback** |
|---|---|---|---|---|---|---|---|---|
| $SV_1$ | p=1 | $p=3.99\cdot10^{-6}$ | $p=5.86\cdot10^{-3}$ | $p=3.99\cdot10^{-6}$ | $p=5.86\cdot10^{-3}$ | $p=5.92\cdot10^{-4}$ | p=1 | $p=2.88\cdot10^{-1}$ |
| $SV_2$ | p=1 | p=1 | p=1 | $p=3.99\cdot10^{-6}$ | $p=5.86\cdot10^{-3}$ | $p=2.12\cdot10^{-3}$ | $p=6.14\cdot10^{-3}$ | $p=9.66\cdot10^{-3}$ |
| $ATC$ | $p=1.62\cdot10^{-8}$ | $p=3.99\cdot10^{-6}$ | $p=5.86\cdot10^{-3}$ | $p=3.99\cdot10^{-6}$ | $P=5.86\cdot10^{-3}$ | $p=1.65\cdot10^{-5}$ | $p=2.26\cdot10^{-17}$ | $p=2.26\cdot10^{-17}$ |

**Supplementary Table ST9.** P-values in Figure 5B.

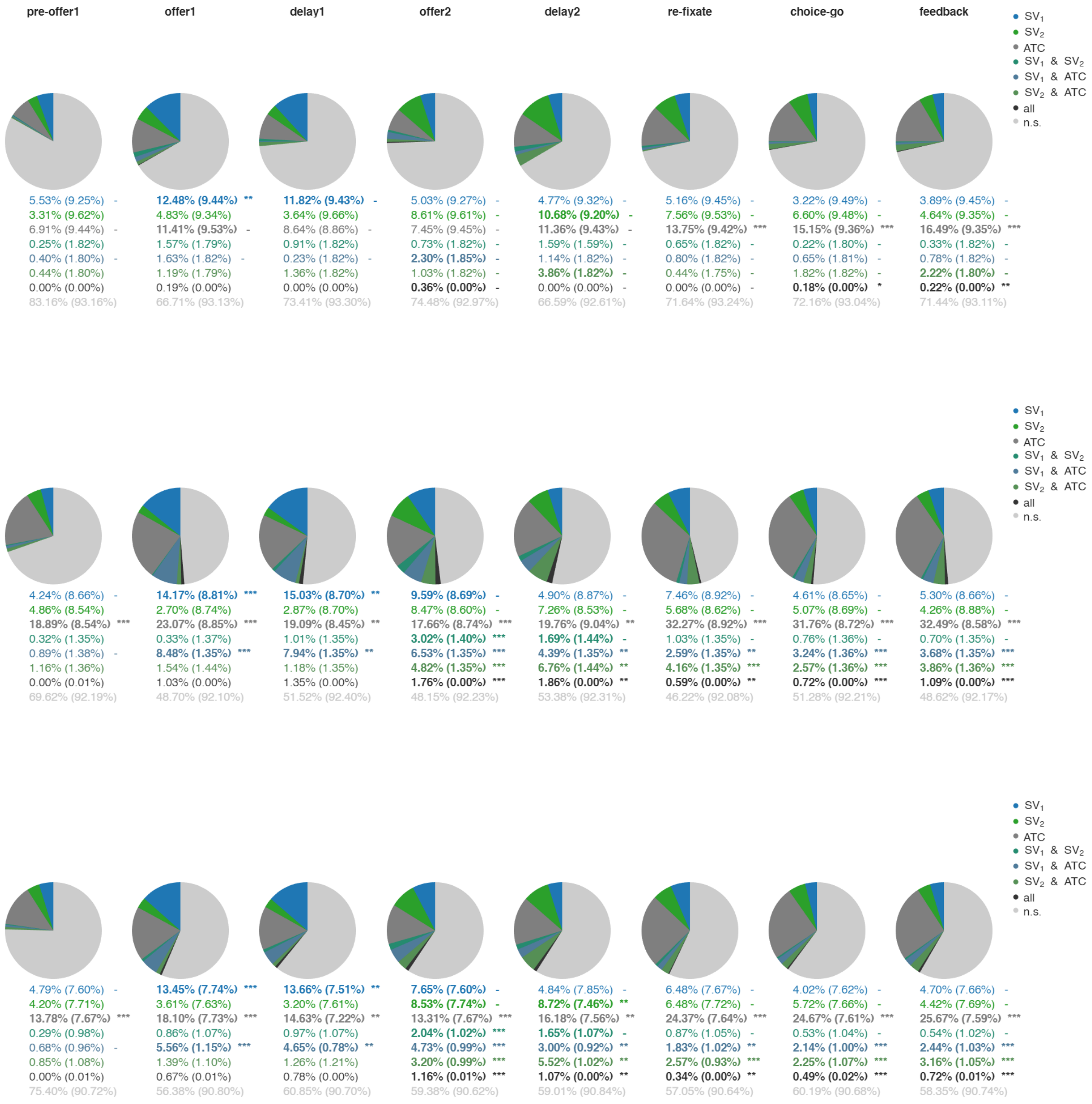


**Supplementary Figure S5.** Same as Fig. 5C, showing numerical fractions of significant cells and for data in the respective time epochs from subject 1 (top, n=55 cells), subject 2 (middle, n=74), subjects 1,2 (bottom, n=129). Fractions are compared to a significance threshold defined by the 95th percentile of equivalent fractions from n=1000 trial-order shuffles shown in parentheses. Fractions exceeding threshold are shown in bold. Significance is tested via one-tailed signed rank tests FDR corrected assessing that empirical fractions exceed significance threshold (*$p<0.05$, **$p<0.01$, ***$p<0.001$).

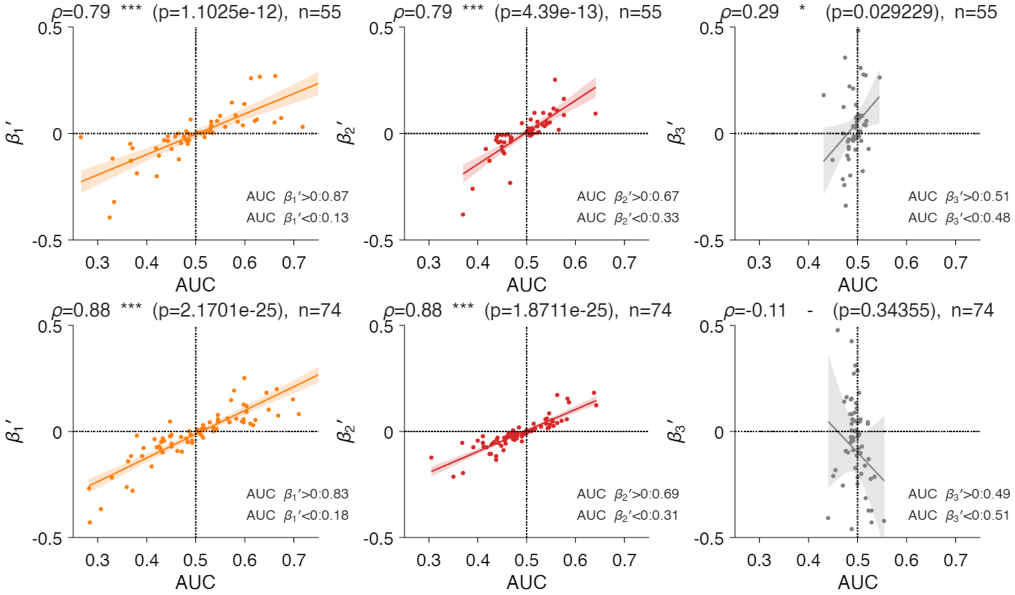


**Supplementary Figure S6.** Same as Fig. 5D, for subject 1 (top) and subject 2 (bottom).

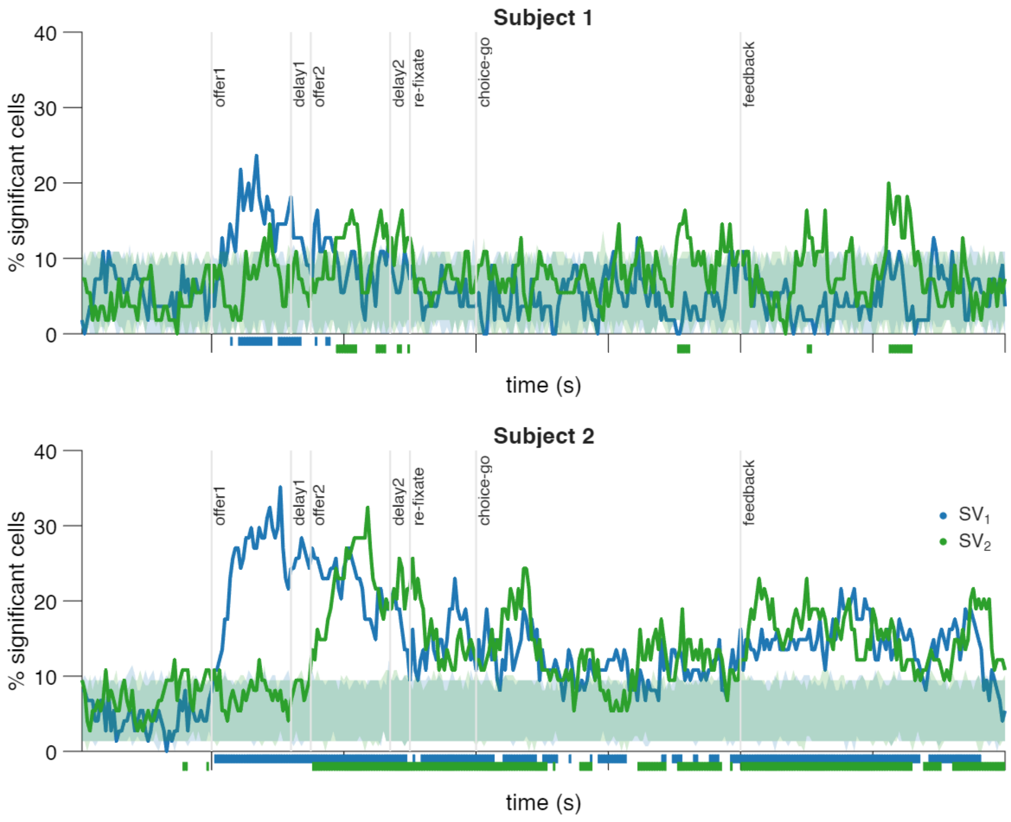


**Supplementary Figure S7.** Same as Fig. 6A, for subject 1 (top, n=55 cells) and subject 2 (bottom, n=74). The shaded areas show the 5th to 95th percentile of equivalent results from n=1000 trial order shuffled data.

| | **Pre-offer1** | **Offer 1** | **Delay 1** | **Offer 2** | **Delay 2** | **Re-fixate** | **Choice-go** | **Feedback** |
|---|---|---|---|---|---|---|---|---|
| $SV_1$ | p=1 | $p=3.93\cdot10^{-6}$ | $p=5.68\cdot10^{-3}$ | $p=2.79\cdot10^{-6}$ | $p=5.68\cdot10^{-3}$ | $p=7.47\cdot10^{-5}$ | p=0.32 | $p=7.68\cdot10^{-12}$ |
| $SV_2$ | p=1 | p=1 | p=0.99 | $p=2.79\cdot10^{-6}$ | $p=5.68\cdot10^{-3}$ | $p=1.40\cdot10^{-5}$ | $p=4.28\cdot10^{-11}$ | $p=1.10\cdot10^{-16}$ |

**Supplementary Table ST10.** P-values in Figure 6B.

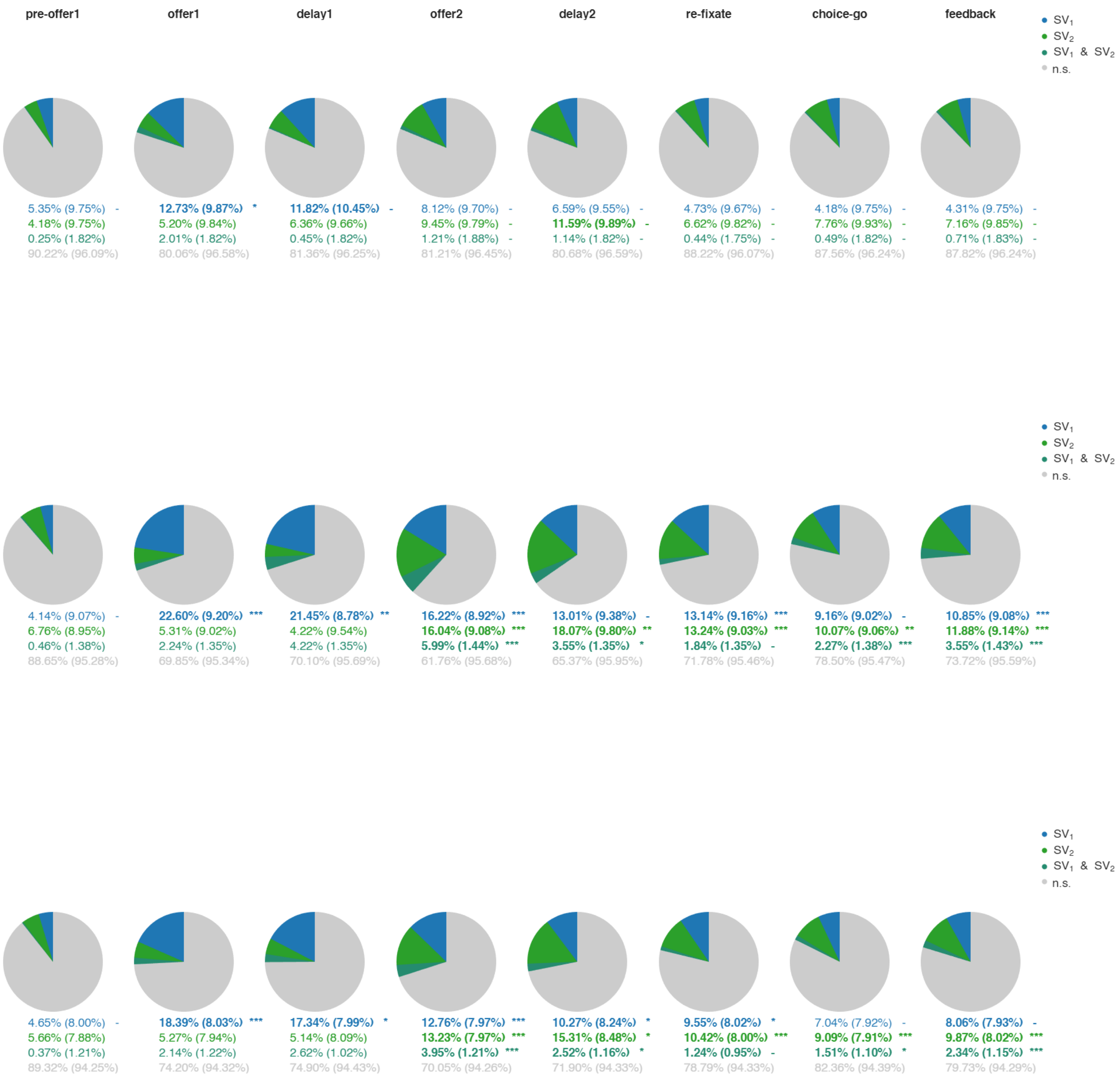


**Supplementary Figure S8.** Same as Fig. 6C showing numerical fractions of significant cells and for data in the respective time epochs from subject 1 (top, n=55 cells), subject 2 (middle, n=74), subjects 1,2 (bottom, n=129). Fractions are compared to a significance threshold defined by the 95th percentile of equivalent fractions from n=1000 trial-order shuffles shown in parentheses. Fractions exceeding threshold are shown in bold. Significance is tested via one-tailed signed rank tests FDR corrected assessing that empirical fractions exceed significance threshold (*p<0.05, **p<0.01, ***p<0.001).

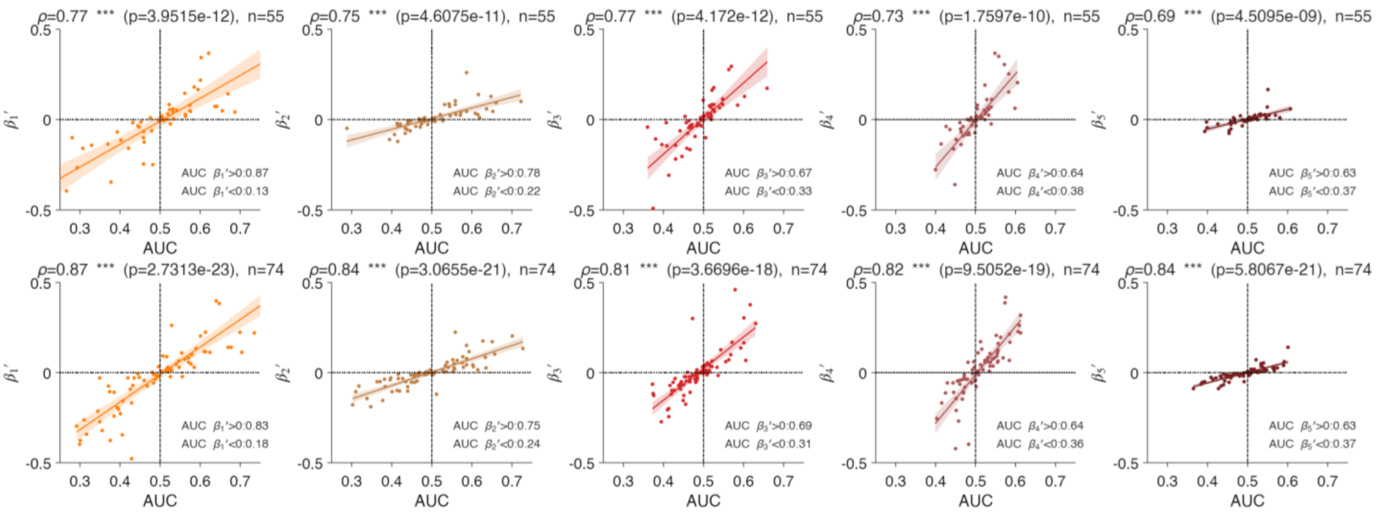


**Supplementary Figure S9.** Same as Fig. 6D, for subject 1 (top) and subject 2 (bottom).

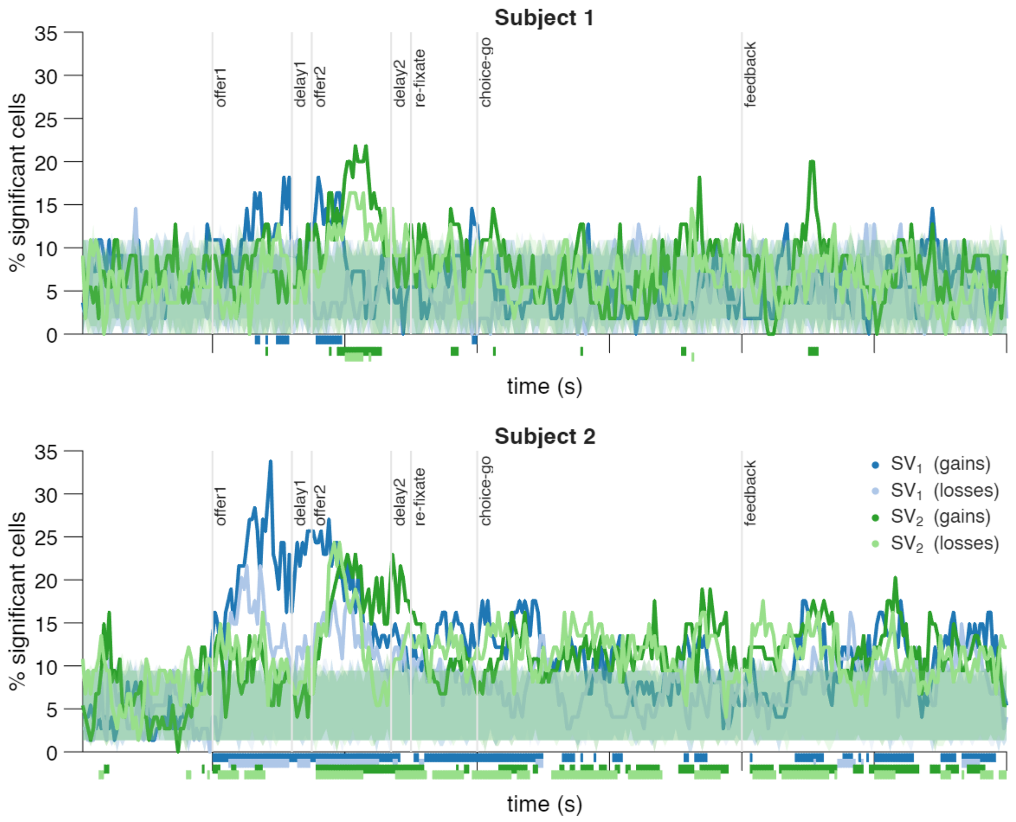


**Supplementary Figure S10.** Same as Fig. 7A, for subject 1 (top, n=55 cells) and subject 2 (bottom, n=74). The shaded areas show the 5th to 95th percentile of equivalent results from n=1000 trial order shuffled data.

| | **Pre-offer1** | **Offer 1** | **Delay 1** | **Offer 2** | **Delay 2** | **Re-fixate** | **Choice-go** | **Feedback** |
|---|---|---|---|---|---|---|---|---|
| $SV_1^+$ | p=1 | $p=7.94\cdot10^{-6}$ | p=0.0081 | $p=9.74\cdot10^{-6}$ | p=0.53 | $p=8.52\cdot10^{-5}$ | p=1 | p=0.057 |
| $SV_1^-$ | p=1 | p=0.004 | p=0.054 | p=0.0024 | p=0.054 | p=1 | p=1 | p=1 |
| $SV_2^+$ | p=1 | p=0.31 | p=1 | $p=7.14\cdot10^{-6}$ | p=0.0081 | p=0.0049 | $p=8.89\cdot10^{-9}$ | $p=1.54\cdot10^{-8}$ |
| $SV_2^-$ | p=1 | p=0.054 | p=1 | $p=2.29\cdot10^{-5}$ | p=0.0081 | p=0.001 | $p=3.76\cdot10^{-7}$ | p=0.055 |
| $SV_1^+$ vs $SV_1^-$ | p=0.045 | $p=7.94\cdot10^{-6}$ | p=0.0081 | $p=2.67\cdot10^{-5}$ | p=1 | p=0.00035 | $p=2.60\cdot10^{-8}$ | $p=1.16\cdot10^{-5}$ |
| $SV_2^+$ vs $SV_2^-$ | p=1 | p=1 | p=0.56 | p=0.00022 | p=0.031 | p=0.86 | p=0.12 | $p=5.01\cdot10^{-6}$ |

**Supplementary Table ST11.** P-values in Figure 7B.

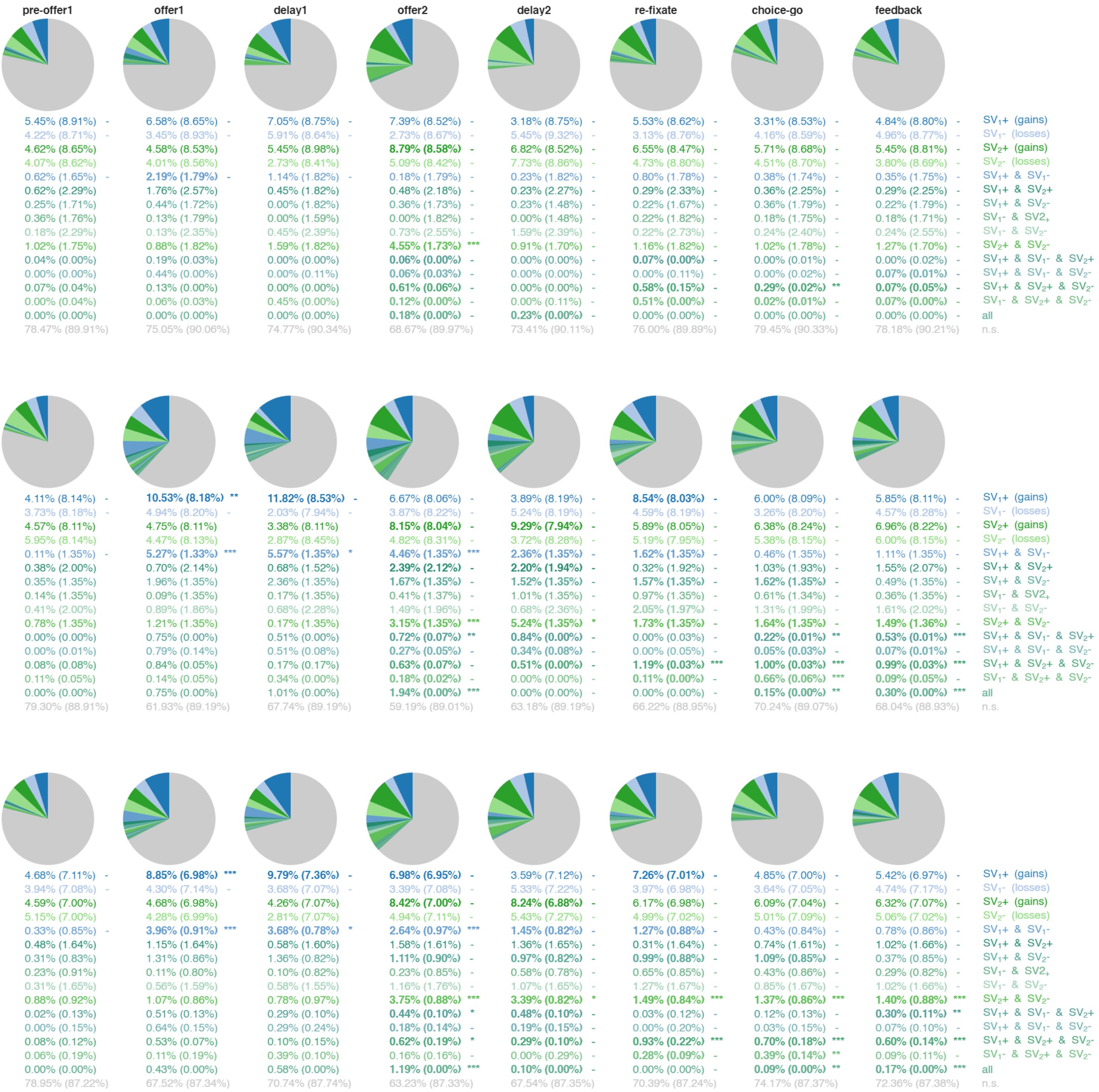


**Supplementary Figure S11.** Same as Fig. 7C showing numerical fractions of significant cells and for data in the respective time epochs from subject 1 (top, n=55 cells), subject 2 (middle, n=74), subjects 1,2 (bottom, n=129). Fractions are compared to a significance threshold defined by the

95th percentile of equivalent fractions from n=1000 trial-order shuffles shown in parentheses. Fractions exceeding threshold are shown in bold. Significance is tested via one-tailed signed rank tests FDR corrected assessing that empirical fractions exceed significance threshold (*p<0.05, **p<0.01, ***p<0.001).

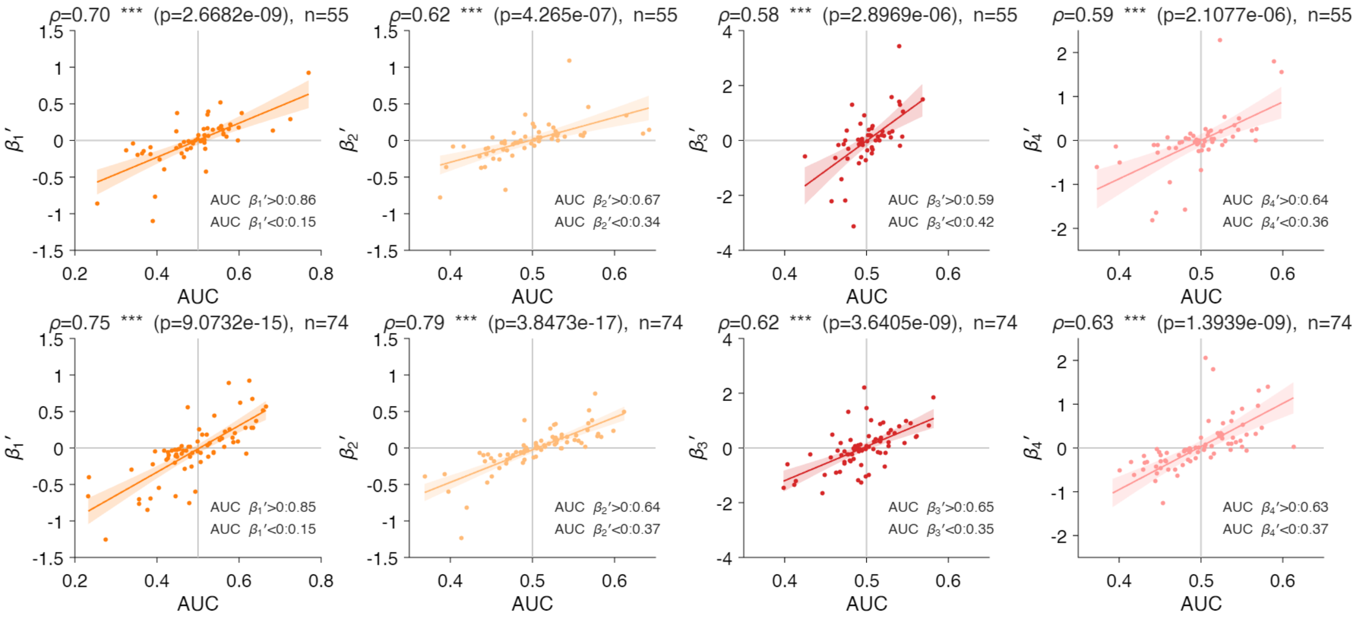


**Supplementary Figure S12.** Same as Fig. 6D, for subject 1 (top) and subject 2 (bottom).

| | $v_1^t$ | $v_1^b$ | $v_2^t$ | $v_2^b$ |
|---|---|---|---|---|
| **gains** | 216.78±4.54 (43.47%) | 213.97±4.32 (42.95%) | 213.37±4.59 (42.72%) | 214.72±4.35 (43.12%) |
| **losses** | 285.03±6.61 (56.53%) | 287.84±6.55 (57.05%) | 288.43±6.50 (57.28%) | 287.09±6.64 (56.88%) |
| **total** | 501.81±9.27 (100%) | 501.81±9.27 (100%) | 501.81±9.27 (100%) | 501.81±9.27 (100%) |

**Supplementary Table ST12.** Number of trials where offer token values resulted in gains or losses according to the 'RDV' utility function (Methods 2.4.3, 3.1.3), mean ± s.e.m. across sessions (percentage of total available trials).

| | $v_1^b$ gains/losses | $v_2^t$ gains/losses | $v_2^b$ gains/losses |
|---|---|---|---|
| $v_1^t$ gains/losses | -0.48±0.006 (100%) | 0.01±0.004 (9.30%) | 0.02±0.004 (12.40%) |
| $v_1^b$ gains/losses | | 0.01±0.004 (10.1%) | 0.01±0.004 (5.43%) |
| $v_2^t$ gains/losses | | | -0.48±0.006 (100%) |

**Supplementary Table ST13.** Pearson's correlation between binary flags indicating whether offer token values resulted in gains or losses according to the 'RDV' utility function (Methods 2.4.3, 3.1.3), mean ± s.e.m. across sessions (percentage of sessions where correlations showed significant, p<0.05 assessed via two-tailed t-tests).

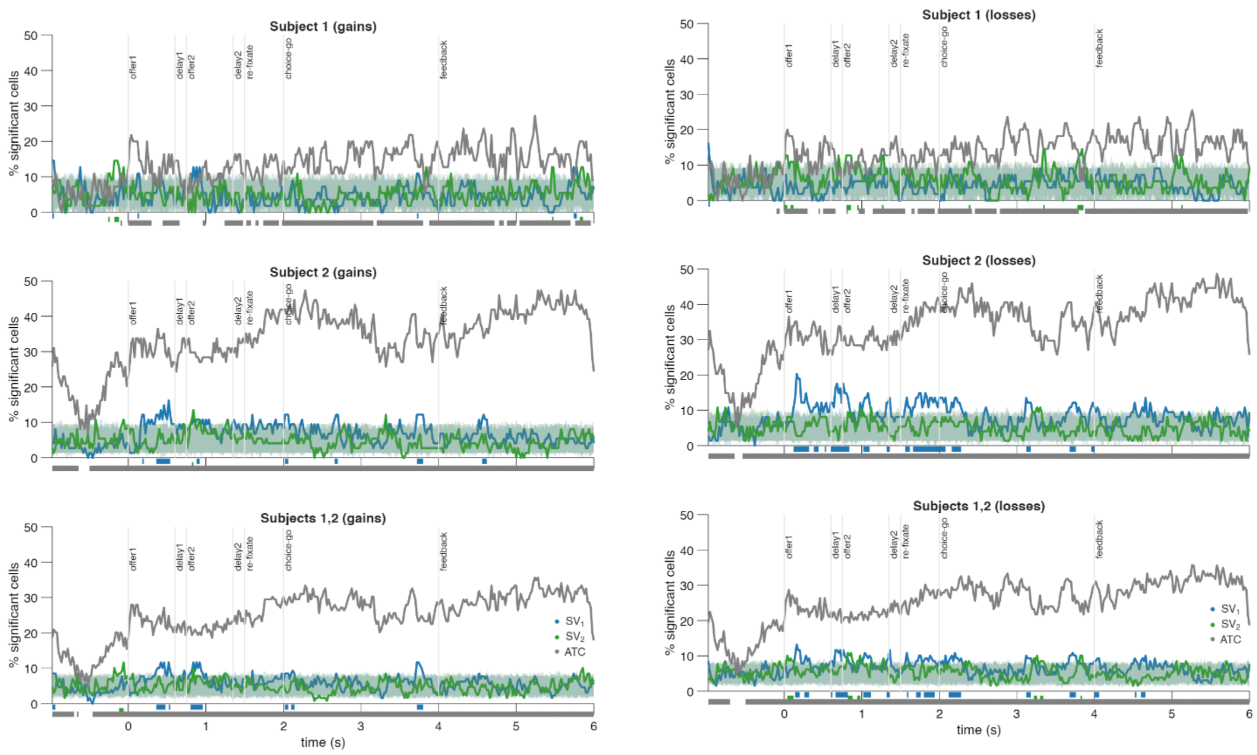


**Supplementary Figure S13.** Same as Fig. 5A, showing results for the 'linear spike-rate model without ATC interactions' for trials resulting in gains (left) and losses (right) defined by the 'RDV' utility function (Methods 2.4.3). Data from subject 1 (top), subject 2 (middle), or combined (bottom). The shaded areas show the 5th to 95th percentile of equivalent results from n=1000 trial order shuffled data.

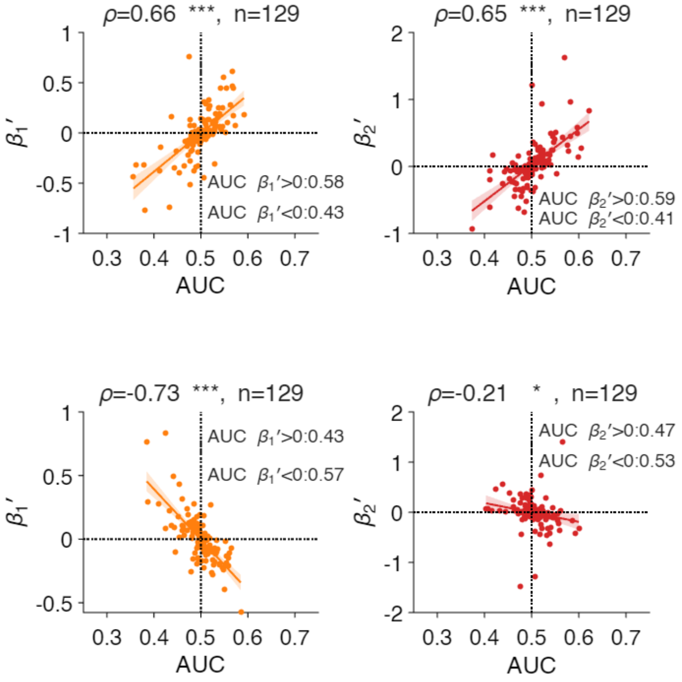


**Supplementary Figure S14.** Same as Fig. 5D, showing results for the 'linear spike-rate model without ATC interactions' for trials resulting in gains (top) and losses (bottom) defined by the 'RDV' utility function (Methods 2.4.3). Data combined in the two subjects.

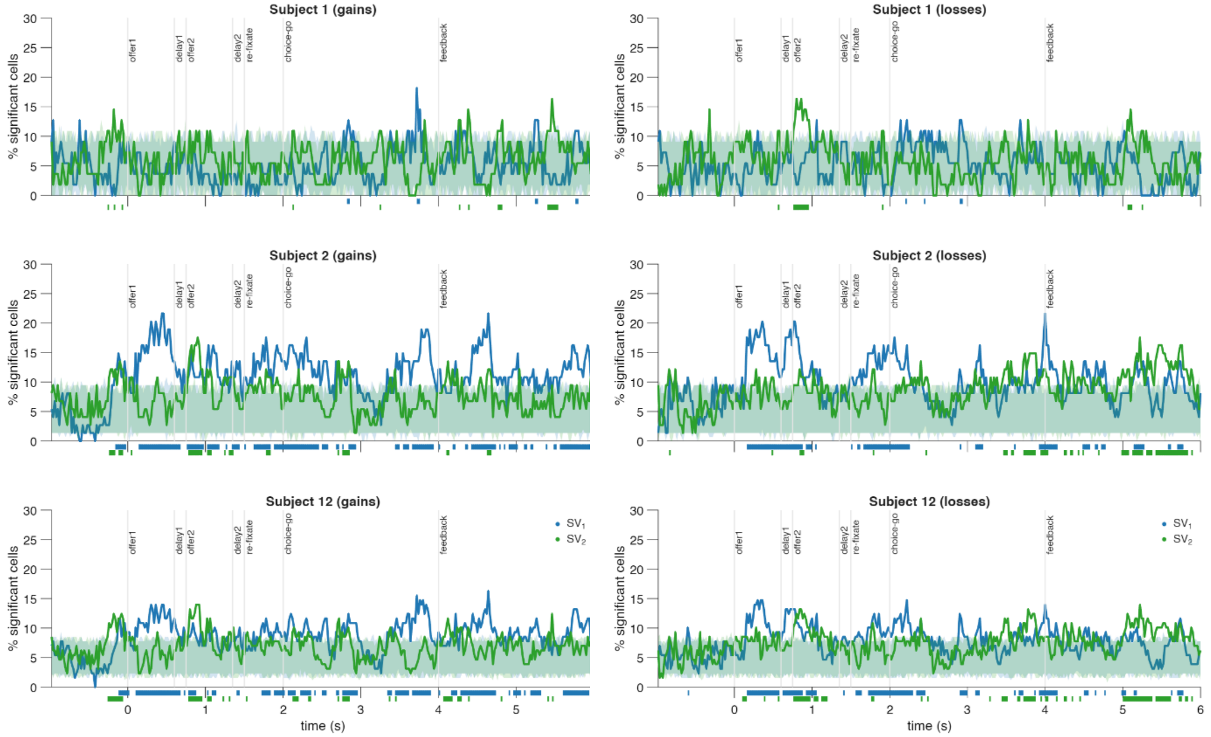


**Supplementary Figure S15.** Same as Fig. 6A, showing results for the 'linear spike-rate model with ATC interactions' for trials resulting in gains (left) losses (left) defined by the 'RDV' utility function (Methods 2.4.3). Data from subject 1 (top), subject 2 (middle), or combined (bottom). The shaded areas show the 5th to 95th percentile of equivalent results from n=1000 trial order shuffled data.

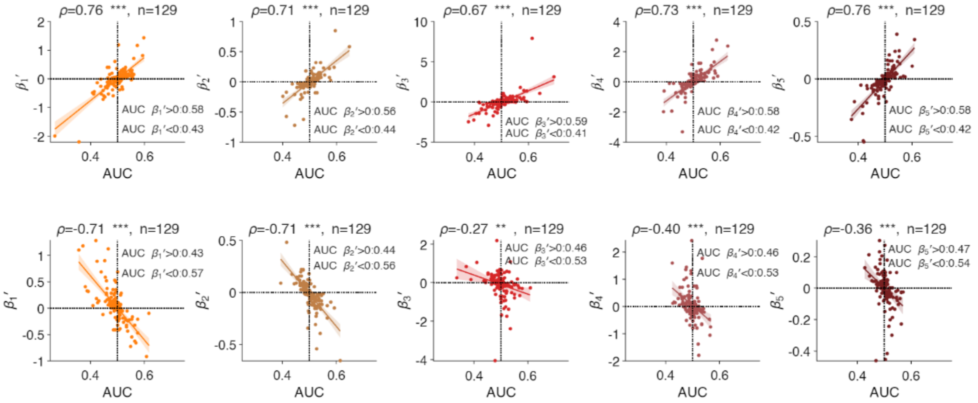


**Supplementary Figure S16.** Same as Fig 6D, showing results for the 'linear spike-rate model with ATC interactions' for trials resulting in gains (top) or losses (bottom) defined by the 'RDV' utility function (Methods 2.4.3). Data combined in the two subjects.

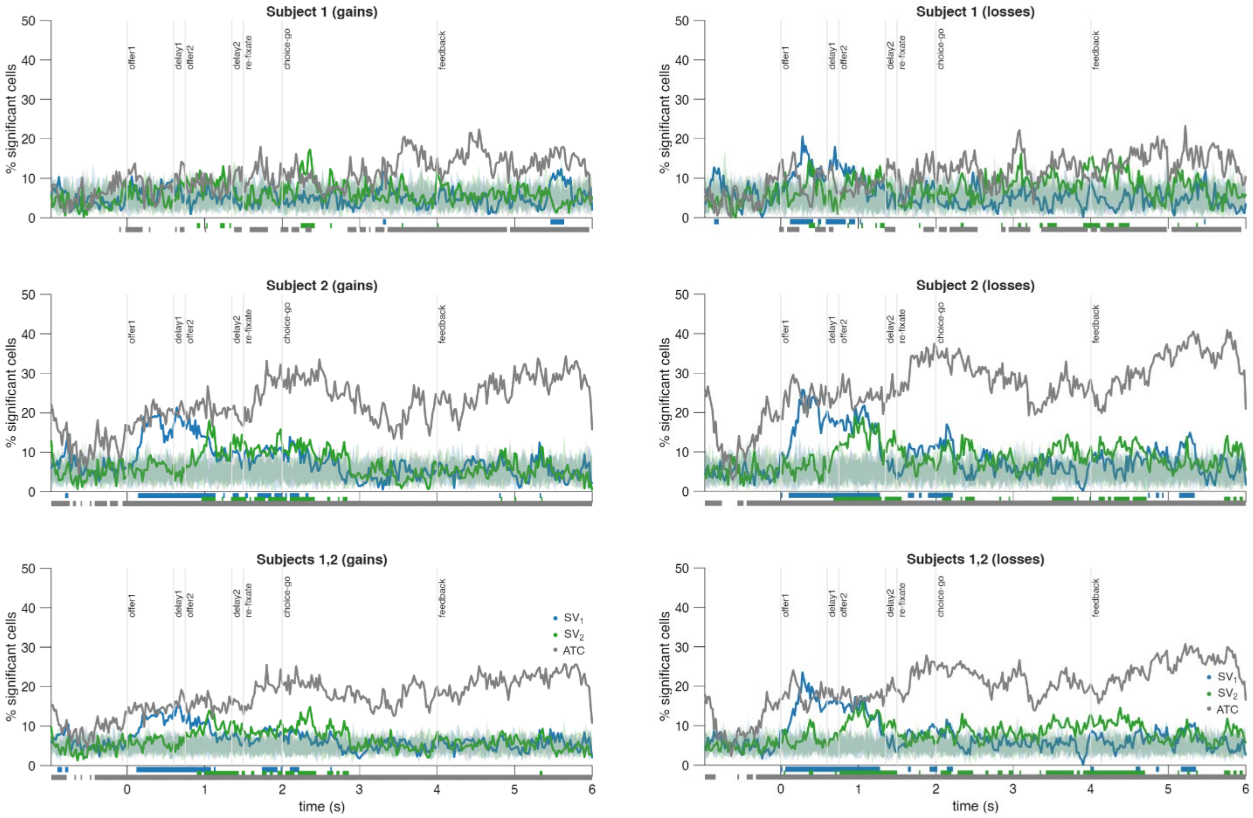


**Supplementary Figure S17.** Same as Figure 5A, showing results for the 'linear spike-rate model without ATC interactions', by considering n=10 subsets of trials randomly drawn from the whole set of trials whose size equals the number of trials resulting in gains (left) or losses (right) defined by the 'RDV' utility function (Methods 2.4.3). The shaded areas show the $5^{th}$ to $95^{th}$ percentile of equivalent results from n=100 trial order shuffled data.

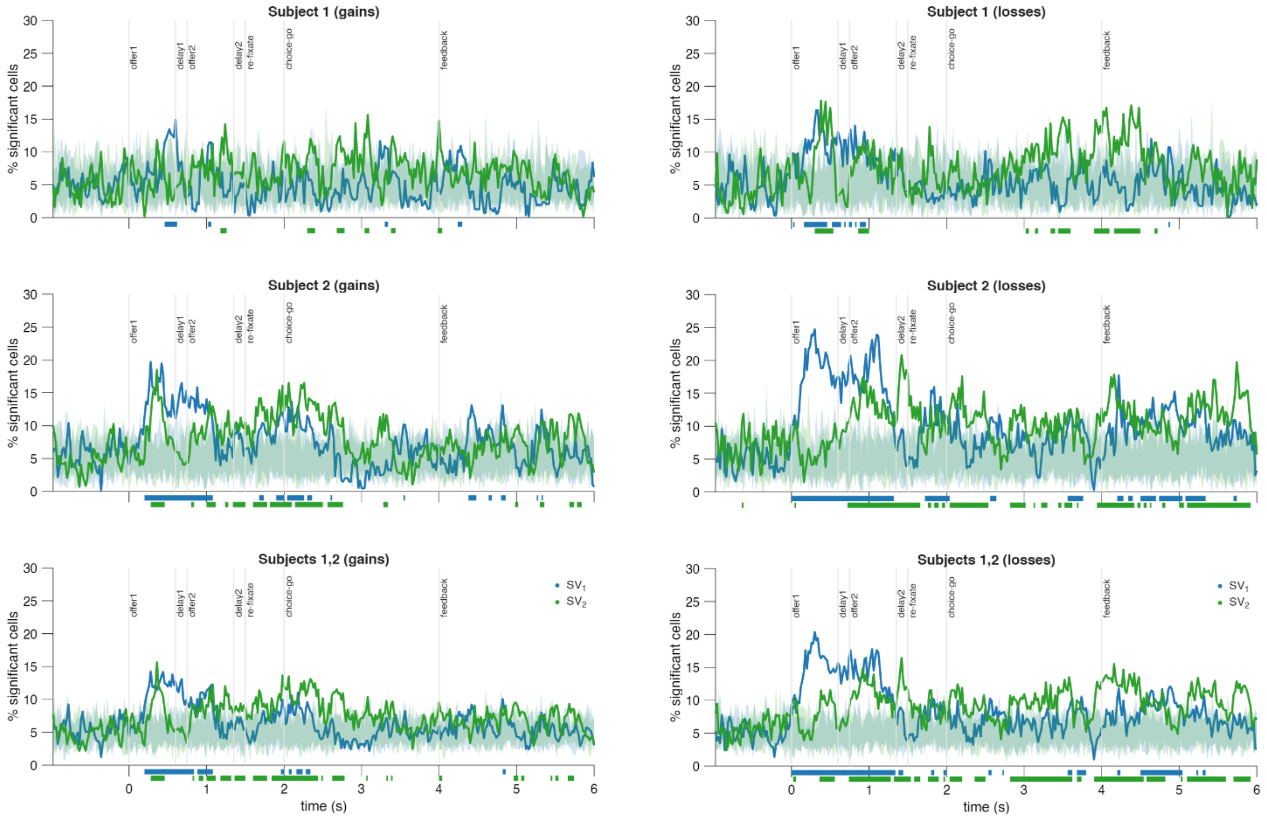


**Supplementary Figure S18.** Same as Figure 6A, showing results for the ‘linear spike-rate model with ATC interactions’, by considering n=10 subsets of trials randomly drawn from the whole set of trials whose size equals the number of trials resulting in gains (left) or losses (right) defined by the ‘RDV’ utility function (Methods 2.4.3). The shaded areas show the 5$^{th}$ to 95$^{th}$ percentile of equivalent results from n=100 trial order shuffled data.

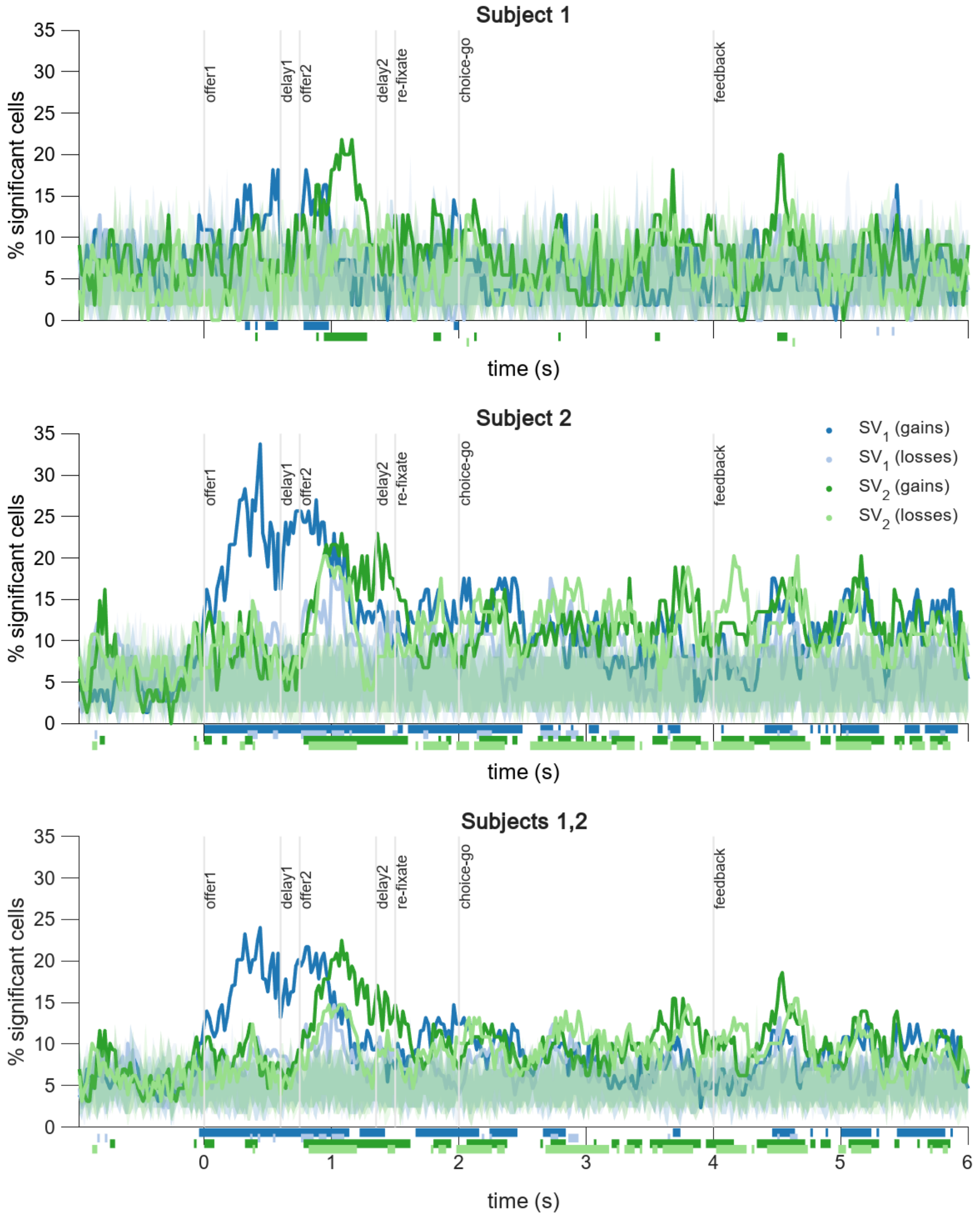


**Supplementary Figure S19.** Same as Figure 7A, but defining $SV_1^- = -\hat{\beta}_2 EU_1^- + \hat{\beta}_6 VU_1^-$, and $SV_2^- = -\hat{\beta}_4 EU_2^- + \hat{\beta}_8 VU_2^-$.

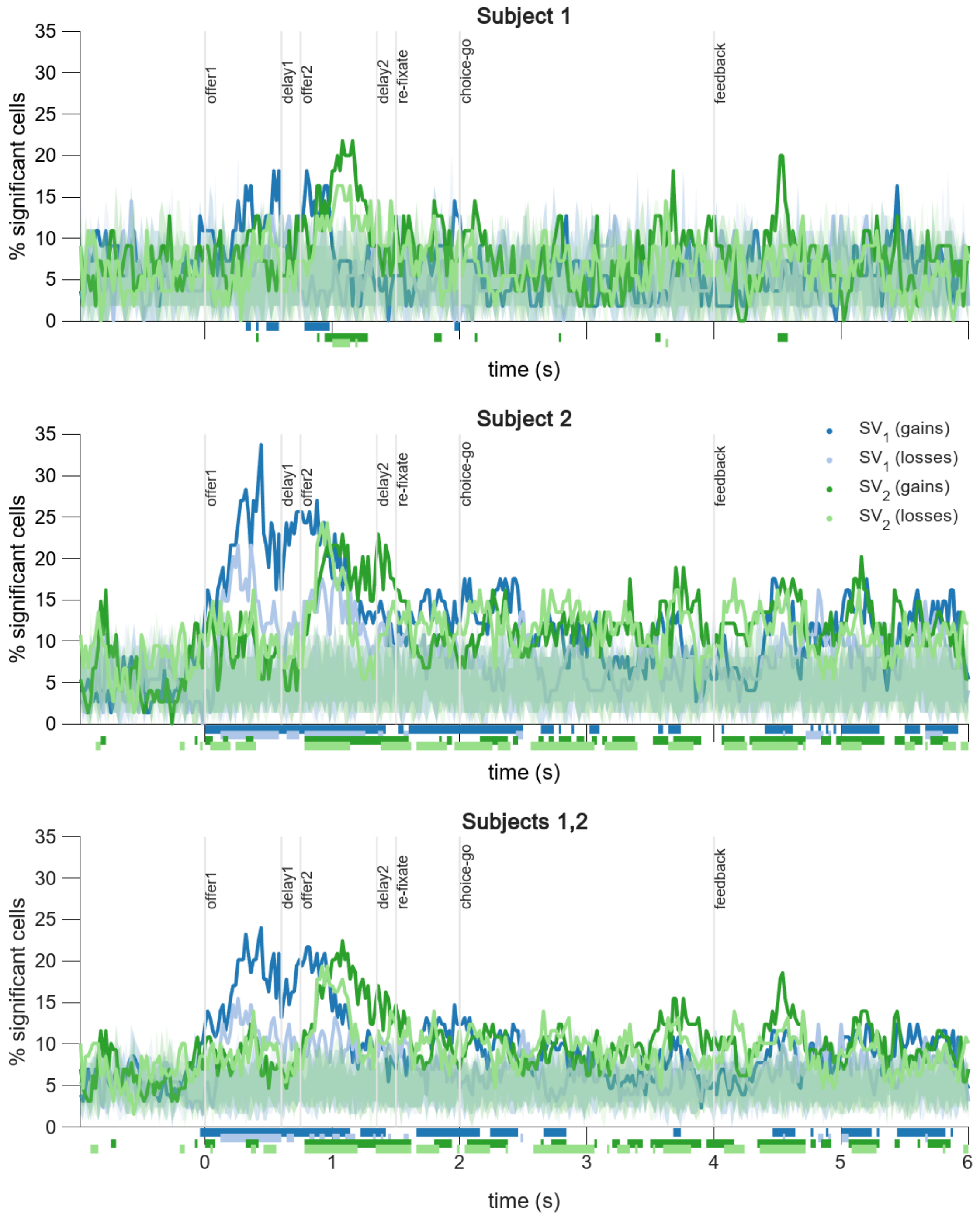


**Supplementary Figure S20.** Same as Figure 7A, but for z-scored $SV_1^-$, $SV_1^+$, $SV_2^+$ and $SV_2^-$.

| Subject | Model | $n_\Theta$ | Accuracy (%) | $R^2_{adj}$ | -log(L) | AIC | BIC |
|---|---|---|---|---|---|---|---|
| 1 | linear | 6 | 92.12 ± 0.32 | 0.687±0.0090 | 178.3±6.09 | 368.7±12.18 | 397.2±12.18 |
| **1** | **linear_RDV** | **10** | **92.26 ± 0.32** | **0.698±0.0088** | **148.9±4.86** | **317.8±9.72** | **365.2±9.72** |
| 1 | TDM | 4 | 92.04 ± 0.33 | 0.660±0.0082 | 192.4±5.97 | 392.8±11.94 | 411.7±11.94 |
| 1 | TDM_RDV | 11 | 91.98 ± 0.31 | 0.654±0.0081 | 196.3±5.84 | 414.6±11.68 | 466.8±11.68 |
| 1 | TDM_RDV and_linear_RDV | 13 | 92.28±0.32 | 0.696±0.0089 | 173.5±5.99 | 373.1±11.98 | 434.8±11.98 |
| 2 | linear | 6 | 88.84 ± 0.33 | 0.585±0.0078 | 224.5±6.07 | 461.0±12.14 | 489.5±12.14 |
| **2** | **linear_RDV** | **10** | **89.27 ± 0.33** | **0.610±0.0075** | **195.2±5.27** | **410.4±10.53** | **457.8±10.53** |
| 2 | TDM | 4 | 88.81 ± 0.33 | 0.572±0.0071 | 232.0±5.89 | 472.1±11.77 | 491.0±11.77 |
| 2 | TDM_RDV | 11 | 88.75 ± 0.33 | 0.570±0.0070 | 231.7±5.75 | 485.4±11.51 | 537.6±11.51 |
| 2 | TDM_RDV and_linear_RDV | 13 | 89.27±0.33 | 0.606±0.074 | 212.2±5.69 | 450.4±11.39 | 512.1±11.39 |
| 1, 2 | linear | 6 | 90.41 ± 0.25 | 0.634±0.0068 | 202.3±4.56 | 416.7±9.12 | 445.1±9.12 |
| **1, 2** | **linear_RDV** | **10** | **90.71 ± 0.25** | **0.653±0.0064** | **173.0±3.91** | **365.9±7.81** | **413.4±7.81** |
| 1, 2 | TDM | 4 | 90.36 ± 0.26 | 0.614±0.0061 | 213.0±4.39 | 434.0±8.77 | 453.0±8.77 |
| 1, 2 | TDM_RDV | 11 | 90.31 ± 0.25 | 0.611±0.0060 | 214.7±4.26 | 451.4±8.51 | 503.6±8.51 |
| 1,2 | TDM_RDV and_linear_RDV | 13 | 90.71±0.25 | 0.649±0.0065 | 193.6±4.31 | 413.3±8.63 | 475.0±8.63 |

**Supplementary Table ST14.** Performances for model comparisons (mean ± s.e.m across sessions). The column $n_\Theta$ shows the number of parameters fit in each model. Subjects are combined be pooling session by session results ($n = 109$ in subject 1, 118 in subject 2.

| Subject | Model | Parameters |
|---|---|---|
| 1 | Linear | $\beta_0$: -0.24 ± 0.02 $^{***}$, $\beta_1$: 2.08 ± 0.07 $^{***}$, $\beta_2$: 0.51 ± 0.02 $^{***}$,<br>$\beta_3$: 0.12 ± 0.02 $^{***}$, $\beta_4$: 0.07 ± 0.01 $^{***}$, $\beta_5$: -0.01 ± 0.00 $^{***}$<br>$p=5.61\cdot10^{-18}$ ; $p=1.72\cdot10^{-19}$ ; $p=1.72\cdot10^{-19}$ ;<br>$p=2.52\cdot10^{-10}$ ; $p=2.54\cdot10^{-6}$; $p=1.11\cdot10^{-4}$ |
| **1** | **Linear_RDV** | **$\kappa_0$: 2.61 ± 0.25 $^{***}$, $\kappa_1$: 3.28 ± 0.05 $^{***}$, $\gamma_0$: 0.79 ± 0.02 $^{***}$, $\gamma_1$: 0.02 ± 0.00 $^{***}$,**<br>**$\lambda_0$: 0.49 ± 0.03 $^{***}$, $\lambda_1$: 0.18 ± 0.02 $^{***}$, $\lambda_2$: -0.02 ± 0.00 $^{***}$,**<br>**$\beta_0$: -0.25 ± 0.02 $^{***}$, $\beta_1$: 3.37 ± 0.08 $^{***}$, $\beta_2$: 0.72 ± 0.04 $^{***}$**<br>$p=1.72\cdot10^{-19}$ ; $p=1.72\cdot10^{-19}$ ; $p=1.72\cdot10^{-19}$ ; $p=1.72\cdot10^{-19}$ ;<br>$p=1.72\cdot10^{-19}$ ; $p=1.72\cdot10^{-19}$ ; $p=3.03\cdot10^{-5}$ ;<br>$p=5.37\cdot10^{-18}$ ; $p=1.72\cdot10^{-19}$ ; $p=1.72\cdot10^{-19}$ |
| 1 | TDM | $\tau$: 3.45 ± 0.04 $^{***}$, $\beta_0$: -0.15 ± 0.01 $^{***}$, $\beta_1$: 10.95 ± 0.22 $^{***}$, $\beta_2$: 2.76 ± 0.14 $^{***}$<br>$p=1.72\cdot10^{-19}$ ; $p=1.32\cdot10^{-17}$ ; $p=1.72\cdot10^{-19}$ ; $p=2.70\cdot10^{-19}$ |
| 1 | TDM_RDV | $\tau$: 3.60 ± 0.04 $^{***}$, $\kappa_0$: 6.42 ± 0.27 $^{***}$, $\kappa_1$: 3.94 ± 0.05 $^{***}$, $\gamma_0$: 1.00 ± 0.00 $^{***}$,<br>$\gamma_1$: 0.10 ± 0.01 $^{***}$, $\lambda_0$: 0.96 ± 0.01 $^{***}$, $\lambda_1$: 0.83 ± 0.02 $^{***}$, $\lambda_2$: -0.07 ± 0.01 $^{***}$,<br>$\beta_0$: -0.16 ± 0.01 $^{***}$, $\beta_1$: 10.40 ± 0.18 $^{***}$, $\beta_2$: 2.30 ± 0.17 $^{***}$<br>$p=1.72\cdot10^{-19}$ ; $p=1.72\cdot10^{-19}$ ; $p=1.72\cdot10^{-19}$ ; $p=8.50\cdot10^{-25}$ ;<br>$p=4.85\cdot10^{-18}$ ; $p=8.75\cdot10^{-19}$ ; $p=7.27\cdot10^{-20}$ ; $p=1.38\cdot10^{-16}$ ;<br>$p=4.17\cdot10^{-17}$ ; $p=1.72\cdot10^{-19}$ ; $p=8.27\cdot10^{-18}$ |
| 1 | TDM_RDV and_linear_RDV | $\tau$: 3.31 ± 0.04 $^{***}$, $\kappa_0$: 2.57 ± 0.28 $^{***}$, $\kappa_1$: 3.88 ± 0.09 $^{***}$, $\gamma_0$: 0.79 ± 0.02 $^{***}$, $\gamma_1$: 0.03 ± 0.00 $^{***}$,<br>$\lambda_0$: 0.48 ± 0.03 $^{***}$, $\lambda_1$: 0.14 ± 0.02 $^{***}$, $\lambda_2$: -0.02 ± 0.00 $^{***}$, $\beta_0$: -0.26 ± 0.02 $^{***}$,<br>$\beta_1$: 0.50 ± 0.06 $^{***}$, $\beta_2$: 0.17 ± 0.02 $^{***}$, $\beta_3$: 3.48 ± 0.08 $^{***}$, $\beta_4$: 0.64 ± 0.04 $^{***}$<br>$p=8.90\cdot10^{-20}$ ; $p=1.72\cdot10^{-19}$ ; $p=1.72\cdot10^{-19}$ ; $p=1.72\cdot10^{-19}$ ; $p=1.32\cdot10^{-15}$ ;<br>$p=3.53\cdot10^{-19}$ ; $p=9.52\cdot10^{-17}$ ; $p=3.01\cdot10^{-5}$ ; $p=3.91\cdot10^{-18}$ ;<br>$p=1.19\cdot10^{-11}$ ; $p=1.25\cdot10^{-14}$ ; $p=1.72\cdot10^{-19}$ ; $p=1.75\cdot10^{-19}$ |
| 2 | Linear | $\beta_0$: -0.44 ± 0.03 $^{***}$, $\beta_1$: 1.25 ± 0.04 $^{***}$, $\beta_2$: 0.48 ± 0.02 $^{***}$,<br>$\beta_3$: 0.19 ± 0.02 $^{***}$, $\beta_4$: 0.17 ± 0.01 $^{***}$, $\beta_5$: -0.04 ± 0.00 $^{***}$<br>$p=1.33\cdot10^{-20}$ ; $p=8.42\cdot10^{-21}$ ; $p=8.42\cdot10^{-21}$ ;<br>$p=1.67\cdot10^{-19}$ ; $p=1.86\cdot10^{-17}$ ; $p=1.57\cdot10^{-16}$ |
| **2** | **Linear_RDV** | **$\kappa_0$: 0.97 ± 0.12 $^{***}$, $\kappa_1$: 3.09 ± 0.06 $^{***}$, $\gamma_0$: 0.66 ± 0.02 $^{***}$, $\gamma_1$: 0.03 ± 0.00 $^{***}$,**<br>**$\lambda_0$: 0.27 ± 0.02 $^{***}$, $\lambda_1$: 0.16 ± 0.02 $^{***}$, $\lambda_2$: -0.01 ± 0.00 $^{*}$,**<br>**$\beta_0$: -0.47 ± 0.03 $^{***}$, $\beta_1$: 3.28 ± 0.06 $^{***}$, $\beta_2$: 0.64 ± 0.05 $^{***}$**<br>$p=8.42\cdot10^{-21}$ ; $p=8.42\cdot10^{-21}$ ; $p=8.42\cdot10^{-21}$ ; $p=8.42\cdot10^{-21}$ ;<br>$p=8.42\cdot10^{-21}$ ; $p=8.42\cdot10^{-21}$ ; $p=3.47\cdot10^{-2}$ ;<br>$p=1.39\cdot10^{-20}$ ; $p=8.42\cdot10^{-21}$ ; $p=1.95\cdot10^{-16}$ |
| 2 | TDM | $\tau$: 3.34 ± 0.04 $^{***}$, $\beta_0$: -0.33 ± 0.02 $^{***}$, $\beta_1$: 9.35 ± 0.17 $^{***}$, $\beta_2$: 3.50 ± 0.17 $^{***}$<br>$p=8.42\cdot10^{-21}$ ; $p=1.74\cdot10^{-20}$ ; $p=8.42\cdot10^{-21}$ ; $p=1.22\cdot10^{-20}$ |
| 2 | TDM_RDV | $\tau$: 3.31 ± 0.04 $^{***}$, $\kappa_0$: 7.83 ± 0.22 $^{***}$, $\kappa_1$: 3.69 ± 0.05 $^{***}$, $\gamma_0$: 0.99 ± 0.01 $^{***}$,<br>$\gamma_1$: 0.05 ± 0.00 $^{***}$, $\lambda_0$: 0.78 ± 0.03 $^{***}$, $\lambda_1$: 0.71 ± 0.02 $^{***}$, $\lambda_2$: -0.05 ± 0.01 $^{***}$,<br>$\beta_0$: -0.34 ± 0.02 $^{***}$, $\beta_1$: 9.50 ± 0.15 $^{***}$, $\beta_2$: 3.39 ± 0.18 $^{***}$<br>$p=8.42\cdot10^{-21}$ ; $p=7.13\cdot10^{-21}$ ; $p=8.42\cdot10^{-21}$ ; $p=2.36\cdot10^{-26}$ ;<br>$p=3.53\cdot10^{-19}$ ; $p=3.22\cdot10^{-21}$ ; $p=8.42\cdot10^{-21}$ ; $p=1.91\cdot10^{-15}$ ;<br>$p=1.81\cdot10^{-20}$ ; $p=8.42\cdot10^{-21}$ ; $p=1.34\cdot10^{-20}$ |
| 2 | TDM_RDV and_linear_RDV | $\tau$: 3.23 ± 0.03 $^{***}$, $\kappa_0$: 1.03 ± 0.14 $^{***}$, $\kappa_1$: 3.21 ± 0.08 $^{***}$, $\gamma_0$: 0.63 ± 0.02 $^{***}$, $\gamma_1$: 0.04 ± 0.00 $^{***}$,<br>$\lambda_0$: 0.35 ± 0.02 $^{***}$, $\lambda_1$: 0.15 ± 0.02 $^{***}$, $\lambda_2$: -0.01 ± 0.00 $^{***}$, $\beta_0$: -0.48 ± 0.03 $^{***}$,<br>$\beta_1$: 0.57 ± 0.05 $^{***}$, $\beta_2$: 0.12 ± 0.01 $^{***}$, $\beta_3$: 3.26 ± 0.07 $^{***}$, $\beta_4$: 0.60 ± 0.06 $^{***}$<br>$p=4.25\cdot10^{-21}$ ; $p=8.42\cdot10^{-21}$ ; $p=8.42\cdot10^{-21}$ ; $p=8.42\cdot10^{-21}$ ; $p=4.85\cdot10^{-18}$ ;<br>$p=1.04\cdot10^{-19}$ ; $p=3.41\cdot10^{-18}$ ; $p=8.54\cdot10^{-4}$ ; $p=1.41\cdot10^{-20}$ ;<br>$p=2.58\cdot10^{-14}$ ; $p=8.63\cdot10^{-15}$ ; $p=8.42\cdot10^{-21}$ ; $p=4.78\cdot10^{-15}$ |

| Subject | Model | Parameters |
|---|---|---|
| 1,2 | Linear | $\beta_0$: -0.35 ± 0.02 ***, $\beta_1$: 1.65 ± 0.05 ***, $\beta_2$: 0.49 ± 0.01 ***,<br>$\beta_3$: 0.15 ± 0.01 ***, $\beta_4$: 0.12 ± 0.01 ***, $\beta_5$: -0.02 ± 0.00 ***<br>$p=7.52\cdot10^{-37}$; $p=3.10\cdot10^{-38}$; $p=3.10\cdot10^{-38}$;<br>$p=1.47\cdot10^{-28}$; $p=8.99\cdot10^{-22}$; $p=1.72\cdot10^{-19}$ |
| **1,2** | **Linear_RDV** | **$\kappa_0$: 1.76 ± 0.14 ***, $\kappa_1$: 3.18 ± 0.04 ***, $\gamma_0$: 0.72 ± 0.01 ***, $\gamma_1$: 0.03 ± 0.00 ***,**<br>**$\lambda_0$: 0.38 ± 0.02 ***, $\lambda_1$: 0.17 ± 0.01 ***, $\lambda_2$: -0.01 ± 0.00 ***,**<br>**$\beta_0$: -0.37 ± 0.02 ***, $\beta_1$: 3.33 ± 0.05 ***, $\beta_2$: 0.68 ± 0.03 *****<br>$p=3.10\cdot10^{-38}$; $p=3.10\cdot10^{-38}$; $p=3.10\cdot10^{-38}$; $p=3.10\cdot10^{-38}$;<br>$p=3.10\cdot10^{-38}$; $p=3.10\cdot10^{-38}$; $p=7.08\cdot10^{-6}$;<br>$p=8.05\cdot10^{-37}$; $p=3.10\cdot10^{-38}$; $p=3.85\cdot10^{-34}$ |
| 1,2 | TDM | $\tau$: 3.39 ± 0.03 ***, $\beta_0$: -0.25 ± 0.01 ***, $\beta_1$: 10.12 ± 0.15 ***, $\beta_2$: 3.15 ± 0.11 ***<br>$p=3.10\cdot10^{-38}$; $p=1.62\cdot10^{-36}$; $p=3.10\cdot10^{-38}$; $p=6.32\cdot10^{-38}$; |
| 1,2 | TDM_RDV | $\tau$: 3.45 ± 0.03 ***, $\kappa_0$: 7.15 ± 0.18 ***, $\kappa_1$: 3.81 ± 0.04 ***, $\gamma_0$: 0.99 ± 0.00 ***,<br>$\gamma_1$: 0.07 ± 0.00 ***, $\lambda_0$: 0.87 ± 0.02 ***, $\lambda_1$: 0.77 ± 0.02 ***, $\lambda_2$: -0.06 ± 0.00 ***,<br>$\beta_0$: -0.26 ± 0.01 ***, $\beta_1$: 9.93 ± 0.12 ***, $\beta_2$: 2.87 ± 0.13 ***<br>$p=3.10\cdot10^{-38}$; $p=3.10\cdot10^{-38}$; $p=3.10\cdot10^{-38}$; $p=2.43\cdot10^{-48}$;<br>$p=4.22\cdot10^{-35}$; $p=1.28\cdot10^{-40}$; $p=3.10\cdot10^{-38}$; $p=6.45\cdot10^{-30}$;<br>$p=4.11\cdot10^{-36}$; $p=3.10\cdot10^{-38}$; $p=8.74\cdot10^{-37}$ |
| 1,2 | TDM_RDV and_linear_RDV | $\tau$: 3.27 ± 0.02 ***, $\kappa_0$: 1.77 ± 0.16 ***, $\kappa_1$: 3.53 ± 0.06 ***, $\gamma_0$: 0.71 ± 0.01 ***, $\gamma_1$: 0.03 ± 0.00 ***,<br>$\lambda_0$: 0.41 ± 0.02 ***, $\lambda_1$: 0.15 ± 0.01 ***, $\lambda_2$: -0.01 ± 0.00 ***, $\beta_0$: -0.37 ± 0.02 ***,<br>$\beta_1$: 0.54 ± 0.04 ***, $\beta_2$: 0.14 ± 0.01 ***, $\beta_3$: 3.36 ± 0.05 ***, $\beta_4$: 0.62 ± 0.04 ***<br>$p=3.10\cdot10^{-38}$; $p=3.10\cdot10^{-38}$; $p=3.10\cdot10^{-38}$; $p=3.10\cdot10^{-38}$; $p=1.52\cdot10^{-31}$;<br>$p=7.88\cdot10^{-37}$; $p=7.83\cdot10^{-33}$; $p=9.58\cdot10^{-8}$; $p=7.52\cdot10^{-37}$;<br>$p=7.97\cdot10^{-24}$; $p=2.59\cdot10^{-27}$; $p=3.10\cdot10^{-38}$; $p=7.65\cdot10^{-32}$ |

**Supplementary Table ST15.** Parameter estimates (mean ± s.e.m) across sessions and $k = 4$ cross-validation folds for the models tested. Subjects are combined be pooling session by session results ($n = 109$ in subject 1, $118$ in subject 2). All parameters show significant magnitude, assessed via two-sided signed-rank tests, FDR corrected via Benjamini-Hochberg procedure.

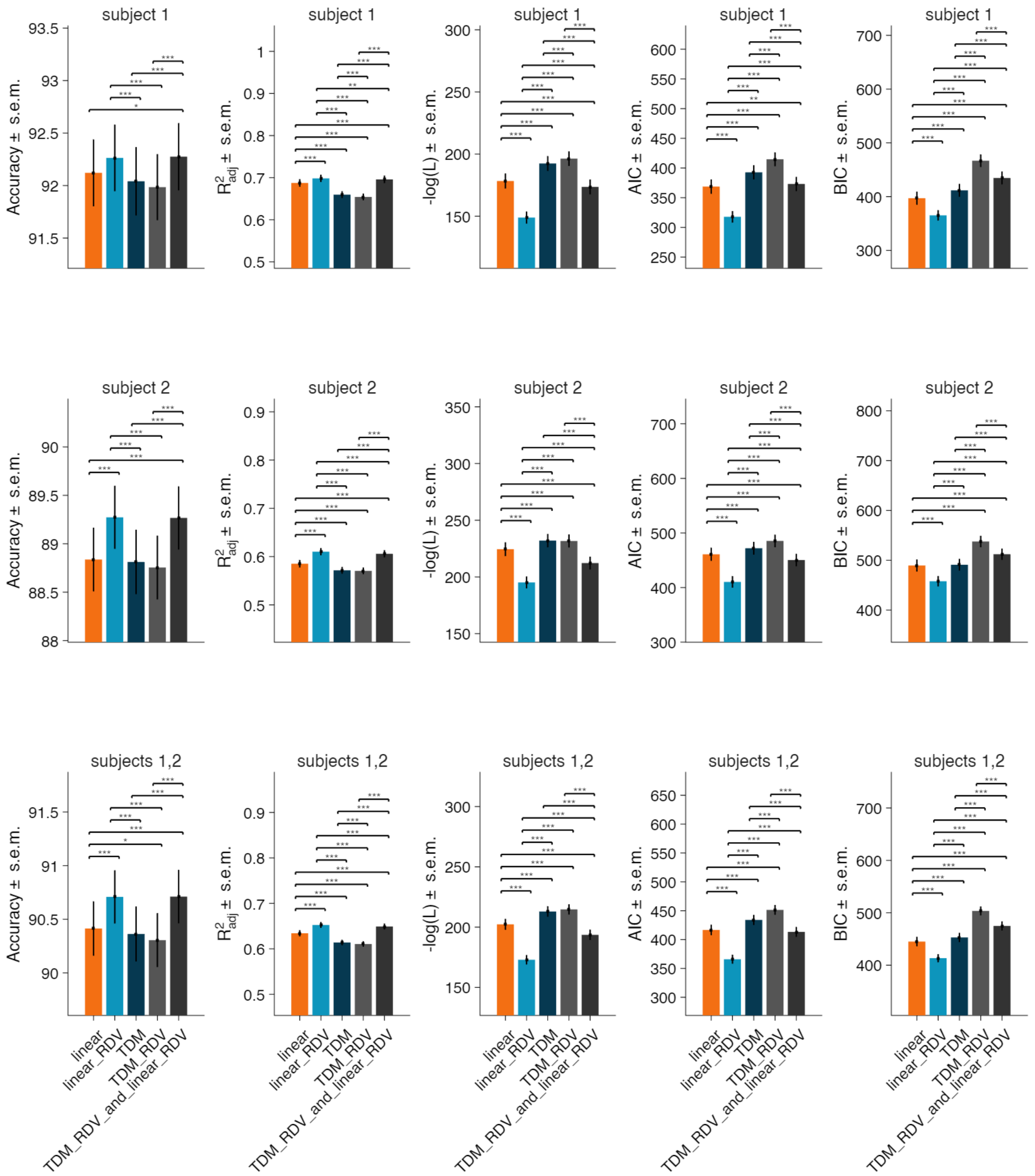


**Supplementary Figure S21.** Model performances (mean ± s.e.m) across sessions. P-values are computed across sessions via signed-rank tests (*** p<0.001, **p<0.01, *p<0.05, listed in Supp. Table S16), pooled and FDR corrected via Benjamini-Hochberg procedure. Subjects are combined be pooling session by session results ($n = 109$ in subject 1, 118 in subject 2).

| Subject | Model comparison | Accuracy | R²_adj | −log(L) | AIC | BIC |
|---|---|---|---|---|---|---|
| 1 | linear vs linear_RDV | p=5.35e-02 | p=5.41e-10 | p=2.30e-16 | p=3.11e-14 | p=2.84e-08 |
| 1 | linear vs TDM | p=4.83e-01 | p=4.98e-18 | p=1.50e-17 | p=3.67e-16 | p=2.21e-10 |
| 1 | linear vs TDM_RDV | p=7.21e-02 | p=1.90e-18 | p=3.12e-18 | p=4.66e-19 | p=3.52e-19 |
| 1 | linear vs TDM_RDV_and_linear_RDV | p=3.08e-02 | p=1.90e-06 | p=1.70e-07 | p=5.62e-03 | p=1.22e-18 |
| 1 | linear_RDV vs TDM | p=7.28e-04 | p=1.31e-18 | p=6.55e-19 | p=1.83e-18 | p=3.18e-12 |
| 1 | linear_RDV vs TDM_RDV | p=1.36e-04 | p=4.32e-19 | p=3.52e-19 | p=3.52e-19 | p=3.52e-19 |
| 1 | linear_RDV vs TDM_RDV_and_linear_RDV | p=8.57e-01 | p=1.67e-03 | p=3.07e-14 | p=9.43e-18 | p=3.52e-19 |
| 1 | TDM vs TDM_RDV | p=1.67e-01 | p=1.94e-05 | p=2.44e-04 | p=7.73e-18 | p=3.52e-19 |
| 1 | TDM vs TDM_RDV_and_linear_RDV | p=7.92e-04 | p=1.11e-18 | p=3.33e-17 | p=1.76e-12 | p=1.34e-14 |
| 1 | TDM_RDV vs TDM_RDV_and_linear_RDV | p=5.40e-05 | p=4.66e-19 | p=1.90e-18 | p=3.82e-18 | p=2.21e-16 |
| 2 | linear vs linear_RDV | p=2.30e-06 | p=1.83e-17 | p=7.19e-20 | p=9.63e-18 | p=1.46e-08 |
| 2 | linear vs TDM | p=7.89e-01 | p=1.12e-11 | p=2.75e-13 | p=3.84e-09 | p=3.27e-01 |
| 2 | linear vs TDM_RDV | p=1.78e-01 | p=1.35e-10 | p=5.51e-10 | p=1.26e-17 | p=1.75e-20 |
| 2 | linear vs TDM_RDV_and_linear_RDV | p=6.44e-06 | p=1.49e-14 | p=3.85e-18 | p=1.78e-06 | p=6.85e-16 |
| 2 | linear_RDV vs TDM | p=3.15e-06 | p=3.59e-20 | p=2.15e-20 | p=3.52e-19 | p=4.41e-10 |
| 2 | linear_RDV vs TDM_RDV | p=1.53e-08 | p=2.04e-20 | p=1.75e-20 | p=1.75e-20 | p=1.75e-20 |
| 2 | linear_RDV vs TDM_RDV_and_linear_RDV | p=8.60e-01 | p=1.65e-08 | p=2.53e-12 | p=1.90e-18 | p=1.75e-20 |
| 2 | TDM vs TDM_RDV | p=4.65e-01 | p=2.13e-01 | p=1.50e-01 | p=5.16e-14 | p=1.75e-20 |
| 2 | TDM vs TDM_RDV_and_linear_RDV | p=7.98e-06 | p=6.48e-20 | p=3.79e-20 | p=6.14e-14 | p=4.97e-13 |
| 2 | TDM_RDV vs TDM_RDV_and_linear_RDV | p=3.79e-08 | p=2.92e-20 | p=2.70e-20 | p=3.80e-20 | p=1.85e-17 |
| 1:2 | linear vs linear_RDV | p=5.86e-07 | p=3.17e-26 | p=1.76e-34 | p=1.43e-29 | p=4.99e-16 |
| 1:2 | linear vs TDM | p=4.74e-01 | p=5.02e-29 | p=1.06e-29 | p=2.76e-24 | p=2.12e-08 |
| 1:2 | linear vs TDM_RDV | p=2.66e-02 | p=7.55e-29 | p=6.78e-28 | p=1.53e-35 | p=1.52e-37 |
| 1:2 | linear vs TDM_RDV_and_linear_RDV | p=1.06e-06 | p=7.39e-20 | p=2.35e-24 | p=9.73e-02 | p=1.37e-33 |
| 1:2 | linear_RDV vs TDM | p=1.27e-08 | p=9.35e-37 | p=2.40e-37 | p=7.31e-36 | p=9.95e-21 |
| 1:2 | linear_RDV vs TDM_RDV | p=7.72e-12 | p=1.89e-37 | p=1.52e-37 | p=1.52e-37 | p=1.52e-37 |
| 1:2 | linear_RDV vs TDM_RDV_and_linear_RDV | p=7.95e-01 | p=3.24e-10 | p=8.02e-25 | p=1.68e-34 | p=1.52e-37 |
| 1:2 | TDM vs TDM_RDV | p=1.25e-01 | p=5.58e-05 | p=1.55e-01 | p=1.12e-30 | p=1.52e-37 |
| 1:2 | TDM vs TDM_RDV_and_linear_RDV | p=3.86e-08 | p=1.43e-36 | p=1.70e-35 | p=9.80e-25 | p=1.05e-25 |
| 1:2 | TDM_RDV vs TDM_RDV_and_linear_RDV | p=1.07e-11 | p=2.40e-37 | p=9.35e-37 | p=3.31e-36 | p=6.15e-32 |

**Supplementary Table ST16.** P-values for model comparisons in Supp. Fig. S21.

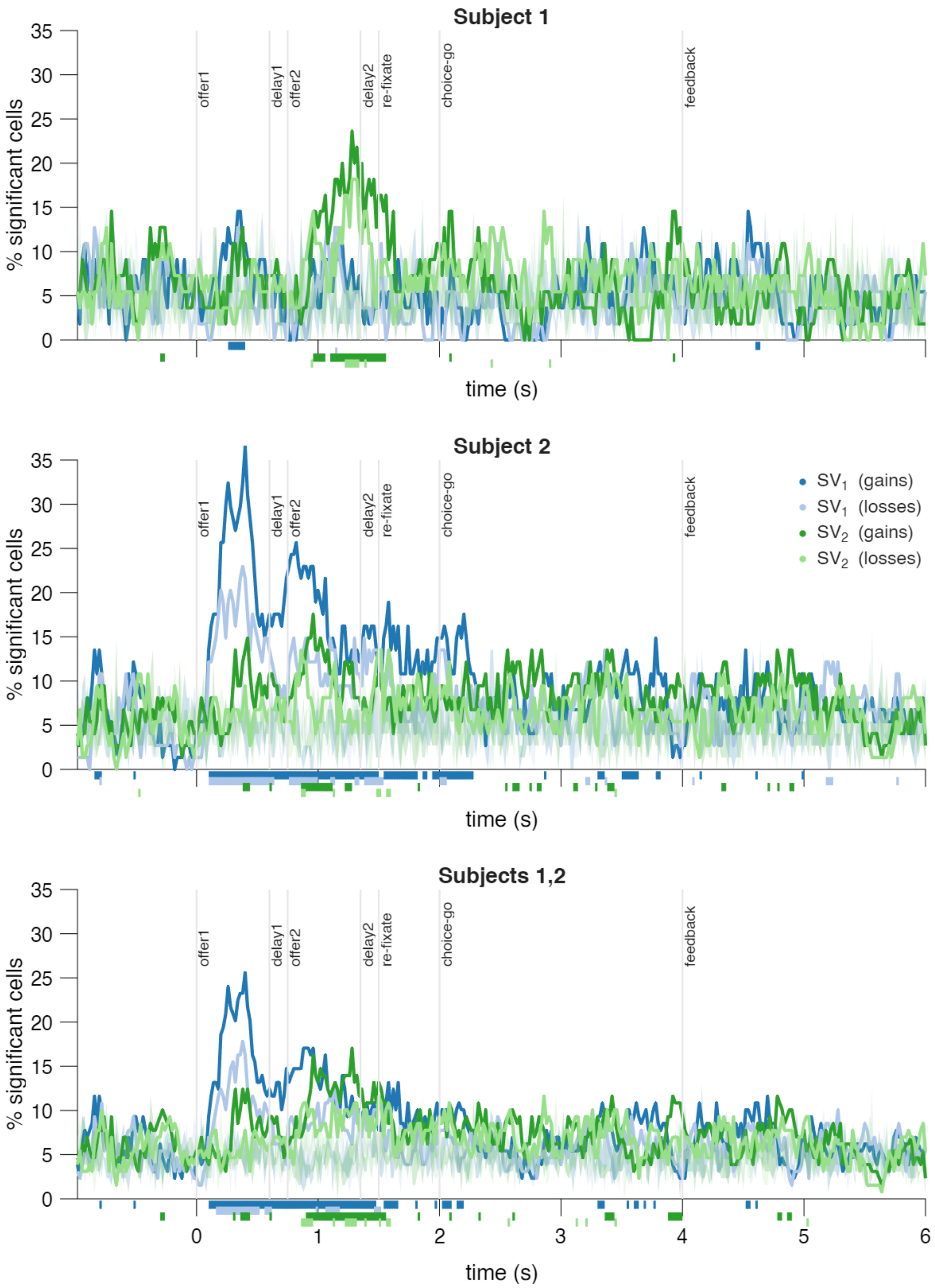


**Supplementary Figure S22.** Same as Figure 7A but using SV definition from the combined model 'TDM_RDV_and_linear_RDV' with both time-discounting and reference-dependent value regressors (Supplementary Methods *v.*).

**Supplementary Methods: comparing reference-dependency to time-discounting**

***i.* Linear Model with ATC interaction (linear)**

To compare the models, we include results for '**linear**' model of choice with ATC, which corresponds to Methods 2.4.2. We compute $EV$s and risk $R$ as:

$$EV = p^t v^t + p^b v^b, \qquad R = p^t (v^t - EV)^2 + p^b (v^b - EV)^2, \tag{1}$$

and regressed the choice for first option as:

$$\begin{aligned} \text{logit}\big(p(ch1)\big) = \beta_0 + \beta_1(EV_1 - EV_2) + \beta_2(EV_1 - EV_2)ATC + \beta_3(R_1 - R_2) \\ + \beta_4(R_1 - R_2)ATC + \beta_5(R_1 - R_2)ATC^2, \end{aligned} \tag{2}$$

i.e., we compute $\widehat{\Theta} = [\hat{\beta}_0, \hat{\beta}_1, \hat{\beta}_2, \hat{\beta}_3, \hat{\beta}_4, \hat{\beta}_5]$ and predict $\hat{p}_i$ that maximizes the likelihood of test data $y_i$, applying $k = 4$ fold cross-validation and ridge penalty to $\beta_1, \beta_2, \beta_3, \beta_4, \beta_5$. This is an upgraded of previous linear model where we include a linear interaction term for $EV$, and both linear and quadratic interaction terms for $R$. This extension follows from previous results showing approximately linear increase in $EV$ with $ATC$ (Fig. 4C, Supp. Fig. 1C) and inverted-u trend between $R$ and $ATC$ (Fig. 4D, Supp. Fig. 1D).

***ii.* Linear and Reference-Dependent Value (linear_RDV)**

Further, we test the reference-dependent linear model '**linear_RDV**' in Methods 2.4.3, where offered values are shaped by the RDV utility function $u(v, ATC)$, modeling the choice as:

$$\begin{aligned} \text{logit}\big(p(ch1)\big) = \beta_0 + \beta_1\big(\mathbb{E}[u(v_{1,}, ATC)] - \mathbb{E}[u(v_{2,}, ATC)]\big) \\ + \beta_2\big(VAR[u(v_{1,}, ATC)] - VAR[u(v_{2,}, ATC)]\big). \end{aligned} \tag{3}$$

This time we compute $\widehat{\Theta} = [\hat{\kappa}_0, \hat{\kappa}_1, \hat{\gamma}_0, \hat{\gamma}_1, \hat{\lambda}_0, \hat{\lambda}_1, \hat{\lambda}_2, \hat{\beta}_0, \hat{\beta}_1, \hat{\beta}_2]$ and predict $\hat{p}_i$ to maximize the likelihood of test data $y_i$, and applying $k = 4$ fold cross-validation and ridge penalty to $\beta_1, \beta_2$.

***iii.* Temporal discounting model (TDM)**

We implemented a temporal discounting model (TDM), for which the subjective value (SV) of an offer is computed as the discounted change in jackpot value based on expected time to achieve jackpot reward $T$, incorporating expected value and risk factors. The SV was used to fit a logistic regression model of choice.

We used the following definitions:

$$T \propto 6 - ATC$$

$$V(J, T) \propto e^{-\frac{T}{\tau}} = e^{-\frac{6-ATC}{\tau}} = e^{\frac{ATC-6}{\tau}}$$

$$SV(v) \equiv V(J, 6 - (ATC + v)) - V(J, 6 - ATC)$$

$$SV(v) \propto e^{-\frac{6-(ATC+v)}{\tau}} - e^{-\frac{6-ATC}{\tau}} = e^{-\frac{6-ATC}{\tau}}\left(e^{\frac{v}{\tau}} - 1\right)$$

Let us define the subjective value, of two alternative gambles of value $v_1, v_2$:

$$SV(v_1, v_2) \equiv V(J, 6 - (ATC + v_1)) - V(J, 6 - (ATC + v_2)),\ v_1, v_2 = [-2, +3].$$

$$SV(v_1, v_2) \propto e^{-\frac{6-ATC-v_1}{\tau}} - e^{-\frac{6-ATC-v_2}{\tau}} = e^{-\frac{6-ATC}{\tau}}\left(e^{\frac{v_1}{\tau}} - e^{\frac{v_2}{\tau}}\right). \tag{4}$$

We can simplify the notation by considering two terms proportional to the above SV.

$$\begin{aligned} SV_1 &= \alpha_1 e^{-6/\tau} e^{ATC/\tau} e^{v_1/\tau}, \\ SV_2 &= \alpha_2 e^{-6/\tau} e^{ATC/\tau} e^{v_2/\tau}, \end{aligned} \tag{5}$$

In our task, values have top and bottom parts $v^t$, $v^b$ with probabilities $p^t, p^b = 1 - p^t$.

$$\begin{aligned} SV_1 &= e^{-6/\tau} e^{ATC/\tau}\left(\alpha_1 \mathbb{E}\left[e^{v_1/\tau}\right] + \alpha_{r_1} R_1\right), \\ SV_2 &= e^{-6/\tau} e^{ATC/\tau}\left(\alpha_2 \mathbb{E}\left[e^{v_2/\tau}\right] + \alpha_{r_2} R_2\right), \end{aligned} \tag{6}$$

We can introduce the notion of expected value and risk:

$$\begin{aligned} \mathbb{E}\left[e^{v/\tau}\right] &= p^t e^{v^t/\tau} + p^b e^{v^b/\tau}, \\ R = VAR\left[e^{v/\tau}\right] &= p^t\left(e^{v^t/\tau} - \mathbb{E}[e^{v/\tau}]\right)^2 + p^b\left(e^{v^b/\tau} - \mathbb{E}[e^{v/\tau}]\right)^2. \end{aligned} \tag{7}$$

In this first version, defined as '**TDM**', the $SV$s are fit to the probability of choosing first offer $p(ch1)$ via the logistic regression

$$\begin{aligned} \text{logit}(p(ch1)) = \beta_0 &+ \beta_1 e^{(ATC-6)/\tau}\left(\mathbb{E}\left[e^{\frac{v_1}{\tau}}\right] - \mathbb{E}\left[e^{\frac{v_2}{\tau}}\right]\right) \\ &+ \beta_2 e^{(ATC-6)/\tau}\left(VAR\left[e^{\frac{v_1}{\tau}}\right] - VAR\left[e^{\frac{v_2}{\tau}}\right]\right), \end{aligned} \tag{8}$$

i.e. by computing $\widehat{\Theta} = [\hat{\tau}, \hat{\beta}_0, \hat{\beta}_1, \hat{\beta}_2]$ via maximum likelihood estimation using train set. Numerically, we minimize negative the log-likelihood:

$$-\log L(\widehat{\Theta}|y_i) = -\Sigma_i[y_i \log \hat{p}_i + (1 - y_i)\log(1 - \hat{p}_i)], \tag{9}$$

using $y_i$ from $i^{th}$ test trial (1 if first option is chosen, 0 otherwise), using predictions

$$\begin{aligned} \hat{p}_i = \text{logit}^{-1}(\hat{\beta}_0 &+ \hat{\beta}_1 e^{(ATC_i-6)/\hat{\tau}}(\mathbb{E}\left[e^{v_{1,i}/\hat{\tau}}\right] - \mathbb{E}\left[e^{v_{2,i}/\hat{\tau}}\right]) \\ &+ \hat{\beta}_2 e^{(ATC_i-6)/\hat{\tau}}(VAR\left[e^{v_{1,i}/\hat{\tau}}\right] - VAR\left[e^{v_{2,i}/\hat{\tau}}\right])). \end{aligned} \tag{10}$$

For stable and interpretable fits, we applied Ridge regularization with $\lambda_{\text{ridge}} = 0.1$ for unbound parameters prone to generate result instability, although we tested that omitting Ridge regression did not qualitatively alter the performances of the model. Specifically, we apply penalization on the temporal discounting constant $(\tau)$ and value sensitivity parameters

$(\beta_1, \beta_2)$. Since the temporal discounting constant is used at the exponent, it can produce large changes in values or lead to regions of the parameter space where the exponentials become too steep or too flat to meaningfully reflect behavioral variation.

Besides choice prediction accuracy, we use the following fit quality metrics:

$$\begin{gathered}
R^2_{adj} = 1 - \left(\frac{\Sigma_i (y_i - \hat{p}_i)^2}{\Sigma_i (y_i - \bar{y}_i)^2}\right) \frac{n_{trials} - 1}{n_{trials} - n_{\widehat{\Theta}} - 1}, \\
-\log L(\widehat{\Theta}|y_i) = -\Sigma_i [y_i \log \hat{p}_i + (1 - y_i) \log(1 - \hat{p}_i)] + \lambda_{\text{ridge}} |\widehat{\Theta}_{\text{ridge}}|^2, \\
AIC = 2n_{\widehat{\Theta}} + 2\left(-\log(L) + \lambda_{\text{ridge}} |\widehat{\Theta}_{\text{ridge}}|^2\right), \\
BIC = \log(n_{\text{trials}})\, n_{\Theta} + 2\left(-\log(L) + \lambda_{\text{ridge}} |\widehat{\Theta}_{\text{ridge}}|^2\right),
\end{gathered} \tag{11}$$

$\widehat{\Theta}_{\text{ridge}} = [\hat{\tau}, \hat{\beta}_1, \hat{\beta}_2]$, $n_{trials}$ is the number of trials used in each run, $n_{\widehat{\Theta}}$ is the total number of $\widehat{\Theta}$ parameter estimates. Train and test sets are defined via $k = 4$ fold cross-validation.

#### *iv.* Time Discounting and Reference-dependent value (TDM_RDV)

We introduce the notion of reference-dependent value using a '**utility function**' in similar way as in Methods 2.4.3:

$$u(v, ATC) = \begin{cases} (v - r(ATC))^{\gamma(ATC)} & \text{if } v \geq r(ATC) \text{ (gains)} \\ -\lambda(ATC)(r(ATC) - v)^{\gamma(ATC)} & \text{if } v < r(ATC) \text{ (losses)} \end{cases}. \tag{12}$$

We introduce the token-dependent reference as:

$$r(ATC) = \frac{6 - ATC}{1 + e^{-\kappa_0 (ATC - \kappa_1)}}. \tag{13}$$

This function defines a reference value $r(ATC) \to 6 - ATC$ for $ATC \geq \kappa_1$, and a reference value $r(ATC) \to 0$ for $ATC < \kappa_1$. In our data, $\kappa_1$ is numerically fit and on average assumes value $> 3$, meaning that value is referenced to $6 - ATC$ when the accumulated tokens count allows reaching jackpot ($ATC + v$ tokens may reach 6 for $ATC \geq 3$, $v \in [-2, +3]$), and it is compared to zero when accumulated tokens do not allow reaching jackpot ($ATC < 3$). This is equivalent to consider offered value as possible "gain" or "loss" respectively if higher or lower than the "missing tokens for jackpot" threshold $(6 - ATC)$ in trials when jackpot is achievable, and as possible "gain" or "loss" respectively if positive or negative, when jackpot is not possibly achieved.

The utility function $u(v, ATC)$ shapes value $v$ via $\gamma(ATC), \lambda(ATC)$:

$$\gamma(ATC) = \gamma_0 + \gamma_1 ATC, \tag{14}$$

$$\lambda(ATC) = \lambda_0 + \lambda_1 ATC + \lambda_2 ATC^2. \tag{15}$$

The function $\gamma(ATC)$ models the curvature of reference-dependent value assuming it increases with $ATC$, in line with suggestions by our previous analyses (Fig. 4C, Supp. Fig.

S1C). The shaping of the increase is defined for "gains" (related to risk-seeking), and is scaled by $\lambda(ATC)$ for "losses" (related to loss-aversion), assuming asymmetry by which losses "hurt more than gains", inspired by Prospect Theory. For losses, we use a quadratic polynomial fit following evidence for an inverted-u shape relationship between risk and $ATC$ (Fig. 4D, Supp. Fig. S1D).

The '**TDM_RDV**' model combines RDV and TDM by fitting choices via:

$$\begin{aligned}\text{logit}\big(p(ch1)\big) &= \beta_0 + \beta_1 e^{(ATC-6)/\tau}\left(\mathbb{E}\left[e^{\frac{u(v_1,ATC)}{\tau}}\right] - \mathbb{E}\left[e^{\frac{u(v_2,ATC)}{\tau}}\right]\right) \\ &+ \beta_2 e^{(ATC-6)/\tau}\left(VAR\left[e^{\frac{u(v_1,ATC)}{\tau}}\right] - VAR\left[e^{\frac{u(v_2,ATC)}{\tau}}\right]\right),\end{aligned} \tag{16}$$

using training set data to compute estimates $\widehat{\Theta} = [\hat{\tau}, \hat{\gamma}_0, \hat{\gamma}_1, \hat{\lambda}_0, \hat{\lambda}_1, \hat{\lambda}_2, \hat{\beta}_0, \hat{\beta}_1, \hat{\beta}_2]$ and maximize likelihood between test data $y_i$ and the predicted variable $\hat{p}_i$ as in TDM. Also in this case, we apply Ridge penalty to $\tau, \beta_1, \beta_2$, and $k = 4$ fold cross-validation.

#### *v.* Putting side to side TDM RDV and linear RDV (TDM_RDV_and_linear_RDV)

This model combines linear RDV (*ii.*) and TDM RDV (*iv.*) by fitting choices via:

$$\begin{aligned}\text{logit}\big(p(ch1)\big) &= \beta_0 + \beta_1 e^{(ATC-6)/\tau}\left(\mathbb{E}\left[e^{\frac{u(v_1,ATC)}{\tau}}\right] - \mathbb{E}\left[e^{\frac{u(v_2,ATC)}{\tau}}\right]\right) \\ &+ \beta_2 e^{(ATC-6)/\tau}\left(VAR\left[e^{\frac{u(v_1,ATC)}{\tau}}\right] - VAR\left[e^{\frac{u(v_2,ATC)}{\tau}}\right]\right) \\ &+ \beta_3(\mathbb{E}[u(v_1,ATC)] - \mathbb{E}[u(v_2,ATC)]) \\ &+ \beta_4(VAR[u(v_1,ATC)] - VAR[u(v_2,ATC)]).\end{aligned} \tag{17}$$

Thus defining:

$$\begin{aligned}SV_i &= \beta_1 e^{(ATC-6)/\tau}\mathbb{E}\left[e^{\frac{u(v_i,ATC)}{\tau}}\right] + \beta_2 e^{(ATC-6)/\tau} VAR\left[e^{\frac{u(v_i,ATC)}{\tau}}\right] \\ &+ \beta_3\mathbb{E}[u(v_i,ATC)] + \beta_4 VAR[u(v_i,ATC)]\end{aligned} \qquad i = 1,2. \tag{18}$$

We use training set data to compute estimates $\widehat{\Theta} = [\hat{\tau}, \hat{\gamma}_0, \hat{\gamma}_1, \hat{\lambda}_0, \hat{\lambda}_1, \hat{\lambda}_2, \hat{\beta}_0, \hat{\beta}_1, \hat{\beta}_2, , \hat{\beta}_3, \hat{\beta}_4]$ and maximize likelihood between test data $y_i$ and the predicted variable $\hat{p}_i$ as in TDM_RDV. We apply Ridge penalty to $\tau, \beta_1, \beta_2, \beta_3, \beta_4$, and $k = 4$ fold cross-validation.

### Model Comparisons: Results

All these explorations lead us to conclude that the best model is the reference-dependent model (*ii.*) 'linear_RDV'. Applying an asymmetric shaping to offer values yields the best performances in terms of choice prediction Accuracy, cross-validated $R^2_{adj}$, negative log likelihood, AIC, BIC in both subjects (Supp. Fig. S21, Supp. Table ST14).

The introduction of an exponential time-discounting model 'TDM' (*iii.*) leads to modest change in accuracy compared to the 'linear' (*i.*) model (Supp. Fig. S21, Supp. Table ST14).

This model entails extra complexity (fitting an exponential denominator term) with respect to linear models, and shows worse likelihood, AIC and BIC, and reduced generalization properties shown by lower cross-validated $R^2_{adj}$. Applying the RDV utility function (Methods 2.4.3) to time-discounted values as in 'TDM_RDV' (*iv.*) only shows modest changes in negative log likelihood, Accuracy, $R^2_{adj}$, AIC and BIC (Supp. Fig. S21, Supp. Table ST14), at the cost of introducing extra complexity.

Overall, we find that applying a linear reference-dependent utility to offer values as in 'linear_RDV' (*ii.*) is the best model design, as it shows significant best performance compared to all other models, including 'TDM' and 'TDM_RDV' in both subjects, in terms of Accuracy, $R^2_{adj}$, likelihood, AIC, BIC (Supp. Fig. S21, Supp. Table ST14).

Lastly, we directly compared the TDM_RDV and linear_RDV regressors within the combined model 'TDM_RDV_and_linear_RDV'. Notably, the regression weights associated with the linear_RDV terms remained significant and of comparable magnitude in both linear_RDV and TDM_RDV_and_linear_RDV (Supp. Table ST15). This indicates that reference dependency plays a central role in shaping choice behavior and that the inclusion of temporal-discounting terms provides no substantial improvement in model performance, consistent with prior work examining discounting at similar timescales (Blanchard et al., 2013; Hayden, 2016). The combined model achieves accuracy comparable to linear_RDV, whereas linear_RDV exhibits superior adjusted $R^2$, likelihood, AIC, and BIC (Supp. Fig. S21) thanks to its reduced complexity. Taken together, these comparisons across a broad set of alternative models demonstrate that linear_RDV provides the most parsimonious and best-performing account of choice data, and we therefore rely on this model for our results and interpretations. However, we now also exhaustively report results from the combined model in Supp. Table ST14-ST15, to show that if we include TDM_RDV next to linear_RDV regressors, we find that TDM_RDV terms will show significant weights, albeit weaker than linear_RDV ones. We have also replicated results from neural analyses using SV definition from the combined model TDM_RDV_and_linear_RDV, reported in Supp. Fig. S22.

**References**

Blanchard, T. C., Pearson, J. M., & Hayden, B. Y. (2013). Postreward delays and systematic biases in measures of animal temporal discounting. Proceedings of the National Academy of Sciences, 110(38), 15491-15496.

Hayden, B. Y. (2016). Time discounting and time preference in animals: A critical review. Psychonomic bulletin & review, 23(1), 39-53.
